# Basal protrusions mediate spatiotemporal patterns of spinal neuron differentiation

**DOI:** 10.1101/559195

**Authors:** Zena Hadjivasiliou, Rachel Moore, Rebecca McIntosh, Gabriel Galea, Jon Clarke, Paula Alexandre

## Abstract

During early spinal cord development, neurons of particular subtypes differentiate with a sparse periodic pattern while later neurons differentiate in the intervening space to eventually produce continuous columns of similar neurons. The mechanisms that regulate this spatiotemporal pattern are unknown. *In vivo* imaging of zebrafish reveals differentiating spinal neurons transiently extend two long protrusions along the basal surface of the spinal cord prior to axon initiation. These protrusions express Delta protein consistent with the possibility they influence Notch signalling at a distance of several cell diameters. Experimental reduction of laminin expression leads to smaller protrusions and shorter distances between differentiating neurons. The experimental data and a theoretical model support the proposal that the pattern of neuronal differentiation is regulated by transient basal protrusions that deliver temporally controlled lateral inhibition mediated at a distance. This work uncovers novel, stereotyped protrusive activity of new-born neurons that organizes long distance spatiotemporal patterning of differentiation.

## Introduction

During the early stages of vertebrate neurogenesis, neurons of particular subtypes initially differentiate along the spinal cord with a sparse periodic pattern but eventually produce more continuous columns of similar neurons (Figure 1A, Dale et al. 1987, Roberts et al. 1987, Higashijima et al 2004a, Higashijima et al 2004b, Kimura et al 2006, Batista et al 2008, England et al 2011). The mechanisms that regulate this pattern of differentiation are unknown. Delta-Notch mediated lateral inhibition is a regulator of vertebrate neurogenesis (for example Chitnis et al 1995, Henrique et al 1997, Appel et al 2001, Okigawa et al 2014) but this conventionally operates in a juxtacrine fashion between Delta expressing cells and their immediate neighbours and cannot explain the spatial and temporal pattern of neuronal differentiation along the embryo spinal cord.

**Figure 1.**
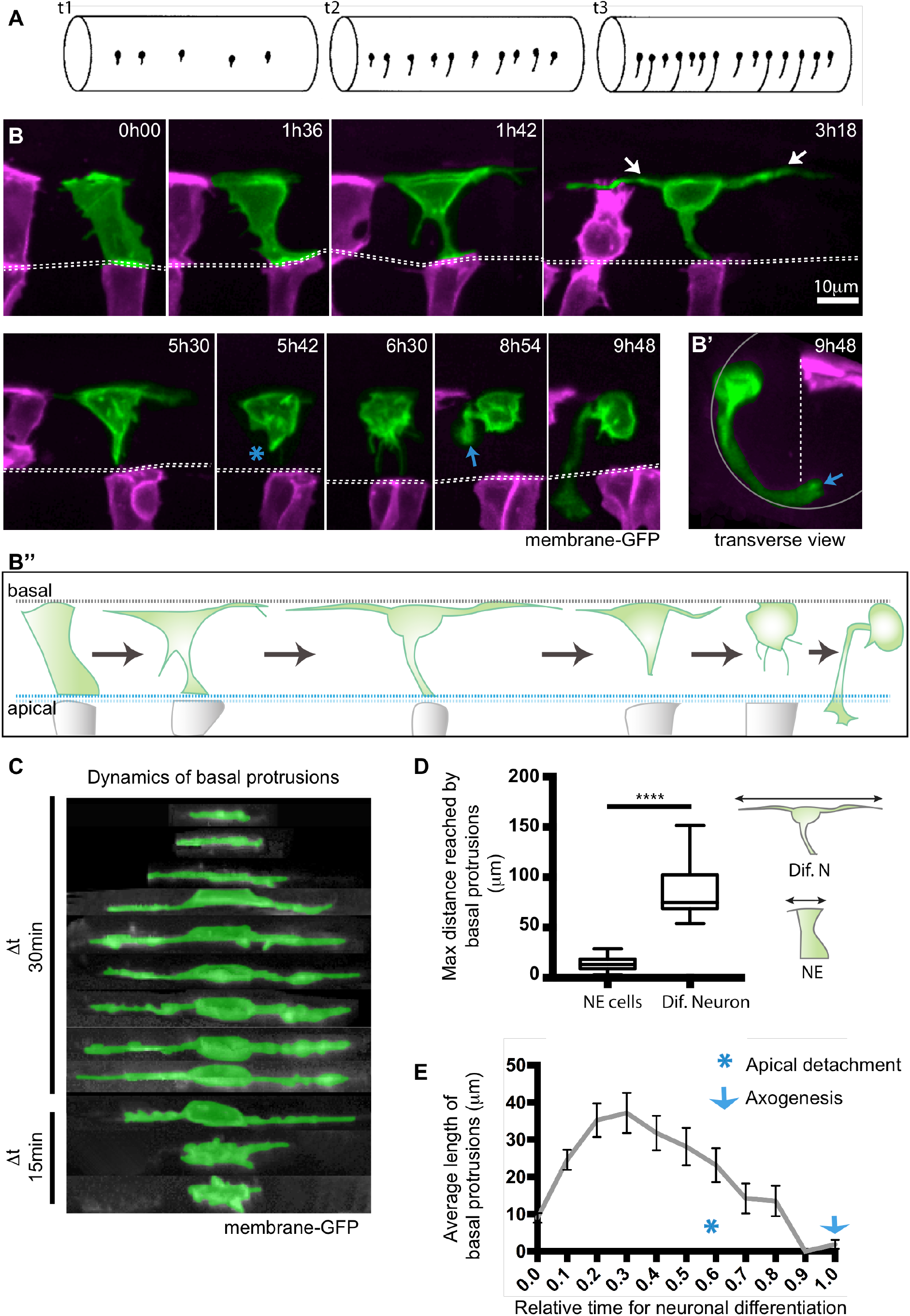
Differentiating spinal neurons transiently elongate two long basal protrusions along the A/P axis before detaching from the apical surface. A) Diagram to show spinal neurons differentiate with an initial long distance spacing pattern (t1). Later differentiating neurons of the same type subsequently fill in the gaps between the earlier differentiated cells (t2 and t3) to generate a near continuous column of neurons. Lateral view of spinal cord, dorsal to top. B) Image sequence from confocal time-lapse from dorsal view illustrates the early steps in neuronal differentiation that precede axogenesis in the spinal cord. A differentiating neuron (green) transiently adopts a T-shape through the maintenance of an apical attachment and the elongation of two long cellular protrusions at the basal surface of the neuroepithelium (arrowed in timepoint 3h18). Following the retraction of basal protrusions, the apical process detaches (blue asterisk in timepoint 5h42). The axon is formed (blue arrow in 8h54) and grows ventrally and across the midline (see Movie 1). Images are maximum projections from confocal z-stacks. B’) shows a transverse reconstruction of B) at 9h48. Cells visualized with membrane-GFP, with non-neuronal cells artificially coloured in magenta. Dashed line shows position of the apical surfaces. B’’) Diagram summarises the steps involved in neuronal differentiation: transient formation of basal protrusions, followed by their retraction, apical detachment and axonal growth. Apical and basal surfaces of the neuroepithelium are outlined by a blue (bottom) and grey dashed lines (top), respectively. C) Kymographic representation of extension and retraction of basal protrusions of a differentiating neuron (green) viewed laterally. D) Box-and-whisker plot showing maximal basal extension of differentiating neurons (mean ± SD, 86.8 ± 25.3 μm, n=21 cells) and non-differentiating neuroepithelial cells (mean ± SD, 14.3 ± 6.2 μm, n=74 cells). The line inside the box represents the median and whiskers represent minimum and maximum values. Data analysed using Kruskal-Wallis with Dunn’s multiple comparison test (p<0.0001). E) Average length of individual basal protrusions during neuronal differentiation (n=13 cells). The time has been normalised from (0), the moment in which differentiating neurons begin elongation of basal protrusions, to (1), when neurons initiate axon formation.

Recent evidence however suggests the distance over which contact mediated signaling of various types can operate can be extended by cellular protrusions capable of spanning several cell diameters (reviewed in Buszczak et al, 2016, Prols et al, 2016). For example, signaling through long cellular protrusions plays a role during limb patterning in the chick embryo (Sanders et al, 2013), in the development of the zebrafish pigmentation stripes (Eom et al, 2015) as well as neural plate patterning also in the zebrafish (Stanganello et al 2015). In fact, dynamic cellular protrusions from the basal surface of sensory organ precursor (SOP) cells have been proposed to mediate long-distance lateral inhibition to regulate the sparse distribution of mechanosensory bristles in the fly notum and wing disk (De Joussineau et al. 2003; Cohen et al. 2010; Hadjivasiliou et al 2016, Hunter et al 2016, 2018). Whether similar protrusive activity mediates long-distance spacing patterns in the vertebrate Central Nervous System (CNS) is not known, but long and short cellular protrusions expressing the Notch ligand Delta-like 1 have been described on intermediate progenitors in the embryonic mammalian cortex (Nelson et al, 2013). Furthermore, dynamic protrusive activity on the surface of recently born spinal neurons can be observed in slice cultures of chick embryo spinal cord (Das and Storey 2014).

To determine whether cellular protrusions could also play a role in the patterning of spinal neuronal differentiation we addressed these issues in the zebrafish embryo spinal cord. Live *in vivo* imaging revealed all spinal neurons transiently extend two long cellular protrusions along the basal surface of the spinal cord prior to axon initiation and apical detachment. We show these long basal protrusions express Delta protein at high level and Notch reporter activation is upregulated in cells in their vicinity. Furthermore, experimental reduction of the basal protrusion length results in reduced spacing between differentiating neurons. Our *in vivo* data is supported by a theoretical model, whose output is consistent with the proposal that neuronal differentiation is regulated by lateral inhibition mediated at a distance by transient basal protrusions. Our work thus reveals novel, stereotyped protrusive activity of differentiating neurons that organizes long distance spatiotemporal patterning of neuronal differentiation in the embryo spinal cord.

## Results

### Differentiating spinal neurons transiently elongate two long basal protrusions along the A/P axis before detaching from the apical surface

To study the early phases of neuronal differentiation *in vivo*, we have labelled small numbers of cells in the zebrafish embryo spinal cord by mosaic expression of membrane GFP and captured their behavior with confocal time-lapse microscopy from 18 to 42 hours post fertilization (hpf). Analysis of more than 100 cells that differentiate into neurons reveals a stereotyped, transient T-shaped transition from a cell that is attached to the apical surface of the neuroepithelium to a basally positioned neuron with the beginnings of a single axon extension. This transition involves the elongation of two longitudinally directed cellular processes that protrude along the basal surface of the neural tube, one protruding anteriorly and the other posteriorly (Figure 1B - time point 1h42 and 3h18; Figure 1B’’, C and Movie 1). These basal protrusions can be asymmetric in length (in 17/28 cells) and each protrusion can reach up to 109 μm (mean ± SD, 42.6 ± 20.2 μm, n=24 cells) with a combined length of up to 151.5 μm (mean ± SD, 86.8 ± 25.3 μm, n=21 cells) (Figure 1D, Supplementary Figure 1). The basal protrusions are typically present on differentiating neurons for several hours (mean ± SD, 6.8 ± 2.2 h, n=13 cells) and grow on average 6x longer than the basal extensions formed by the non-differentiating neural progenitors (mean ± SD, 14.3 ± 6.2 μm, n=74 cells) (Figure 1D). After reaching their maximum length, basal protrusions begin to retract back to the cell body, and this is accompanied by the detachment and retraction of the apical process (19/24 cells) (Figure 1B - from time point 3h18 to 5h42, Figure 1E). After these three processes have retracted, cells adopt a near spherical shape and the cell body becomes highly enriched in filopodial activity that diminishes prior to axon formation (23/27) (Figure 1B-timepoint 6h30 and 8h54, B’, Movie 1). The transient basal protrusions contain dynamic microtubules (Supplementary Figure 2A, Movie 2) and often produce filopodia that are directed radially towards the apical surface (Supplementary Figure 2B, Movie 3). Basal protrusions from nearby differentiating cells can overlap (Supplementary Figure 2C).

**Figure 2.**
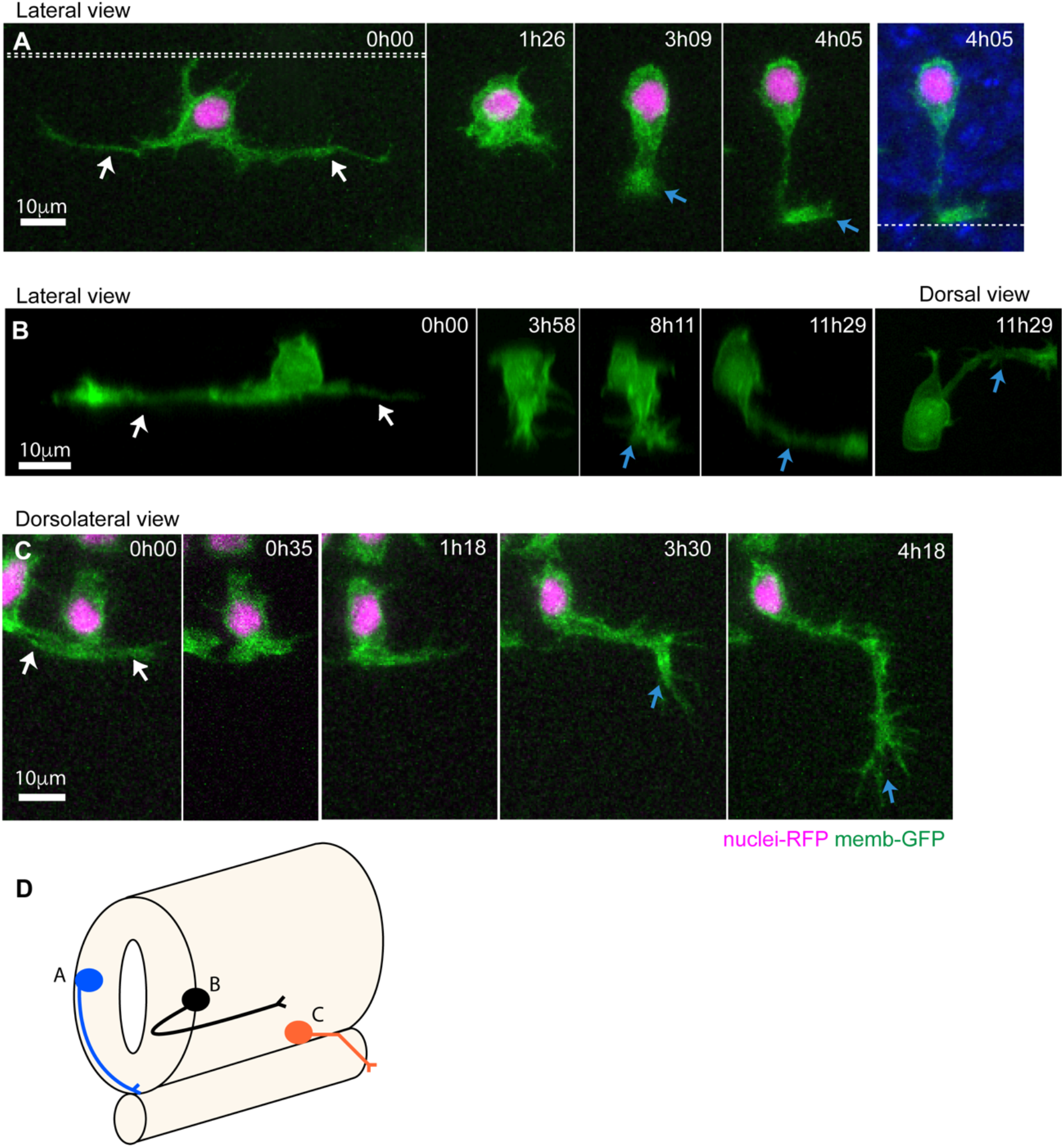
Stereotyped axon formation follows basal protrusion retraction. A) Image sequence from a time-lapse showing a neuron with long basal protrusions (white arrows) that are fully retracted before axon initiation (blue arrow at time 3h09). The axon grows circumferentially and crosses the ventral floor plate (blue arrow at time 4h05) (Movie 4). Double dashed line shows the apical surfaces. Single dashed line is the ventral surface of the spinal cord. B) Image sequence from a time-lapse shows a neuron with long basal protrusions (white arrows) that are fully retracted before axon initiation (blue arrow at time 8h11). The axon is initiated from the ventral surface of the neuron and then grows longitudinally and ipsilaterally along the spinal cord (blue arrow at time 11h29). C) Image sequence from a time-lapse of a motor neuron with short basal protrusions (white arrows) that are retracted by timepoint 0h35. The exact point of axon extension is not clear, but the axon (blue arrow) changes direction to leave ventral spinal cord and grow into muscle at time 3h30.D) Summary diagram of neuron morphologies shown in A) to C). Neurons were labelled with membrane-GFP (green) and H2B-RFP to show nuclei in A and C. All images are projected images from confocal z-stacks.

Differentiating spinal neurons thus stereotypically adopt a transient T-shape prior to apical detachment and axon formation (summarized in Figure 1B”). These observations reveal a new *in vivo* cellular behaviour that precedes axogenesis and distinguishes the neuronal precursors in the process of differentiation from surrounding neural progenitors.

### Stereotyped axon formation follows basal protrusion retraction

Studies of neuronal differentiation *in vitro* have revealed that axons derive by selection and specialization of one neurite from several pre-existing neurites (Dotti et al. 1988, Craig & Banker et al 1994, Barnes et al, 2009). To investigate whether the axons of spinal neurons *in vivo* might derive from the transient long basal protrusions, we monitored axon initiation. Neurons were located at many different dorsoventral (D/V) levels of spinal cord and thus likely represent many different subtypes of spinal projection neuron. Our 3D reconstruction analyses revealed that axonal outgrowth almost always follows the full retraction of basal protrusions (27/31 cells) (Figures 2A, B, Movies 1, 4) and, in contrast to *in vitro* observations, axons never differentiated from an existing cellular protrusion. The majority of subtypes of spinal neurons have an axon that runs ventrally and circumferentially from the cell body before either crossing the ventral floor plate or turning anteriorly or posteriorly to join the ipsilateral longitudinal axon tracts (Bernhardt et al 1990). Our observations show this ventral circumferential axon trajectory is initiated stereotypically at the outset of axon growth, directly from the cell body and is spatially independent of and perpendicular to the preceding transient basal protrusions (Figures 1B, 2A, B, Movies 1, 4). In only one case have we seen a neuron generate what appears to be a forked axon with two ventrally directed branches. In this case, one of these branches was quickly retracted leaving the usual morphology of a single ventral axon.

Our analysis does not include the primary sensory Rohon-Beard neurons which develop three axons (two central longitudinal axons and a peripheral axon) and are likely to use a different programme of axogenesis (Andersen and Halloran 2012). Our data also contains only one definitive motoneuron because their very ventral location impedes imaging. However, the single motoneuron has short basal protrusions and was the only neuron that did not have a ventral trajectory to its initial axon growth, instead it directed its axon laterally from the cell body towards the nearby somite boundary before exiting the cord to innervate the muscles (Figure 2C, observations summarized in diagram in Figure 2D).

### Non-apical progenitors in spinal cord also extend basal protrusions prior to apical detachment

In addition to the apical progenitors that generate most of the neurons of zebrafish CNS, a scarce population of basal progenitors that divide in non-apical locations is also present (Alexandre et al, 2010, McIntosh et al, 2017). We call these progenitors, Non-Apical Progenitors (or NAPs) and previously demonstrated that the majority of spinal NAPs express Vsx1 and share molecular and regulatory mechanisms with neurons (McIntosh et al, 2017). This prompted us to investigate whether spinal NAPs might also share the morphological programme of differentiation with neurons. We were able to monitor 7 NAPs by confocal time-lapse microscopy all of which undergo the stereotypical T-shape transition characteristic of differentiating neurons prior to their basal mitosis (Figure 3A and Movie 5). The NAP exemplified in Figure 3A has a basal cell body that transiently extends a pair of long basal protrusions that are filopodia rich while still attached to the apical surface (Figure 3A, Movie 5). The basal protrusions on NAPs are often asymmetric in length (6/7 cases). On some cells, basal protrusions do not fully retract before NAP mitoses (4/7 cells) (Figure 3A, B, Movie 5). In these cases, the retraction of basal protrusions is completed after mitosis (green arrow in Figure 3A, B) but still prior to axon formation in the two daughter neurons (blue arrow in Figure 3A, B, Movie 5).

**Figure 3.**
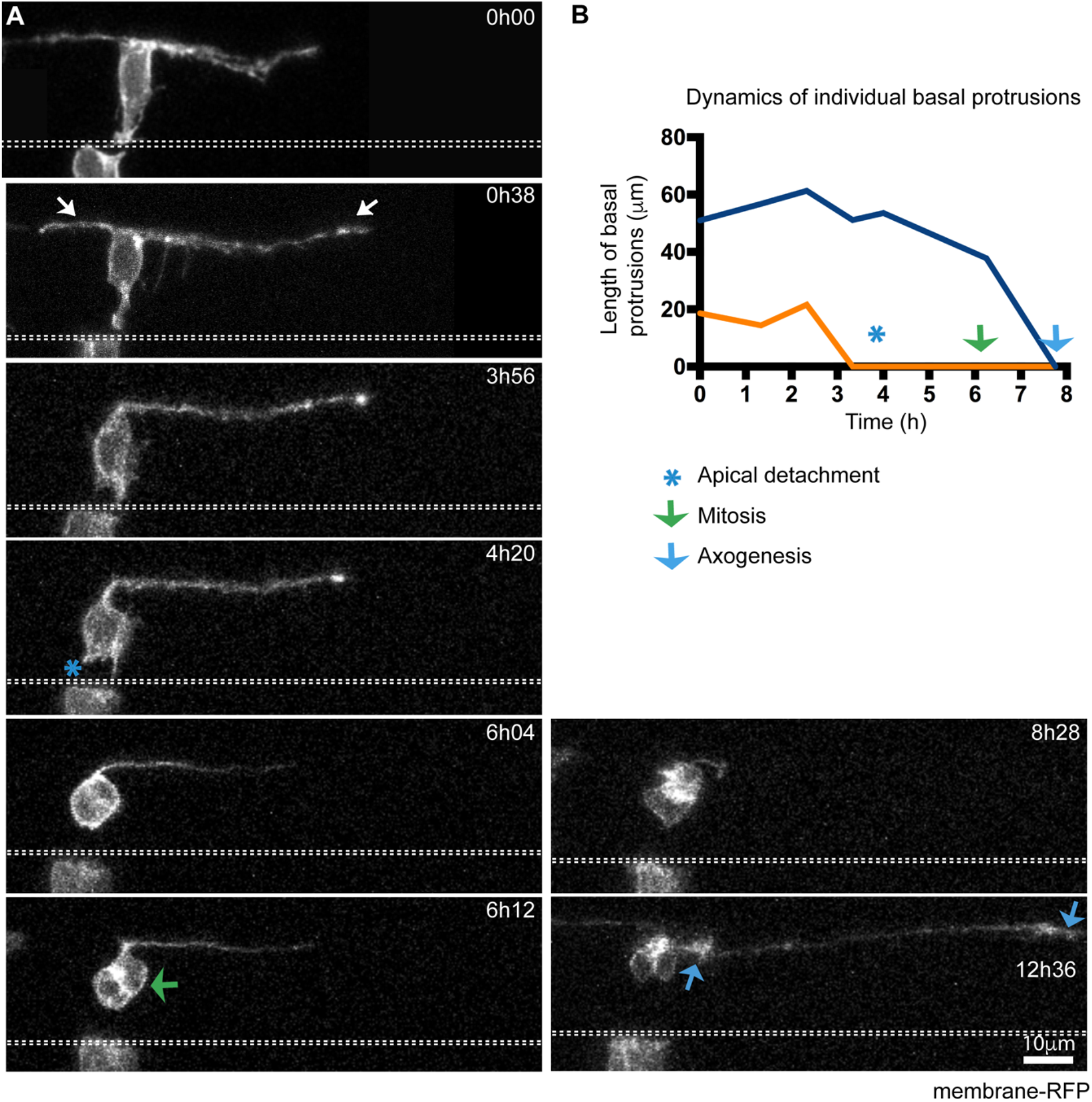
Non-apical progenitors in spinal cord also extend basal protrusions prior to apical detachment. A) Image sequence showing a non-apical progenitor (NAP) with elongated basal protrusions (white arrows). The NAP retracts the apical attachment (blue asterisk in timepoint 4h20) before basal protrusions fully retract. Following apical detachment, the cell body rounds up away from apical surface of the neuroepithelium and undergoes mitosis (green arrow at time-point 6h12). The NAP is neurogenically committed and produces two neuron daughters, each initiating axon growth at different time-points (blue arrows indicate two growth cones at time 12h36) (Movie 5). The apical surface is outlined by white dashed line. View is dorsal. All images are projected images from confocal z-stacks.B) Graph showing the changes in length over time of the two basal protrusions from the NAP shown in A). Timepoints of when apical detachment, mitosis and first axon elongation take place are also indicated.

These observations show spinal neurons and NAPs share common stereotypical morphological behaviours, and further confirm that spinal Vsx1 NAPs and differentiating neurons share cellular and molecular characteristics as suggested previously (McIntosh et al, 2017).

### Differentiating telencephalic neurons do not form long transient basal protrusions

The elongation of basal protrusions seems to be a consistent feature of differentiating neurons and non-apical progenitors in the zebrafish spinal cord. To investigate whether the T-shape transition is common to differentiating neurons in other regions of the zebrafish CNS we analysed neuronal differentiation in the dorsal telencephalon from 20 to 40hpf. Using this approach, we find that differentiating neurons in the telencephalon do not extend transient basal protrusions prior to apical detachment and axogenesis (Figure 4) (n=16/16 cells) (Movie 6). In these cells, axon formation derives from the basal end of the new neuron’s radial process and usually immediately follows the detachment of the neuron from the apical surface (Figure 4 and Movie 6).

**Figure 4.**
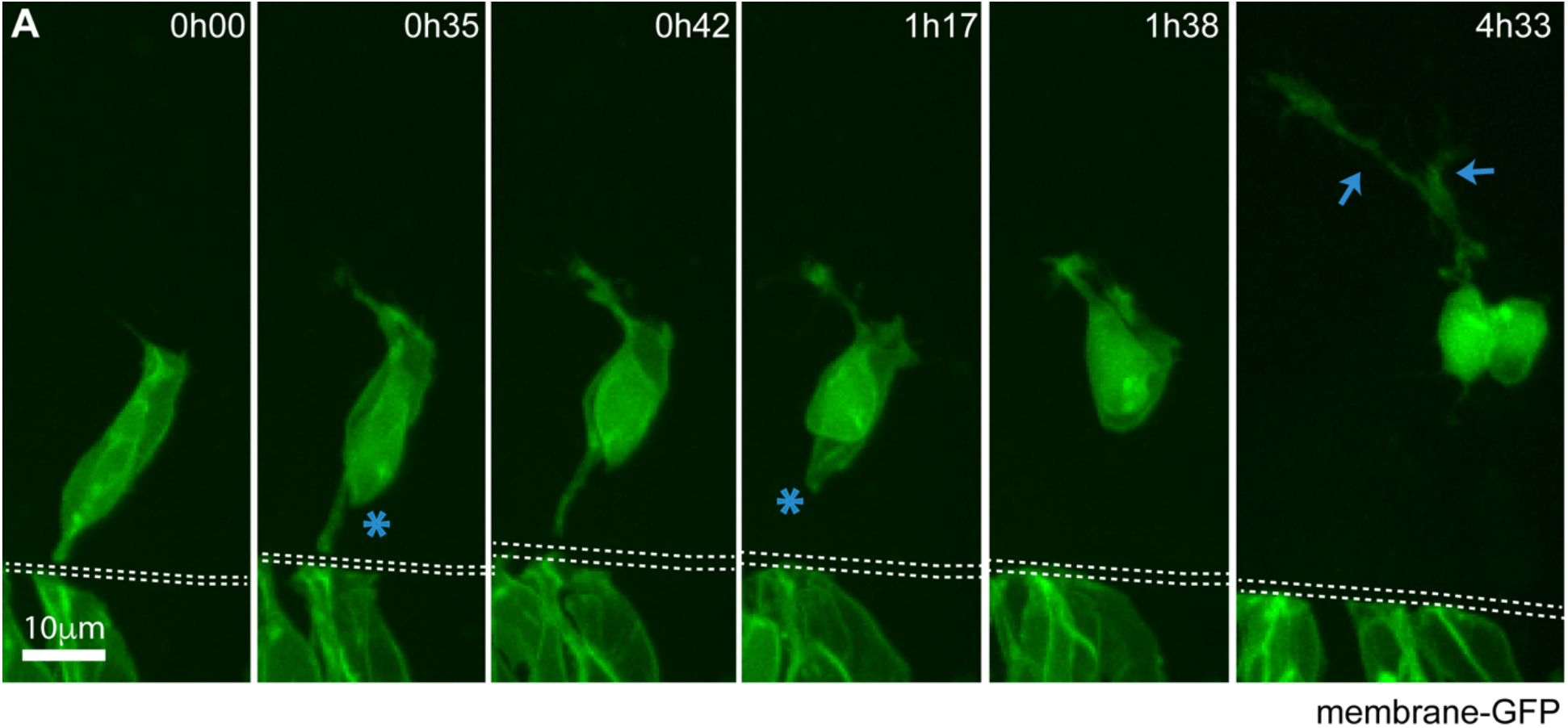
Differentiating telencephalic neurons do not form long transient basal protrusions. A) Image sequence from time-lapse showing a pair of differentiating telencephalic neurons. Long basal protrusions are not observed. Short basal protrusions from time point 0h35 on are the initial growth of axons. The neurons detach from the apical surface at 0h35 and 1h17 (blue asterisks). Extending axons are visible at 4h33 (blue arrows) (Movie 6). Dashed lines show apical surfaces. Images are projections of confocal z-stacks. View is dorsal.

These observations demonstrate the programmes of axogenesis and apical release are regionally distinct, suggesting a region-specific role for the T-shape transition in spinal differentiation.

### Neurons rarely differentiate close together in time and space

To quantify the spatiotemporal dynamics of spinal neuron differentiation we used *in vivo* confocal microscopy to determine the spatiotemporal pattern of differentiation of Vsx1:GFP expressing neurons in the zebrafish spinal cord. Vsx1:GFP neurons are born in pairs from the terminal division of *vsx1*-expressing NAPs (Kimura et al 2008, McIntosh et al 2017). GFP is detected in their progenitor immediately before terminal division and maintained in their daughters (Figure 5A). The appearance of adjacent GFP expressing daughters thus offers a distinct and easily recognised time point to record as the start of differentiation of those neurons (Figure 5A). Using this criterion we recorded the position and time of the start of differentiation of every pair of Vsx1 positive neurons in a 250 to 400μm length of spinal cord at the level of somites 9 to 14 and between 19 and 27 hpf. We did this for both left and right sides in 17 embryos, thus recording 449 Vsx1 differentiation events in space and time within 34 equivalent stretches of spinal cord (Figure 5B and Supplementary Figure 5A).

**Figure 5.**
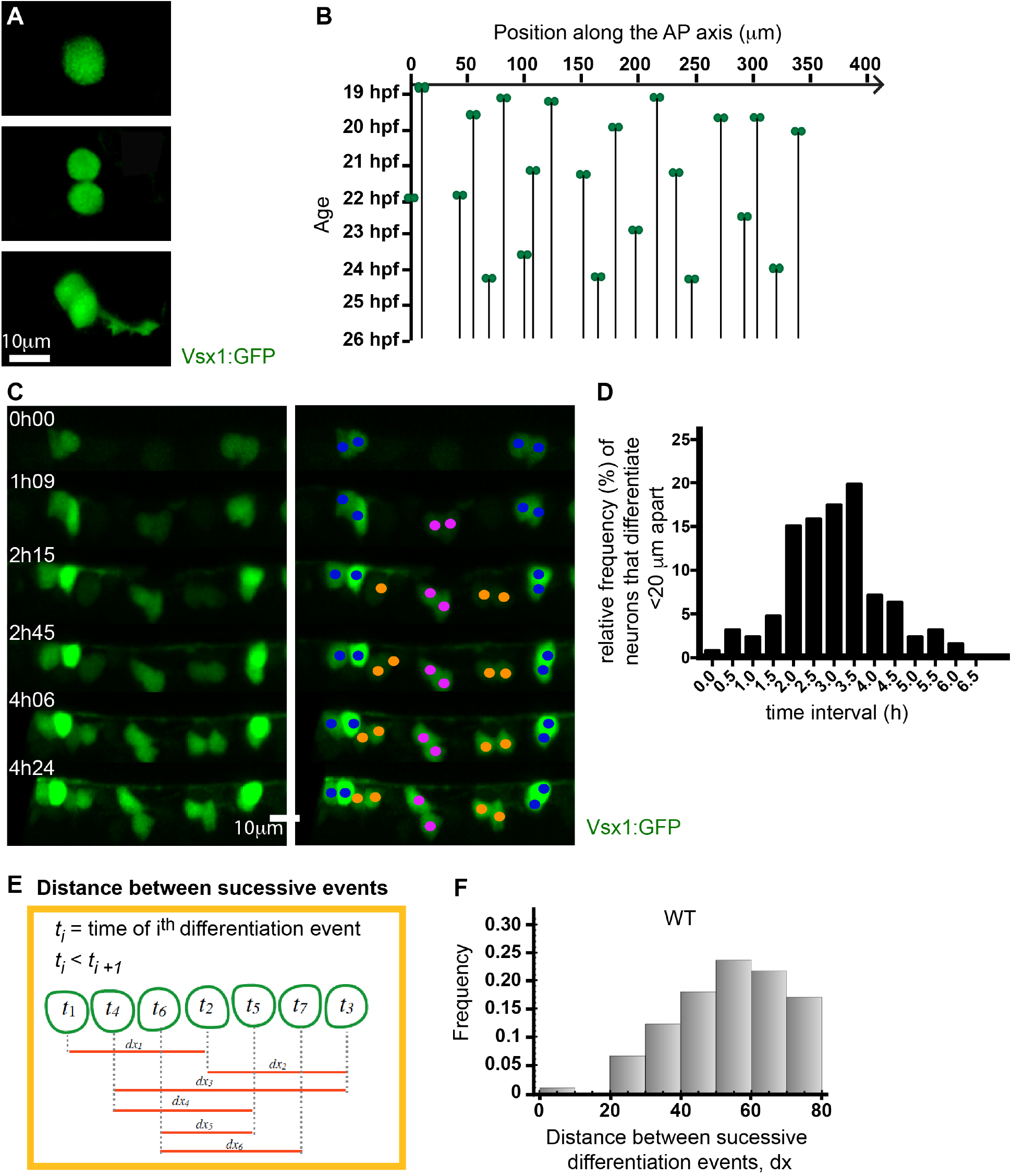
Neurons rarely differentiate close together in time and space. A) Vsx1:GFP expression in a single cell before, during and after a NAP division. Following mitosis, the GFP expression is maintained and axogenesis can be followed in both daughter neurons. B) Spatiotemporal pattern of Vsx1:GFP neuronal precursor differentiation from 19 to 27 hpf. The location of Vsx1:GFP NAPs at the time of mitosis are represented as pairs of green circles and plotted in space (X-axis) and time (Y-axis). The black lines descending through time from the pair of green circles represent the position held by the daughter cells after mitosis. C) Image sequence from a time-lapse showing the differentiation of Vsx1:GFP neurons in one section of spinal cord through time. The left panel shows Vsx1:GFP neurons differentiating over time. In the right panel, cells have been colour-coded to denote sister pairs. All images are projections from small confocal z-stacks. (See also Movie 7.) D) Frequency distribution showing the difference in time between Vsx1:GFP mitoses that occur less than 20μm apart. E) Diagram illustrating the method used to calculate the distance between successive Vsx1:GFP differentiation events from a time lapse movie. *t* gives time of differentiation and *dx* the distance between successive differentiation events. F) Histogram showing the distribution of the distance between successive Vsx1:GFP differentiation events in WT embryos.

This data confirms that Vsx1 neurons differentiate in a long distance spacing pattern with later born neurons differentiating in the gaps between already existing neurons (Figure 5C Movie 7). Time-lapse movies show no evidence that Vsx1 neurons or their progenitors migrate into this space, rather these cells maintain stable positions. This pattern of sequential differentiation in the gaps continues for the next 6 hours at which time a near continuous line of Vsx1 neurons has been generated (Figure 5B, C, Movie 7 and Supplementary Figure 5A).

To quantify this spatiotemporal pattern of differentiation we looked at the timing of Vsx1 differentiation events that happened less than 20μm apart. Neuroepithelial cells are typically 10.5 ± 4.1μm (mean ± SD, n= 95 cells) wide at their basal pole, so this correlates to less than two cell diameters. Of the 449 Vsx1:GFP differentiation events, in only 7 cases (1.6%) were the differentiation events closer in time and space than 20μm and 60 min apart (Figure 5D). The majority (68.3%) of events that occurred within 20μm occurred between 2 and 3.5 hours apart. Additionally, most consecutive Vsx1 differentiation events (i.e. those that occur closest in time) occur at a distance of 50-60μm (Figure 5E, F).

This data suggests the operation of a mechanism that regulates the spatiotemporal differentiation of Vsx1 neurons in order to sequentially transform a long-distance spacing pattern into a continuous column of neurons.

### Transient basal protrusions express DeltaD and Notch activity is upregulated in their vicinity

Our previous section analysed Vsx1 neurons to show that neuronal differentiation in the embryonic zebrafish spinal cord occurs with an initial sparse pattern followed by sequential infilling (Figure 5). Similar patterns of differentiation are also apparent in previous studies of other neuronal subtypes (Gribble et al, 2009, Hutchinson et al, 2006, 2007, Kimura et al, 2008, England et al, 2009). This data suggests a mechanism may exist to transiently inhibit neuronal differentiation over a distance of several cell diameters from each newly differentiating cell, and that this mechanism is sequentially released to allow differentiation in the initially inhibited space. We hypothesise that the transient basal protrusions on newly differentiating neurons and non-apical progenitors could mediate this long distance inhibition in time and space. Since Delta-Notch signalling has been suggested to mediate lateral inhibition at a distance to regulate sparse pattern formation in other systems (reviewed in Prols et al, 2016), we tested whether the transient basal protrusions on differentiating neurons could potentially mediate transient Delta-Notch signalling in our system.

Using an antibody against the DeltaD protein and a DeltaD transgenic reporter line Tg(*DeltaD:GAL4c;UAS:GFP*) (Scheer et al 2001), we were able to determine that the *DeltaD* transgene highlights cells with typical T-shape morphology and that DeltaD protein is specifically enriched in the basal protrusions and cell body of these cells (Figure 6A, A’). Furthermore, if the basal protrusions participate in long-range lateral inhibition we expect them to activate Notch signalling pathway in the surrounding cells contacted by the basal protrusions, importantly this should occur in cells out of range of contact from the neurons cell body. To test whether this is the case, we randomly labelled differentiating neurons in the Notch reporter line Tg(TP1:VenusPEST) (Ninov et al 2012) and monitored the dynamics of Notch activation in nearby cells. We measured the relative mean intensity values of VenusPEST expression in a neuroepithelial region contacted by the labelled basal protrusion (but not the neuronal cell body) and compared it to a control region that had not been contacted by an identified protrusion (Figure 6B). We assessed VenusPEST expression 2h after basal protrusions reached their maximum length. We found the amount of VenusPEST expression is significantly increased in regions spatially related to the identified protrusions when compared to the control region (Figure 6C). These observations are therefore consistent with the hypothesis that basal protrusions activate Notch signaling in order to delay neuronal differentiation in cells at a distance from the differentiating neuronal body.

**Figure 6.**
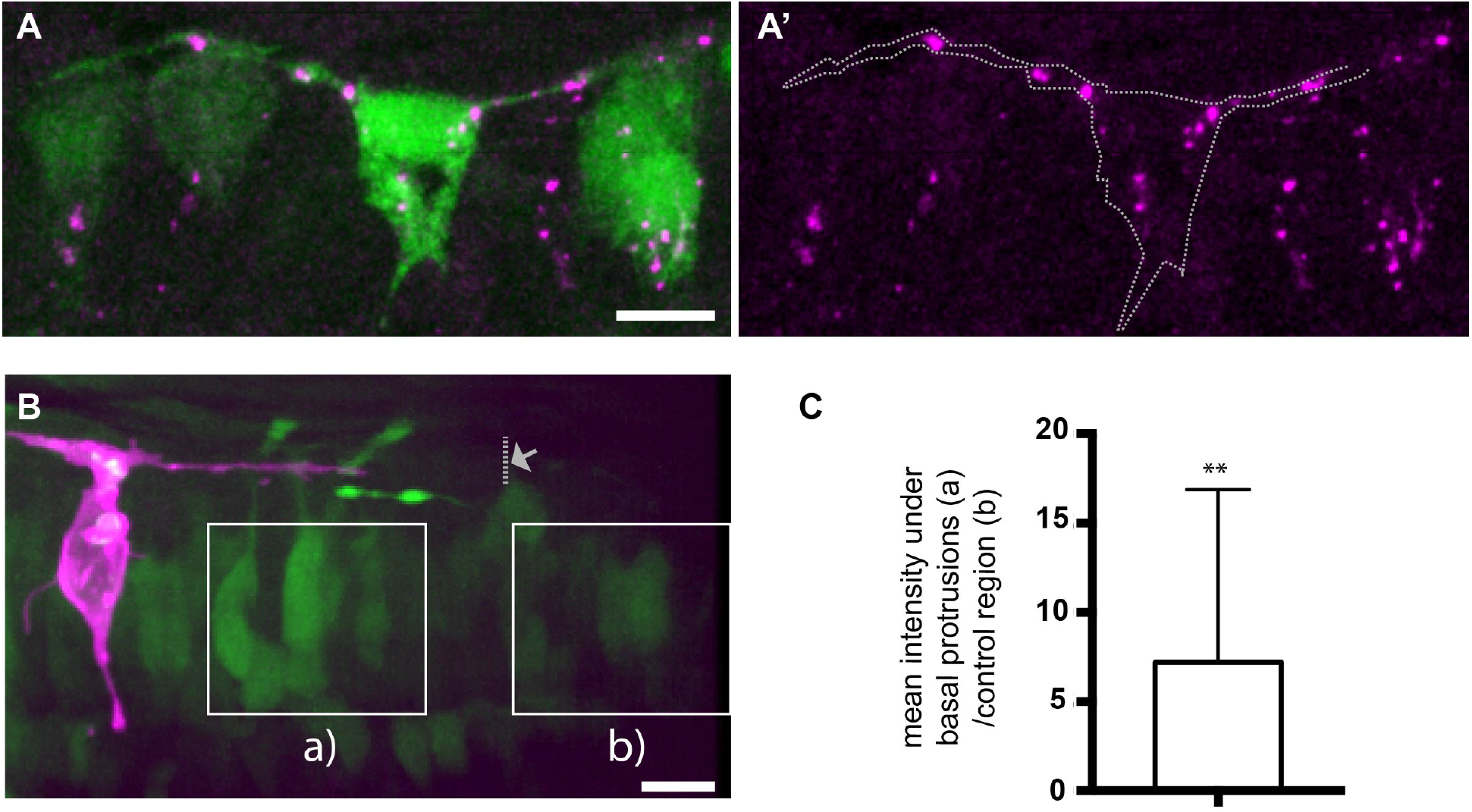
Transient basal protrusion express Delta protein and Notch activity is upregulated in their vicinity. A, A’) DeltaD immunoreactivity (magenta) shows the localisation of DeltaD aggregates in the basal protrusions and cell body of a T-shaped cell. The T-shaped cell expresses cytoplasmic GFP (green) under the DeltaD promotor. B) A T-shaped cell labelled with membrane-mKate (magenta) extends basal protrusions in a Tg(Tp1:VenusPEST) (green) embryo. The maximal extension of one basal protrusions is labelled with an arrow and dotted line. Squares indicate the two areas used for analysis of Tp1:VenusPEST expression. C) Graph showing the relative mean Tp1:VenusPEST fluorescence intensity under the basal protrusions compared to a control region outside the basal protrusions (unpaired 1-tailed t-test, p=0.016, the average (a intensity/ b intensity) is significantly different from 1).

Since basal protrusions extend bidirectionally along the same D/V level as the differentiating cell body, these protrusions will be perfectly placed to preferentially interact with neural progenitors located at the same D/V level (i.e. progenitors likely to generate neurons of the same subtype) and promote the neuronal spacing pattern observed in the zebrafish spinal cord. This suggests the relative positions of neurons of different subtypes could be independent of each other. To test this we measured the relative positions between different neuronal subtypes (*evx1, eng1b* and Vsx1:GFP, Supplementary Figure 3A-H). This analysis revealed that positions of *evx1* and *eng1b* neurons had no consistent alignment with Vsx1:GFP expressing neurons (Supplementary Figure 3F-H), suggesting there is no pre-pattern for the relative position of different neuronal subtypes along the anteroposterior axis, and that regulation of differentiation of a particular neuronal subtype is independent of interactions with neurons of other subtypes.

Together these results are consistent with the existence of a long distance lateral inhibition mechanism that operates between differentiating neurons of the same subtype and their progenitors at the same D/V level. The expression of DeltaD in transient basal protrusions and the increase in Notch activation in cells spatially related to these basal protrusions suggests the basal protrusions could control both the spatial and temporal pattern of differentiation through long distance but transient Notch Delta lateral inhibition.

### Laminin depletion reduces both basal protrusion length and spacing between successively differentiating neurons

To further test whether basal protrusions could regulate the spatiotemporal pattern of Vsx1 neuron differentiation, we modified basal protrusion length and quantified the pattern of neuronal differentiation *in vivo*. Since the transient basal protrusions grow at the basal surface of the neuroepithelium we predicted that extracellular matrix proteins in the basement membrane could be required for their growth. To test this, we monitored neuronal differentiation in *lamc1* mutants that have no detectable Laminin at the basal surface of the neuroepithelium at the developmental stages we are studying. Neurons differentiating in *lamc1* mutant spinal cords develop significantly shorter basal protrusions (mean ± SD, 12.3 ± 4.7 μm, n=39) than neurons in wild type embryos (mean ± SD: 42.6 ± 20.2 μm, n=24 cells, unpaired, two-tailed t-test p<0.0001) (Figure 7A, B, Movie 8), consistent with a role for Laminin in basal protrusion extension.

**Figure 7.**
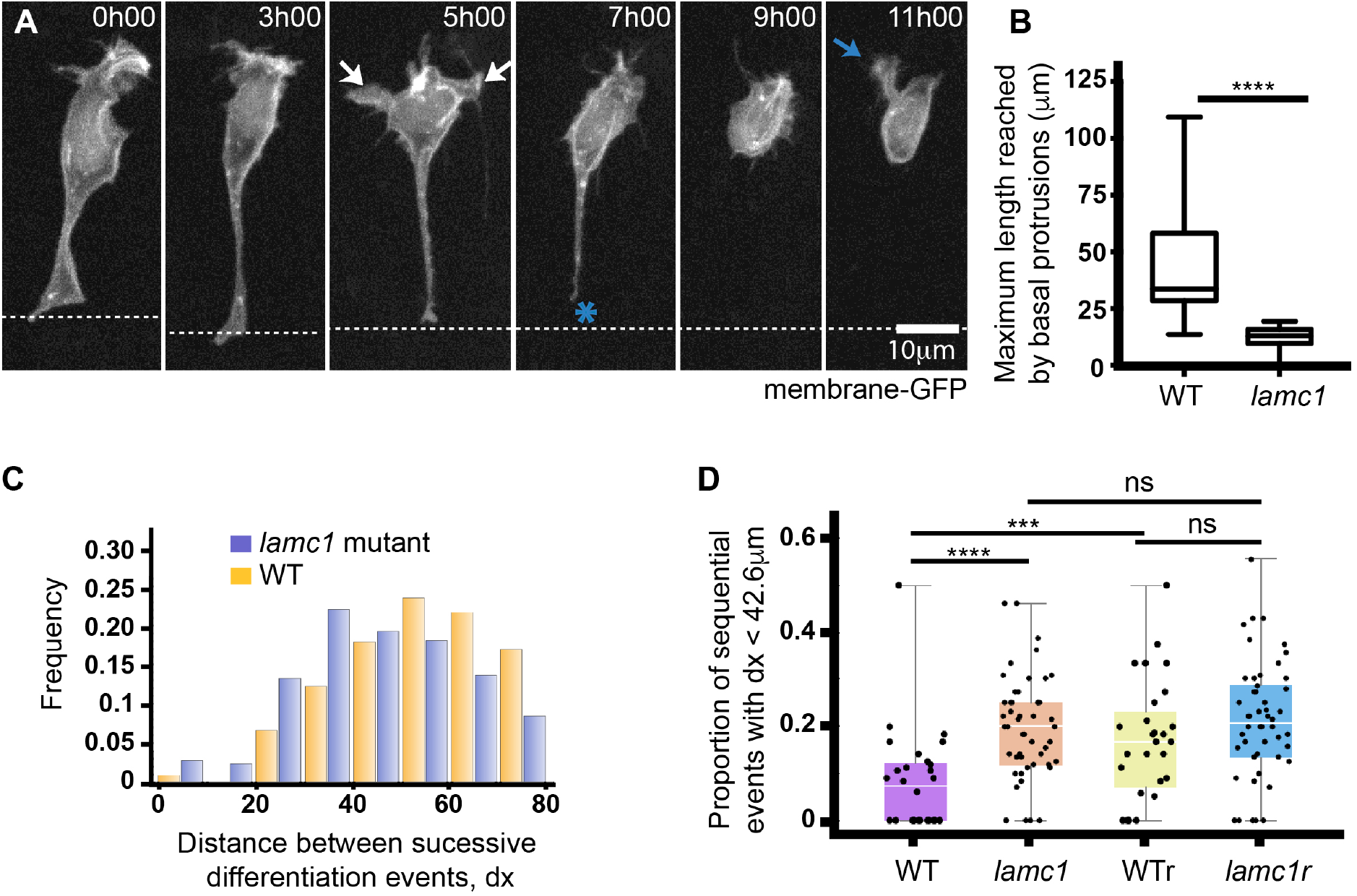
Laminin depletion reduces basal protrusion length and spacing between successively differentiating neurons. A) Time-lapse sequence showing a differentiating neuron in a Laminin-depleted spinal cord (see Movie 8). It has only short basal protrusions (white arrows in time point 5h00). The short basal protrusions are retracted before detachment from apical surface (blue asterisk at 7h00) and axon initiation (blue arrow at 11h00). Cell is labelled with membrane-GFP. View is dorsal. B) Box-and-whisker plot showing the maximum length reached by basal protrusions in WT (mean ± SD, 42.6 ± 20.2μm, n=24 cells) and *lamc1* mutant embryos (mean ± SD, 12.3 ± 4.7μm, n=39). The line inside the box represents the median and whiskers represent minimum and maximum values. Data analysed using unpaired two-tailed t-test (p<0.0001). C) Histogram showing the distribution of the distance between successive Vsx1:GFP differentiation events in wild type (orange) and *lamc1* mutant embryos (purple) (mean ± SD, 54.00 ± 1.52μm in wild type and 45.3 ± 0.99μm in *lamc1*, t-test p-value = 3.16 10^-6^). D) Graph showing the proportion of successive Vsx1:GFP differentiation events that occur within a 42.6 μm interval (the average size of wild type basal protrusions) in wild type embryos, *lamc1* embryos, randomised wild type distributions and randomised *lamc1* distributions. The Kolmogorov-Smirnov test was used to compare wild type and *lamc1* (p = 0.000066); wild type and randomised wild type distributions (p = 0.000224); and, *lamc1* and randomised l*amc1* distributions (p = 0.213).

To determine whether the reduced length of basal protrusions in Laminin depleted embryos could affect the spatiotemporal pattern of neuron differentiation, we performed time-lapse microscopy and compared the pattern of differentiation of Vsx1:GFP neuron pairs in *lamc1* mutants (Supplementary Figure 5B)(n=721 differentiation events in 50 stretches of spinal cord in 25 embryos) and wild type. We found that successive differentiation events occur closer together in *lamc1* mutants than in wild type (Figure 7C, mean ± SD, 54.00 ± 1.52 μm in wild type and 45.3 ± 0.99 μm in *lamc1*, t-test p-value = 3.16 10^-6^), with the highest frequency of these events occurring 30-40μm apart in the mutant compared to 50-60μm apart in the wild type (Figure 5F, 7C).

Since wild type basal protrusions extend 42.6μm on average (and can potentially influence differentiation in this range) we then determined the proportion of sequential differentiation events that occurred within 42.6μm of each other in the wild type and *lamc1* background. This verified that differentiation events are twice as likely to occur within this range in the *lamc1* mutant (0.19 ± 0.11) than in the wild type embryos (0.080 ± 0.11)(Kolmogorov-Smirnov test, p-value = 0.0000666) (Figure 7D). We further compared the wild type and *lamc1* differentiation data to randomly generated differentiation events and found that the proportion of sequential events that occurred within 42.6μm in the wild type, but not the *lamc1* data, is significantly different from random (Kolmogorov-Smirnov test, p = 0.000224 and p = 0.213) (Figure 7D).

To discard the possibility that a decrease in neuronal spacing in *lamc1* mutants is due to an overall increase in neuronal differentiation we quantified the rate of neurogenesis. We determined the ratio between neurons to progenitors (N/P) at early stages of embryonic development and found no difference between wild type and mutant (Supplementary Figure 4A, B). In addition, we analysed the overall organisation of the spinal cord in *lamc1* mutants and showed that patterns of polarity proteins, the locations of progenitor divisions and the location of neuronal differentiation are normal (Supplementary Figure 4A-C). These experiments suggest that gross neuroepithelial organisation and rates of differentiation are normal in *lamc1* mutant embryos at early stages of embryonic development.

Overall these results are consistent with the hypothesis that basal protrusions transiently extend the range of influence of lateral inhibition and longer basal protrusions can regulate differentiation over a longer distance.

### Theoretical predictions support the role of basal protrusions in patterning differentiation through Delta Notch mediated lateral inhibition

To determine whether the pattern of neuronal differentiation can be explained by Delta Notch mediated lateral inhibition delivered via transient basal protrusions, we developed a physical description of lateral inhibition coupled to the observed protrusions dynamics. The dynamics of Delta Notch signalling have been modelled extensively (Binshtok and Sprinzak, 2018). Here we built on (Cohen et al, 2010) and (Collier et al, 1996) and describe the process of lateral inhibition by,

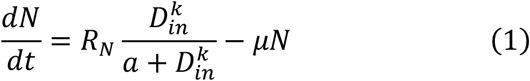

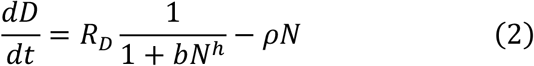

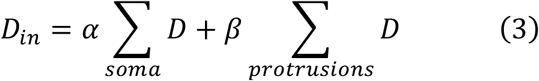

These equations describe the dynamic process of gene activation and inhibition between signaling proteins in contacting cells. *N* and *D* refer to the amount of active Notch and Delta within cells, and *D*_*i*__*n*_ is the total signal received by a cell from all cells it is in contact with. We assume that cells only mediate signalling through their protrusions and set *α* = 0 and *β* = 1. Nonzero values of *α* are considered in the Supplemental Text. We further assume that the probability of neuronal differentiation will correlate with a cell’s level of Delta expression (Hunter et al, 2016) and that neuronal differentiation commences with basal protrusion extension. The temporal and spatial dynamics of basal protrusions follow the experimentally observed dynamics. See Methods and Supplementary Information 1 for further details of the theoretical set up.

We first performed simulations to predict the distribution of *dx* assuming differentiation events occur randomly along the spinal cord (Figure 8A). If differentiation events occur at random, the distance between successive events should also be random. With random differentiation the predicted distribution of *dx* (mean ± SD, 40.90 ± 21.55 μm) differs significantly from both the wild type experimental distribution (mean ± SD, 54.53 ± 18.92 μm; Kolmogorov-Smirnov test, p-value < E^-10^) (compare Figure 8A to Figure 5F), and the *lamc1* mutant distribution (mean ± SD, 46.32 ± 18.68 μm; Kolmogorov-Smirnov test, p-value = 9.6E^-7^) (compare Figure 8A to Figure 7C) confirming that the spatiotemporal patterns of differentiation *in vivo* are unlikely to be randomly generated.

**Figure 8.**
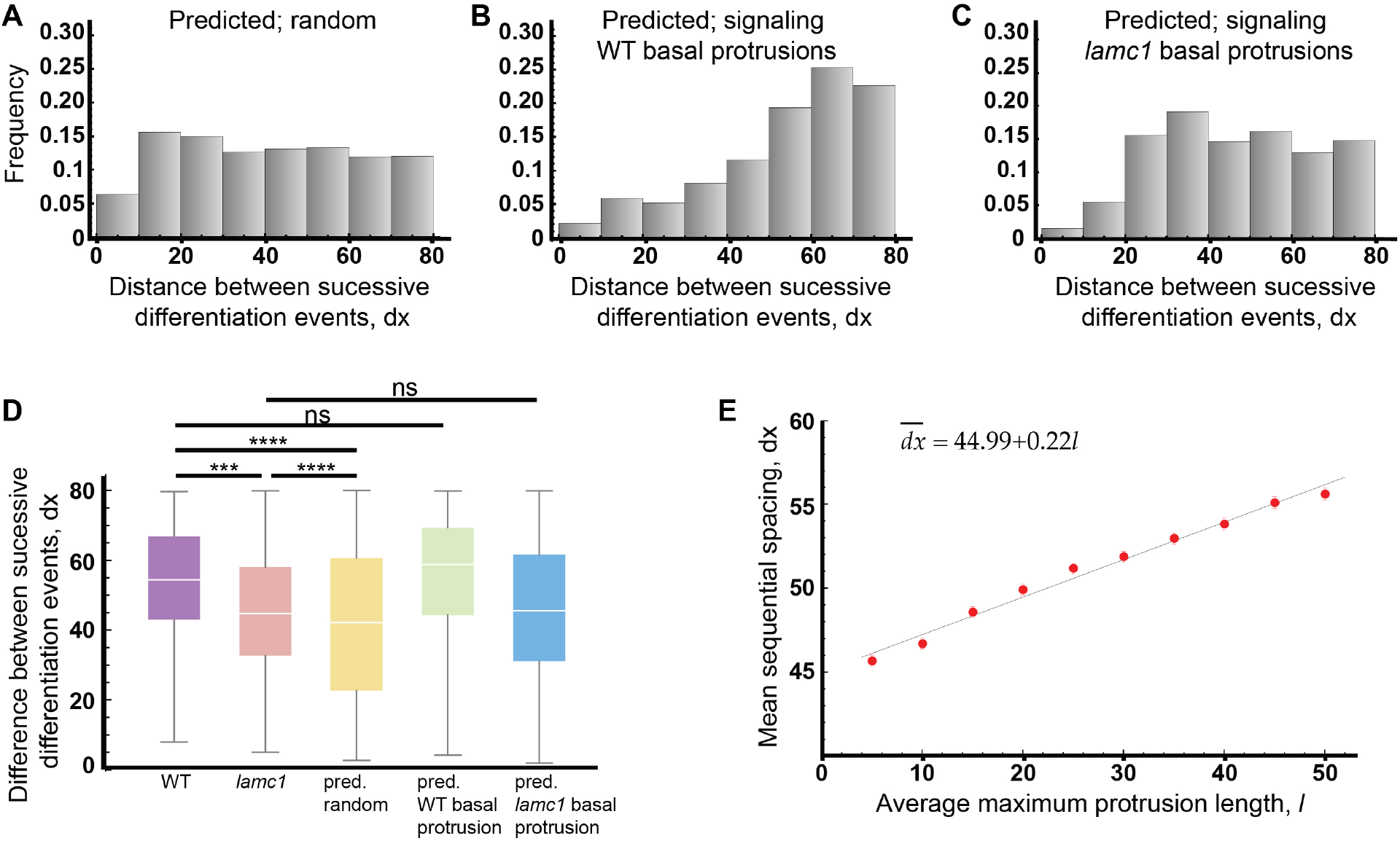
Theoretical predictions support the role of basal protrusions in patterning differentiation through Delta Notch mediated lateral inhibition. A to C) Histograms of the distributions of the distance between successive differentiation events predicted by theoretical model assuming a random distribution of differentiation events (A) (mean ± SD, 40.90 ± 21.55 μm), assuming lateral inhibition signalling occurs through basal protrusions of wild type length (B) (mean ± SD, 54.53 ± 18.92 μm), or, assuming lateral inhibition signalling occurs through basal protrusions of *lamc1* length (C) (mean ± SD, 46.32 ± 18.68 μm). D) Box-and-whisker plots of the distance between successive differentiation events under various *in vivo* conditions and model predictions. The Kolmogorov-Smirnov test was used to compare wild type and lamc1 distribution (p-value = 0.000167), wild type and predicted random distribution (p-value < E^-12^), *lamc1* mutant and predicted random distribution (p-value = 9.6E^-7^), wild type and predicted distribution when basal protrusions of wild type length convey lateral inhibition (p-value = 0.121), *lamc1* mutant and predicted distributions when short and slower (*lamc1* length and dynamics) basal protrusions convey lateral inhibition (p-value = 0.181). E) Predicted relationship between the average maximum length of basal protrusions and the mean distance between sequential differentiation events.

We then performed simulations assuming that the protrusion dynamics follow those of the wild type fish. The predicted distribution between successive differentiation events (*dx*) in this case is in agreement with our experimental measurements (compare Figure 5F and Figure 8B; Kolmogorov-Smirnov test, p-value = 0.121). We repeated the analysis but now assuming that the length and dynamics of protrusions follow those of the Laminin deficient *lamc1* mutant. Now the predicted distribution is in agreement with the distribution of *dx* in the *lamc1* mutant found *in vivo* (compare Figure 7C to Figure 8C; Kolmogorov Smirnov test, p-value = 0.181). Furthermore, the *lamc1* mutant distributions are significantly different to simulations with wild type length protrusions (compare Figure 7C and Figure 8B; Kolmogorov Smirnov test p-value < E^-10^). These results together suggest that the spatiotemporal dynamics of differentiation in wild type and the *lamc1* mutant can both be explained by protrusion mediated lateral inhibition (Figure 8D). The differences in the distribution of *dx* between the wild type and *lamc1* mutant can be explained by differences in basal protrusions length.

In order to understand how changes in basal protrusion length and dynamics impact on the spatiotemporal pattern of differentiation we performed simulations while continuously varying the protrusion length. We found that the average distance between sequential events 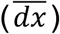 follows a linear relationship with the protrusion length (Figure 8E). However, a given change in the protrusion length, *dl*, only confers a change in the mean spacing 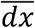 which is about 22% of *dl* (Figure 8E). This can be understood as follows. The protrusions specify a transient region where neurogenesis is inhibited. Although this generates a minimal spacing between sequential events, the events do not have to occur right at the boundary and this alters the mean of the distribution (as seen in the noise around the peaks in Figures 5F, 7C and 8B-C). This effect becomes stronger as the protrusions become smaller which explains why large changes in the protrusion length in the *lamc1* mutant do not produce equally drastic shifts in the average value of dx (Figure 5F and 7C, see Supplementary Text and Figure S13 for detailed explanation). The relative impact of the protrusions on the spacing between sequential events in our region of interest declines for smaller and slower protrusions. These considerations together explain why large changes in the protrusion length in the *lamc1* mutant do not produce equally drastic shifts in the average value of dx (Figure 7C, see Supplementary Text and Figure S13 for detailed mathematical derivation and explanation).

To explore how the position and timing of differentiation are related, we also computed the spatial and temporal relationship between differentiation events. These analyses both *in vivo* and using our theoretical model showed that there is a negative correlation between the distance between two cells and the time at which they differentiate so that cells that are closer in space tend to differentiate further away in time (Supplementary Figures S10 and S11). *In vivo* the wild type and *lamc1* mutant data both followed this trend, however, the range over which this correlation was present in *lamc1* mutants was reduced, consistent with the reduced basal arm length in *lamc1* mutants (Supplementary Figure S10C, F). These spatiotemporal correlations also appear in our theoretical model when long or short basal protrusions mediate lateral inhibition (but not when differentiation occurs randomly), further supporting the role of basal protrusions in patterning neuronal differentiation (Supplementary Figure S11).

Finally, we have performed simulations that assess differentiation patterns when lateral inhibition takes place only at soma-to-soma contacts and a combination of soma and basal protrusion contacts or only via basal protrusion contacts. We found that simulations that include soma-to-soma contacts to mediate lateral inhibition prior to protrusion extension cannot recapitulate our *in vivo* observations (Supplementary Figure S12). This suggests that that soma-to-soma contacts play a minimal role in the mechanism that determine the pattern of differentiation between spinal neurons.

## Discussion

Using live-imaging in zebrafish we have uncovered a new cellular behaviour for vertebrate neurons that regulates the spatiotemporal dynamics of neuronal differentiation along the spinal cord. Differentiating neurons and non-apical progenitors transiently develop two long basal protrusions prior to apical detachment and axogenesis. These basal protrusions express Delta at high levels and activate Notch signaling at a distance from the cell body. The dynamics of basal protrusion extension and retraction are consistent with a role in delivering Delta-Notch mediated lateral inhibition at a distance to regulate the position and time of spinal neuron differentiation. Additionally, previous work has shown Delta expression is required for the sparse spatial pattern of zebrafish spinal neurons (Okigawa et al 2014). We show that experimental manipulation of basal protrusions *in vivo* and in a mathematical model of cells with and without signaling basal protrusions also support the role of basal protrusions in mediating lateral inhibition at a distance to regulate both the position and time of spinal neuron differentiation. Protrusion mediated lateral inhibition has been proposed to control sparse differentiation patterns in the fly peripheral nervous system (De Joussineau et al. 2003; Cohen et al. 2010). Our work demonstrates a similar cell protrusion mediated mechanism operates in the spinal cord of a vertebrate.

The extension and retraction of basal protrusions on spinal neurons is highly stereotyped and is the earliest morphological feature of neuronal differentiation once the nucleus of the newly born neuron has reached the basal surface of the neural tube. Therefore, influencing the differentiative behaviour of surrounding cells is prioritized over other essential neuronal behaviours such as axon outgrowth. Basal protrusions are robust microtubule based processes and always appear in pairs – one directed strictly anteriorly along the spinal cord and one directed strictly posteriorly. In contrast to the random protrusive activity observed on vertebrate neurons differentiating *in vitro* (Dotti et al, 1988), protrusive activity on spinal neurons differentiating *in vivo* is highly directed and predictable. We hypothesize that this directed longitudinal growth of basal protrusions is an effective way to preferentially contact and influence the behavior of neural progenitors at the same dorsoventral level in the spinal cord. Progenitors from the same dorsoventral level will likely generate neurons of the same subtype and this directed basal growth maximizes the chance of influencing differentiation of similar neuronal subtypes. We find non-apical progenitors (Vsx1 expressing progenitors) also undergo this predictable basal protrusive activity prior to their terminal division close to the basal surface of the spinal cord. They will therefore also be able to influence differentiation of similar non-apical progenitors. Thus this morphological transition is another similarity between neurons and non-apical progenitors during their paths to differentiation (McIntosh et al 2017).

Our analyses suggest neuronal basal protrusions deliver Delta mediated lateral inhibition at a distance, a similar role to that proposed for the basal protrusions of SOPs on the fly notum and wing disk (De Joussineau et al. 2003; Cohen et al. 2010, Hunter et al. 2018). Although basal protrusions on SOPs and spinal neurons share some similarities, there are some major differences between the two systems. SOPs radiate thin actin-based filopodia in all directions along the basal surface of the epithelium, while zebrafish neurons develop two substantial microtubule based protrusions that grow in predictable orientations. The basal protrusions of zebrafish neurons often also have filopodia on their surface which may increase the interactions between differentiating cells and their near neighbours. Cell bodies of zebrafish neurons also have filopodia on their surface and although these are much shorter than the basal protrusions. Contrary to dynamic basal filopodia on SOPs, basal protrusions on spinal neurons remain relatively stable and extended for several hours. Importantly however, spinal neuron protrusions are transient and their retraction releases cells from long distance lateral inhibition and allows other neurons to differentiate in the previously inhibited space. This suggests that spinal basal protrusions regulate both the time and space of neuronal differentiation.

Protrusive activity that could influence surrounding cell behaviours has previously been suggested in the rodent cortex. There, Basal Intermediate Progenitors in the rat and mouse subventricular zone have a large number of multidirectional membrane extensions that have alternatively been suggested to sense local factors prior to mitosis (Noctor 2004) or to mediate Delta Notch signaling between Basal Intermediate Progenitors and apical Radial Glia cells, which maintains the proliferative progenitor population (Nelson 2013). Although the protrusions on rodent progenitors do not appear to have a stereotypic orientation and their relation to the spatial and temporal progression of neurogenesis in the cortex has not been assessed, it remains possible they serve similar functions to the basal protrusions of spinal neurons and progenitors. Our observations in the zebrafish telencephalon show new born neurons in this region behave quite differently to spinal neurons. Early telencephalic neurons do not elaborate long basal protrusions prior to axogenesis and there is no obvious long distance spacing pattern of differentiation in this region. Thus programmes of cell morphogenesis and neuronal differentiation are region specific.

Many of the neurons in the spinal cord arise from asymmetrically fated divisions (Alexandre et al 2010, Das and Storey, 2012, Saade et al 2013, Kressmann et al 2015) where daughter cell fate is also regulated by Delta-Notch interactions. In asymmetric divisions Delta-Notch signaling is likely to be mediated exclusively between the sister cells of each division (Dong et al 2012, Kressmann et al 2015). Our modeling suggests that such lateral inhibition between immediate neighbours cannot explain the long distance spacing pattern of neuronal differentiation, nonetheless this local mechanism that operates during progenitor divisions must be integrated with the long distance mechanism delivered through basal protrusions. We have not investigated how these two processes might work together, but we favour the possibility that lateral inhibition through long basal protrusions delays neuron (and non-apical progenitor) differentiation after their birth rather than regulating the time of their birth or particular fate. Our own unpublished data shows that neurons born at the same time begin to express the neuronal transgene HuC:GFP within a very wide time window (from 4 to 12 hours after their birth), thus neurons can progress through their differentiation pathways at very different rates. Prospective neurons can initially maintain high levels of Notch activity and reduction of Notch activation accelerates their differentiation (Baek et al, 2018), raising the possibility that the transient long distance lateral inhibition mediated by basal protrusions controls this time of differentiation but does not change cell fate.

To test the potential for basal protrusions to mediate the spatial pattern of differentiation *in vivo* we examined spinal neuron differentiation in Laminin depleted spinal cords. We found that basal protrusion growth is significantly reduced in the absence of Laminin and this correlates with a predicted reduction in the distance between differentiation events. Laminin depletion did not completely abolish basal protrusions from spinal neurons and we show that the short protrusions that remain can explain the altered spatiotemporal dynamics of differentiation in the mutant. Although we cannot eliminate the possibility that Laminin depletion alters the spatial pattern of differentiation through mechanisms other than reduced basal protrusion length, this experimental approach is consistent with our major hypothesis. The overall architecture and cellular organisation of the Laminin depleted spinal cord is grossly normal and we propose that a Laminin rich extracellular matrix may be required for basal protrusion growth, perhaps in a similar way to Laminin’s proposed role in axonal growth at the basal surface of neuroepithelium (Randlett et al 2011).

A theoretical model that captures the protrusion dynamics in our in vivo system supports the hypothesis that basal protrusions mediate the spatiotemporal pattern of differentiation. We show that protrusion length is proportional to the spacing between successively born neurons. Furthermore, our theory recapitulates our in vivo spatiotemporal patterns in both wild type and laminin depleted measurements.

The biological function of regulating neuronal differentiation in a spatiotemporal manner is unclear. However, we speculate that it may be advantageous for neuronal circuit formation if the initial connections are made between a minimal number of spatially distributed neurons. Later differentiating neurons can then be added to a functioning circuit to consolidate or modify the circuit function. This could be particular important in zebrafish and amphibian embryos as they develop externally and need to quickly build a functional motor circuit for survival.

## Methods

### Animals

The zebrafish wild type (Ekkwill, AB/Tuebingen or Tuepfel long fin), transgenic (Tg(*vsx1*:GFP) (Kimura et al. 2008); Tg(*deltaD:Gal4;UAS:GFP*) (Scheer et al. 2001); Tg(TP1:VenusPEST) (Ninov et al. 2012)) and *lamc1* mutant (*sleepy* (*lamc1*^*sa379*^), Kettleborough et al. 2013) lines were maintained under standard conditions as described in Westerfield, 2000. The embryos were generally raised in water or E2 medium containing 0.003% 1-phenyl-3-(2-thiazolyl)-2-thiourea at 28.5°C.

### *In situ* hybridisation and immunohistochemistry

Zebrafish embryos fixed in 4% PFA at 22hpf, have been processed for whole-mount *in situ* hybridisation according to the protocol described in (Thisse et al. 1998). The antisense RNA DIG probes *eng1b* (Batista et al. 2008), *evx1* (Thaeron et al. 2000), *Vsx1* (Passini et al. 1997) were detected using Fast Red (Roche) substrate. To compare the relative distribution of neuronal subtypes, we performed *in situ* hybridisation for *eng1b* or *evx1* in the Tg(*vsx1*:GFP) transgenic embryos followed by the detection of GFP expression by immunohistochemistry (chicken anti-GFP, Abcam, diluted 1:1000).

Whole-mount immunohistochemistry was performed on 22-28 hpf wild type, *lamc1* mutant or Tg(*deltaD:Gal4;UAS:GFP*) (Scheer et al. 2001) embryos. Embryos were fixed for 2 hours at room temperature in 4% PFA. Primary antibodies used were HuC/D (mouse anti-HuC/D, Invitrogen, diluted 1:100), aPKC (rabbit anti-aPKC, Santa Cruz Biotechnology, diluted 1:500) or DeltaD (mouse anti-DeltaD, zdd2, Cancer Research Technology, diluted 1:30). The embryos were incubated with primary antibody for 2 to 3 days at 4°C in a solution containing PBS Triton 0.5%, 2% BSA and 10% goat serum (detailed protocol described in Wright et al. 2011). Sytox Green (ThermoFischer Scientific, diluted 1:5,000) was added to secondary antibody to label nuclei.

### Live imaging and image processing

To monitor individual cells, we injected mRNAs coding for membrane tagged-RFP (m-RFP), m-GFP, m-mKate2, nuclear tagged RFP (n-RFP) or Eb3-GFP in zebrafish embryos at 16-64-cell stage. To characterise the dynamics of neuronal differentiation we followed the differentiation of Vsx1:GFP-expressing neurons in the transgene Tg(*vsx1:GFP*) or *lamc1*^*sa379-/-*^;Tg(*vsx1:GFP*) embryos. Prior to imaging, embryos were anaesthetised in MS-222 (Sigma) and mounted for live-imaging as explained in Alexandre et al, 2010. Embryos expressing the transgenes Tg(*vsx1*:GFP) (Kimura et al., 2008) and Tg(TP1:VenusPEST) (Ninov et al, 2012) were imaged either on SP5 confocal (Leica), a Spinning disk confocal (PerkinElmer) or LSM 880 laser scanning confocal (Zeiss) microscopes using 20x water immersion objective with a numerical aperture (NA) of 0.95 or higher and an environmental chamber heated at 28.5C. A series of z-stacks were obtained every 3 to 8 minutes for between 3 to 20 hours depending on the experiment. All the images and movies shown in the present manuscript result from small projections of confocal Z-stacks created using ImageJ. Extra cells were occasionally removed from the field of view using Fiji to show examples of individual cells clearly. Volocity software (PerkinElmer) was used for measurements in 4D.

### Quantification and statistical analysis

The length of basal protrusions and the distances between neurons were measured in 3D at single and multiple time points using Volocity software.

The individual basal protrusions were measured from the cell body to the periphery of the basal protrusion. The maximum overall length reached by basal protrusions includes the cell body width. To compare the maximum average length of cellular protrusions in neuronal and non-neuronal cells we used Kruskal-Wallis with Dunn’s multiple comparison test. To compare the average maximum distance reached by basal protrusion in the wild type and *lamc1* mutant we applied the unpaired, two-tailed T-test.

The distances between successive Vsx1 differentiation events are determined by measuring the distance (dx) between the last and the next neuron born within an 80μm and 42.6μm space interval. We used the Kolmogorov-Smirnov test to compare the differences in distribution of successive Vsx1 differentiation events between wild type, *lamc1* mutant and simulated data and unpaired two-tailed t-test to compare their means. We used the Kolmogorov-Smirnov test to compare the proportion of successive differentiation events occurring within 42.6μm in wild type and *lamc1* mutant. The relative position of different neuronal subtypes was analysed using the Kruskal-Wallis with Dunn’s multiple comparison test.

To compare the intensity of Tg(TP1:VenusPEST) in the vicinity and away from the influence of the basal protrusions, we produced small z-projections, corrected drift and subtracted the background using Fiji. We used Fiji to measure the mean intensity values 2 hours after the basal protrusions reached their maximum length and analysed only the area that has been in contact with the basal protrusions for at least 1h, and away from the neuronal cell body. For each case we calculated the ratio between the mean intensity under basal protrusions and control region (away from the basal protrusions). Statistical analysis was performed using one-tail paired t-test.

### Mathematical and computational details

We used a mathematical model to simulate Notch-Delta mediated lateral inhibition. The model, as defined by equations (1-3), describes the dynamics gene activation and inhibition via cell-cell signalling. We applied the model to a 1D array of cells of variable size following the measured size distribution. The signalling dynamics in individual cells could then be fully defined by the coupled system of differential equations (1-3). Cells could make contact at the soma cell membranes and / or via basal cellular protrusions. We modelled basal protrusion dynamics by allowing cells to extend protrusions if their Notch expression falls below a threshold (Hunter et al, 2016). Differentiating cells send but do not receive a signal (Sprinzak et al 2010, 2011). Protrusions were extended at a constant rate and stopped growing when they reached length > *l*_*max*_where *l*_*max*_was sampled from a normal distribution with mean 42.6μm and s.d. 20.2μm following the *in vivo* measurements for maximum basal protrusion length. Once maximum length was reached the protrusions retracted at a rate 1.7 faster than the extension rate (following *in vivo* dynamics). For the *lamc1* mutants we modified the distribution of *l*_*max*_ to follow the mutant distribution with mean 12.3 μm s.d. 4.7*μ*m and implemented extension and retraction rates that were 1.4 times slower than the WT and retraction rates 2.5 times slower than the WT, following the rates measured experimentally. A cell was assumed to have differentiated when both its right and left protrusion were fully retracted. Differentiated cells no longer participated in signalling and we ran simulations until all virtual cells underwent differentiation. The model was solved numerically using the Euler method. Furthermore, a Gaussian noise term was applied to initiate protein concentrations and to the concentrations at each time step in the simulation. The numerical simulations produced a differentiation time for each individual cell together with its position. We used this information to compute (*dx*) as described above and used the Kolmogorov-Smirnov test to compare simulated to experimental data. We repeated each simulated experiment 10 times to mitigate noise and improve statistical power. The analysis of the simulated and experimental data was performed using scripts written on Wolfram Mathematica. Further details on computational methods and algorithm can be found in the SI text.

## Supporting information

All supplemental information

Movie 1:Differentiating spinal neurons transiently two long basal protrusions prior to apical detachment.

Movie 2: Basal protrusions contain dynamic microtubules.

Movie 3: Basal protrusions form filopodia.

Movie 4: Axon formation follows basal protrusion retraction.

Movie 5: Non-apical progenitors also extend basal protrusions prior to apical detachment.

Movie 6: Differentiating telencephalic neurons do not form long transient basal protrusions.

Movie 7: New Vsx1:GFP neuronal precursors appear in the gaps between already existing Vsx1 neurons.

Movie 8: Basal protrusions length is reduced in lamc1 mutant background.

## Acknowledgements

We thank Buzz Baum for discussions and comments on the manuscript, Christian Sousa-Reid and Julian Lewis for the DeltaD antibody, Kate Lewis for the plasmids to produce the antisense probes for the *in situ* Hybridisation, Gavin Wright for the plasmid containing DeltaD cDNA, Bill Harris for giving us the tg(Vsx1:GFP) line, Robert Knight and Sami Sultan for providing the tg(TP1:VenusPEST) embryos, and Wellcome Trust Sanger Zebrafish Mutation Project for the *lamc1* mutant. Leopold Laurer and Laura Ward for Figures S4B and S4C. ZH thanks Nicolas Levernier for discussions. We also thank the fish facilities at UCL and KCL and the imaging facility at UCL GOS UCL. This research was funded by an EPSRC Fellowship (EP/L50488/) and HFSP Long Term Fellowship to ZH, an MRC studentship to R McIntosh, a Wellcome Investigator award to JC, Royal Society fellowship to PA, and NIHR Great Ormond Street Hospital Biomedical Research Centre. The views expressed are those of the authors and not necessarily those of the NHS, the NIHR or the Department of Health.

## Author Contributions

PA and JC conceived the project, supervised the work and wrote the manuscript. PA characterised the individual neuronal behaviours and prepared figures and movies. PA and R Moore characterised the neuronal behaviours and Vsx1 distribution in *lamc1* mutants. R McIntosh characterised the Vsx1 distribution in wild type embryos. PA and R Moore obtained the Delta and Tg(TP1:VenusPEST) experimental data. GG, PA and ZM developed the method to quantify Tg(TP1:VenusPEST). ZH developed the mathematical model, wrote all computational code, performed simulations, did most of the statistical analysis and contributed to writing the manuscript. PA and ZH processed and analysed all the Vsx1 experimental data. All authors commented on the manuscript.

## Supplemental Movies

**Movie 1: Differentiating spinal neurons transiently two long basal protrusions prior to apical detachment.**

Dorsal view of maximum projection of confocal stack. Central cell differentiates through the T-shaped transition into a neuron with commissural axon. Transient protrusions from the differentiating cell can be seen at the basal surface from 102 mins to 348 mins. Basal surface to top.

**Movie 2: Basal protrusions contain dynamic microtubules.**

Dorsal view of maximum projection of confocal stack. The plus ends of microtubules are labelled with EB3-GFP and are seen penetrating the basal protrusions. Cell membrane labelled in magenta in first frame. Basal surface to top.

**Movie 3: Basal protrusions form filopodia.**

Dorsal view of maximum projection of confocal stack. Membrane labelled cell in t-shaped transition showing dynamic filopodial extensions from basal protrusions. Filopodia of variable length largely extend towards the apical surface. Basal surface to the top.

**Movie 4: Axon formation follows basal protrusion retraction.**

Lateral view of maximum projection of confocal stack. Membrane labelled differentiating spinal neuron is seen through the chevron shaped myotome. At time 0min two basal protrusions are already formed and the third shorter protrusion is directed back to the apical surface. These three protrusions are all retracted by timepoint 84min and ventrally protruding axon is seen at time 126min. Growth cone reaches and crosses the ventral floorplate, while a second growth cone from a different neuron on the contralateral side of the neural tube enters the frame at time 161min. Dorsal to top.

**Movie 5: Non-apical progenitors also extend basal protrusions prior to apical detachment.**

Dorsal view of maximum projection of confocal stack. At time 0min a membrane labelled cell has already extended two basal protrusions. At 244min the apical process is detached. Basal protrusions are almost completely retracted by time 372min when the cell undergoes mitosis. Axons can be seen extending from the two daughter neurons at 556min and 756min. Dorsal to top.

**Movie 6: Differentiating telencephalic neurons do not form long transient basal protrusions.**

Dorsal view of maximum projection of confocal stack. Two membrane labelled neurons are attached to apical surface at time 0min. They are both detached from the apical surface at time 75min. Neither generates long basal protrusions before axogenesis at time 52.5min and 90min. Basal to top.

**Movie 7: New Vsx1:GFP neuronal precursors appear in the gaps between already existing Vsx1 neurons.**

Dorsal view of maximum projection of confocal stack. Strong Vsx1:GFP expression in two widely spaced pairs of neurons at time 0min. Sequentially differentiating Vsx1:GFP neurons appear in between the initial pairs and increase their GFP expression in the subsequent 222 minutes. Axon growth can be seen at the basal surface from time 96min.

**Movie 8: Basal protrusions length is reduced in *lamc1* mutant background.**

Dorsal view of maximum projection of confocal stack. Membrane labelled differentiating neuron in a *lamc1* mutant. Apical process detaches at time 75min. Very short basal protrusions visible at time 150min. Axon growth seen from time 495min. Basal to top.

