## Supplementary material for "Basal protrusions mediate spatiotemporal patterns of spinal neuron differentiation": All supplemental information

#### Figure S1

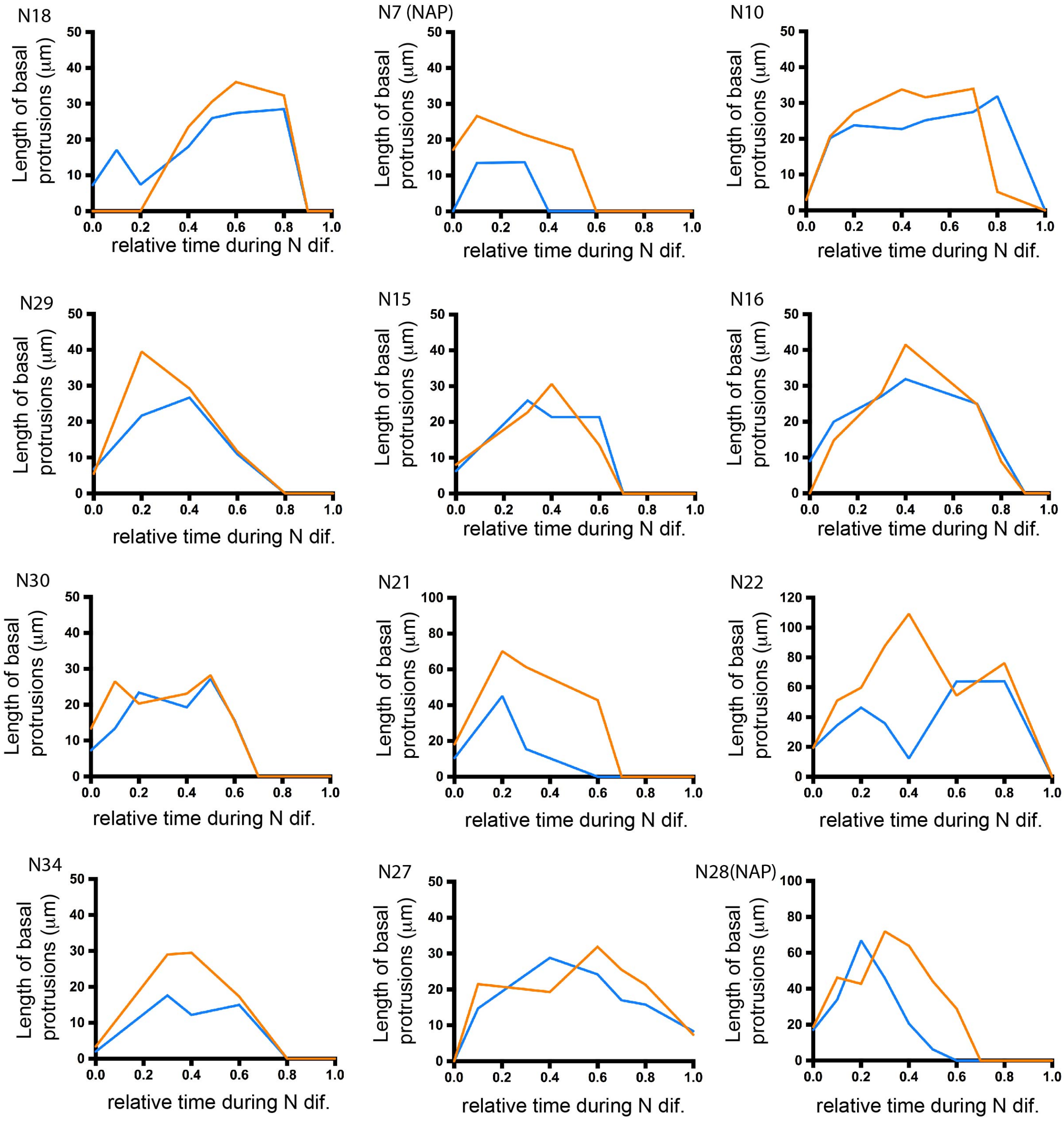

**The dynamics of basal protrusions growth during neuronal differentiation**

Graphs showing the change in length over time of the left (orange) and right (blue) basal protrusions from individual NAPs and individual differentiating neurons. The time has been normalised from (0), the moment in which differentiating neurons begin elongation of basal protrusions, to (1), when neurons initiate axon formation.

#### Figure S2

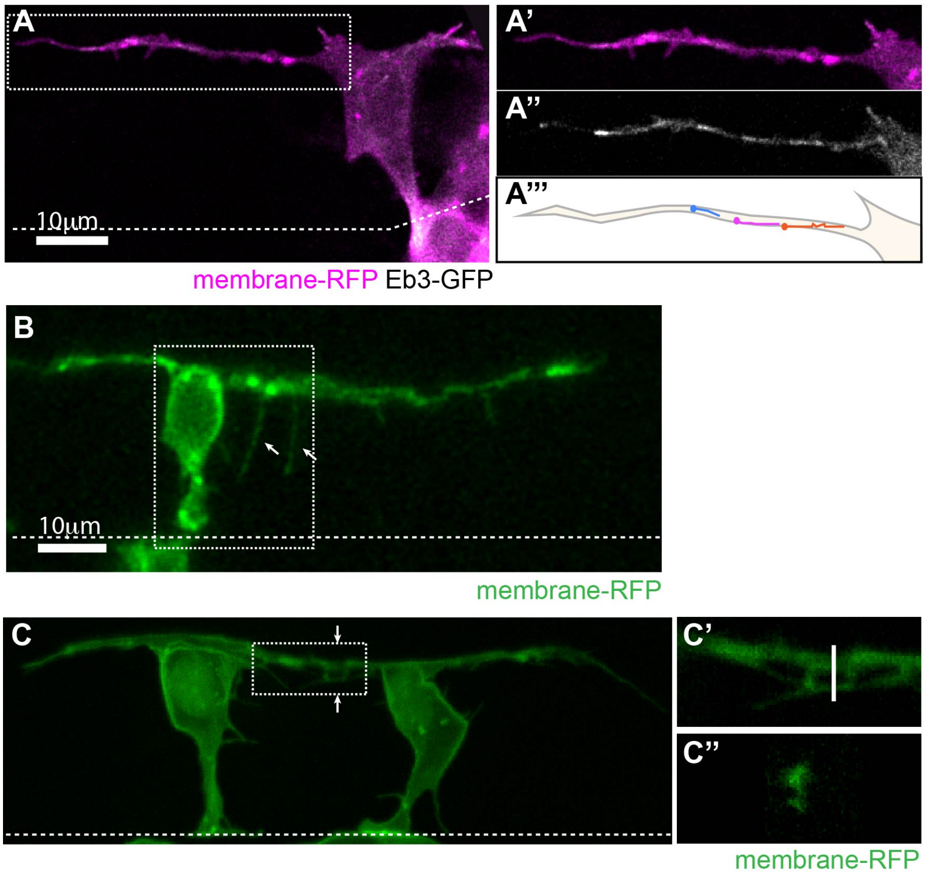

**Basal protrusions contain dynamic microtubules, form filopodia and can overlap.**

A) T-shaped differentiating neuron with a long basal protrusion. White box marks the area of higher magnification shown in A'-A'''). A') Membrane-RFP. A'') EB3-GFP (Movie 2). A''') Illustration of EB3-GFP comet trajectories during a 30 second time-lapse period.

B) T-shaped cell with filopodia (arrows) on basal protrusion (Movie 3).

C) Two neurons differentiating 18μm apart with overlapping basal arms. C') Higher magnification of white box in C). C'') Cross-section of the basal protrusions at the line shown in C').

All images are projected images from confocal z-stacks. Dashed line shows the apical surface.

#### Figure S3

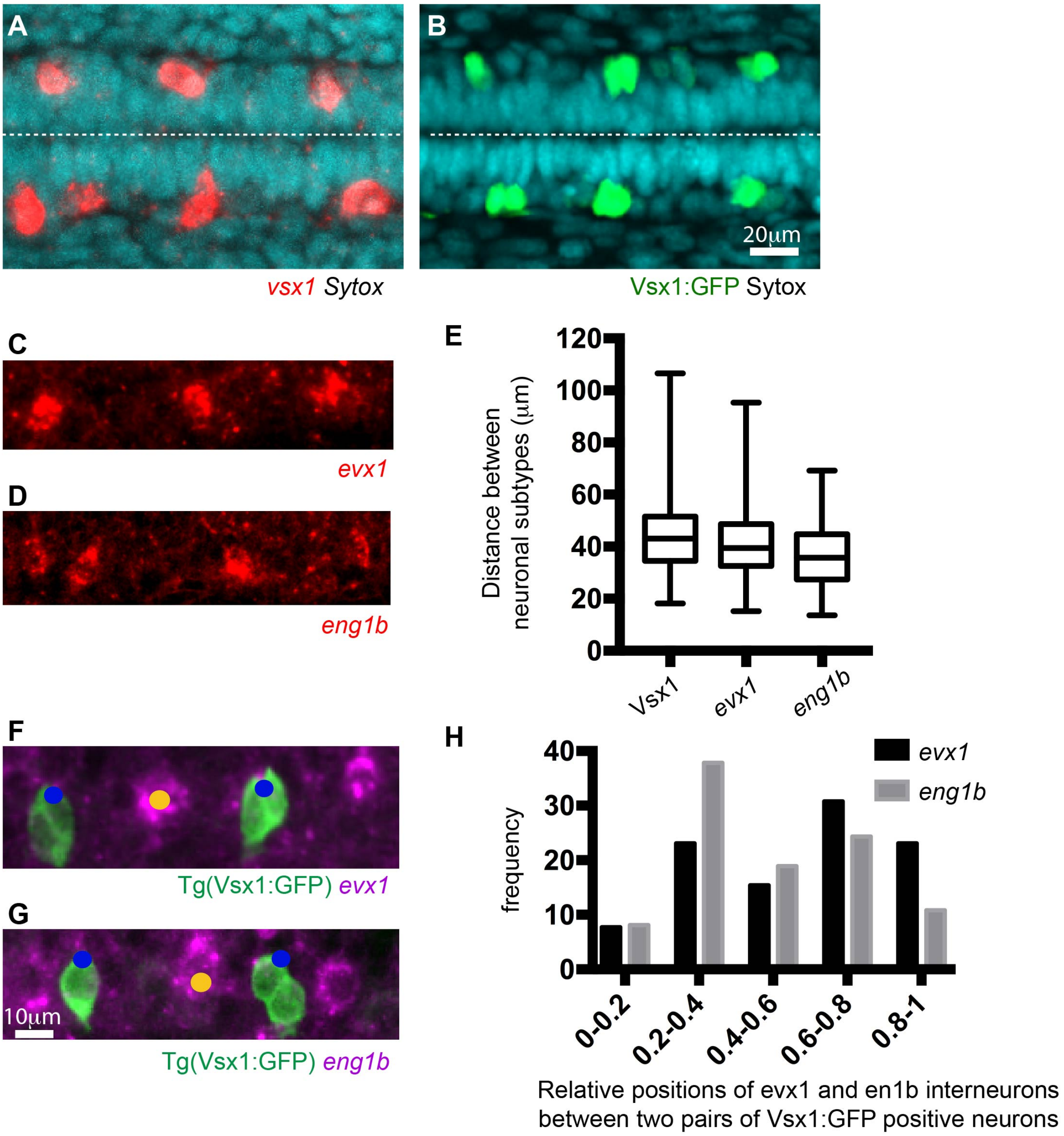

**Neuronal spacing pattern in the zebrafish spinal cord**

A) and B) Dorsal views of Vsx1 expressing cells revealed by in situ hybridization (A) and Vsx1:GFP transgene expression (B) at 20 hfp. Dashed line shows the apical surface.
C) and D) Lateral views of evx1 (C) and eng1b (D) expressing spinal neurons revealed by in situ hybridization at 20hpf.
E) Box-and-whisker plot showing the distance between neurons of the same subtype in the most caudal spinal region at 20hpf (mean ± SD; Vsx1:GFP: 44.4 ± 14.1 μm, n=18; evx1: 41 ± 15.4 μm, n=7; eng1b: 35.7 ± 13.3 μm, n=5 embryos). The line inside the box represents the median and whiskers represent minimum and

maximum values. Data analysed using Kruskal-Wallis with Dunn's multiple comparison test (non-significant).
F and G) Dorsolateral views showing the relative positions of Vsx1:GFP and evx1 (F) or eng1b (G) expressing cells revealed by in situ hybridisation.
H) Frequency distribution chart shows the relative positions of evx1 and en1b expressing interneurons between two pairs of Vsx1:GFP expressing neurons. The distance between the two pairs of Vsx1:GFP neurons has been normalised from 0 (anterior) to 1 (posterior). All evx1 and eng1b expressing neurons that shared the same position with a Vsx1 neuron have been included in position (1).

#### Figure S4

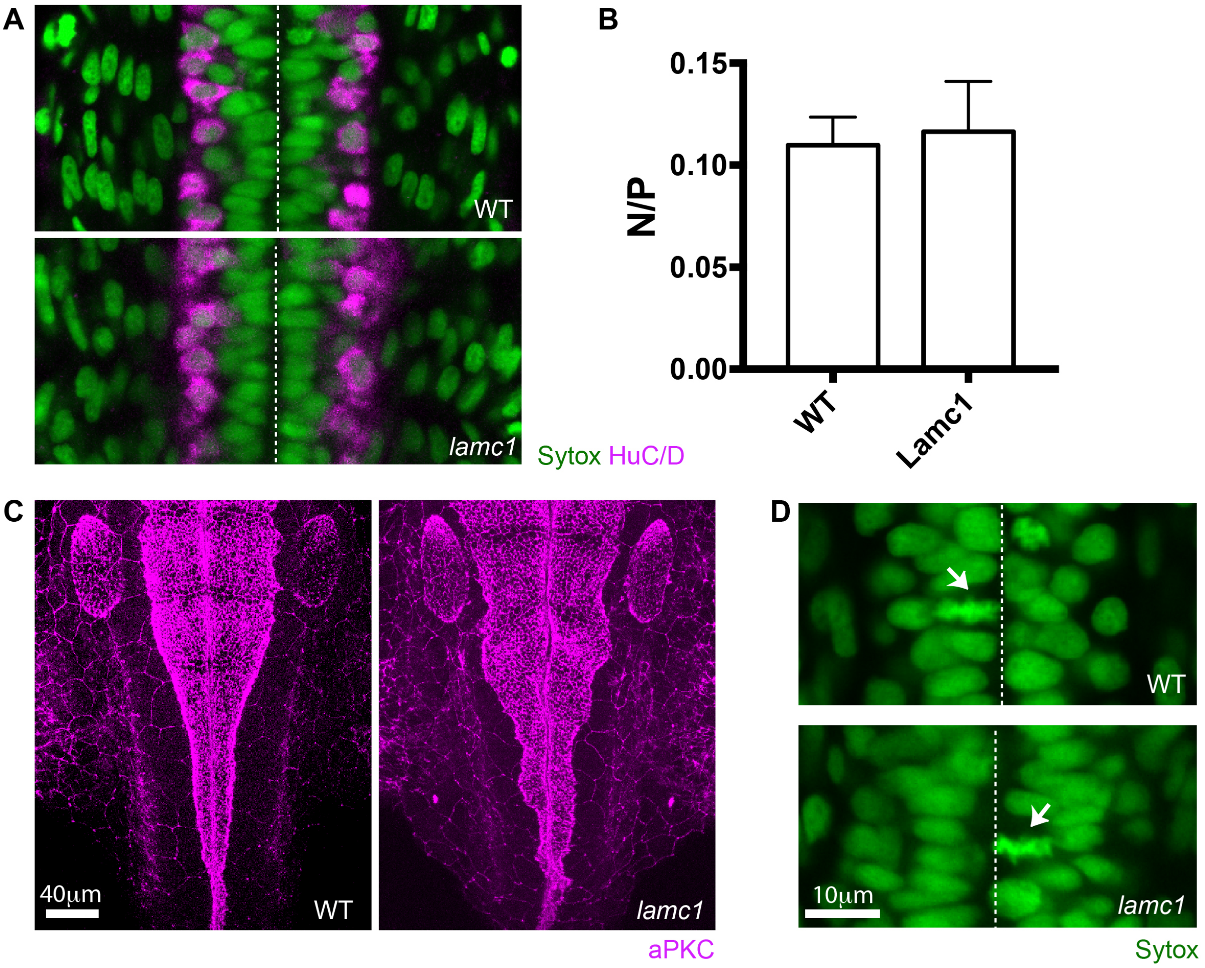

***Lamc1* mutant characterization**

A) Dorsal view of spinal cord in wild type and *lamC1* mutant embryos. HuC/D immunoreactivity (magenta) reveals the position of spinal cord neurons at 24 hpf.
B) Graph showing the ratio of neurons to progenitors in wild type and *lamC1* mutant spinal cords at 24 hpf (mean ± SD, 0.11 ± 0.01 for wild type and 0.12 ± 0.02 for *lamc1* mutant, unpaired two-tailed t-test p=0.8857).
C) Dorsal view of wild type and *lamC1* mutant embryos at the hindbrain. aPKC immunoreactivity (magenta) shows the position of the apical surfaces at 28 hpf.
D) Dorsal view of spinal cord in wild type and *lamC1* mutant embryos showing apical mitoses (arrows).
Dashed line shows position of the apical surfaces in A) and D). Nuclei labelled by Sytox in A) and D).

**Figure S5A**

### Spatiotemporal pattern diagrams illustrating Vsx1:GFP neuronal differentiation events in time and space for 34 stretches of developing zebrafish spinal cord.

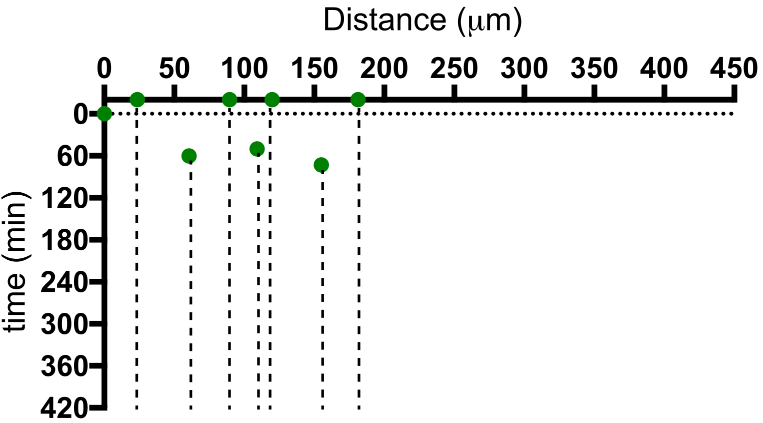

WT emb BML102 a1 a

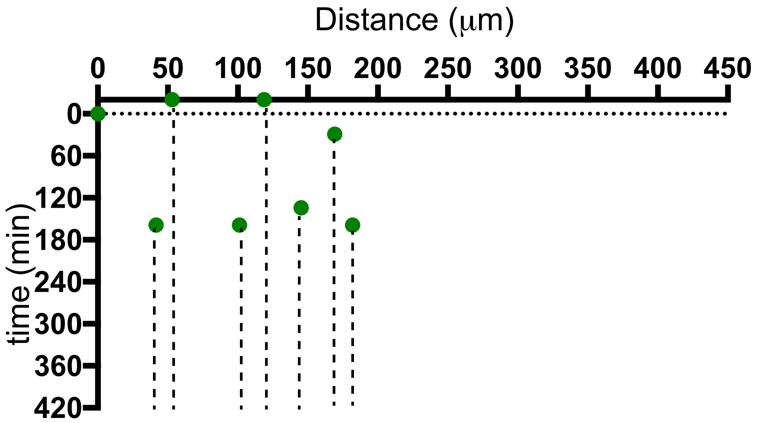

WT emb BML102 a1 b

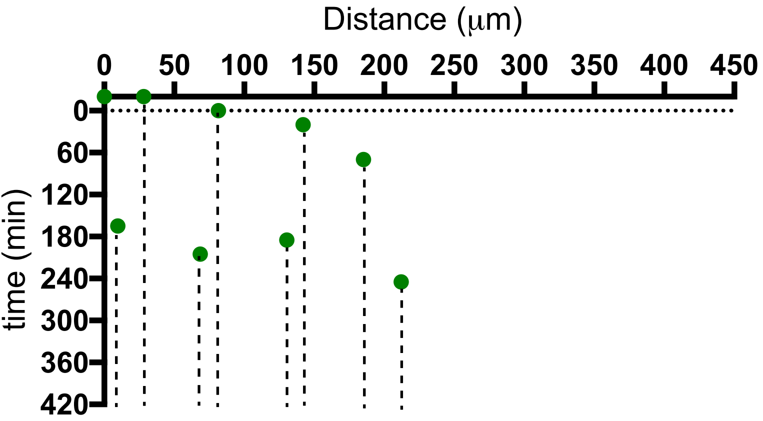

WT emb BML102 a2 b

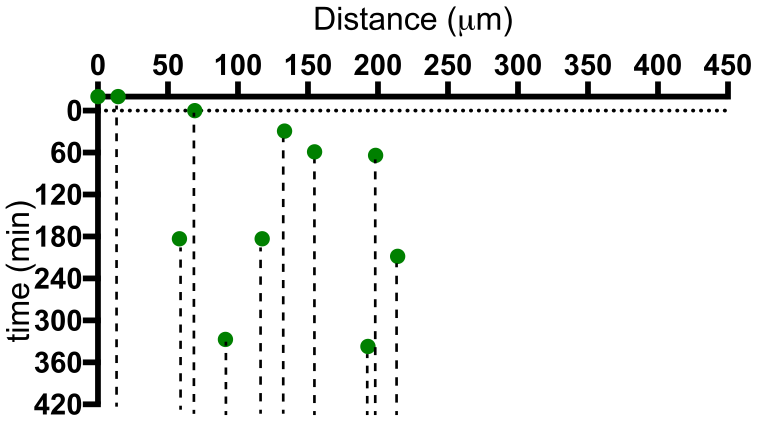

WT emb BML102 a2 a

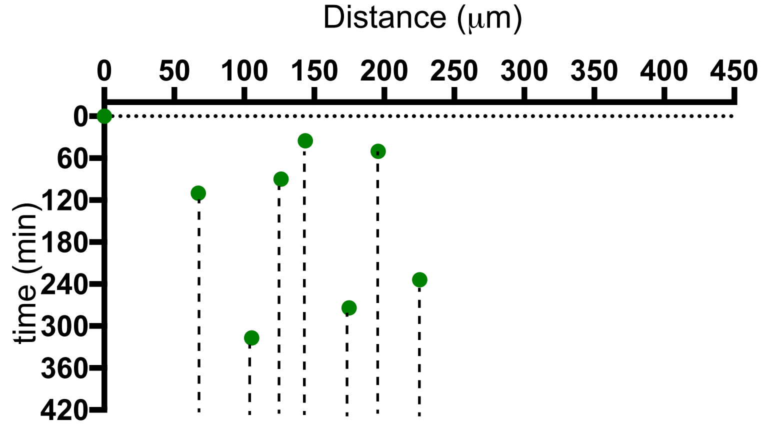

WT emb BML102 b2 a

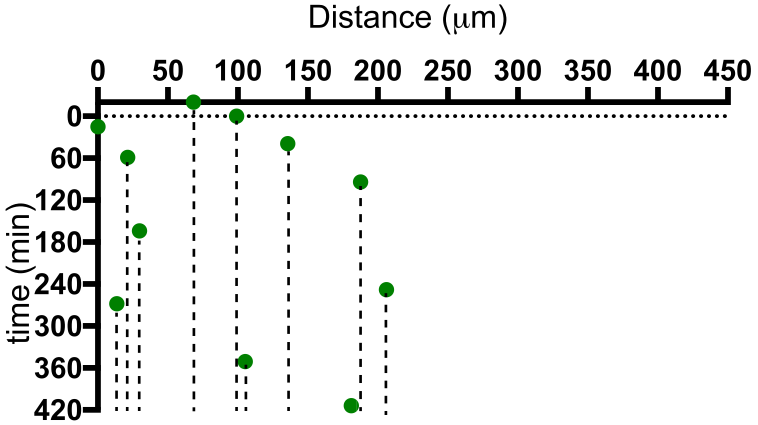

WT emb BML102 b2 b

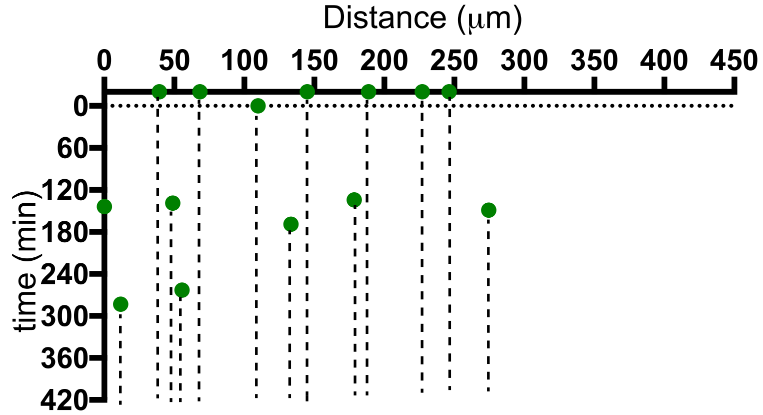

WT emb BML102 b1 a

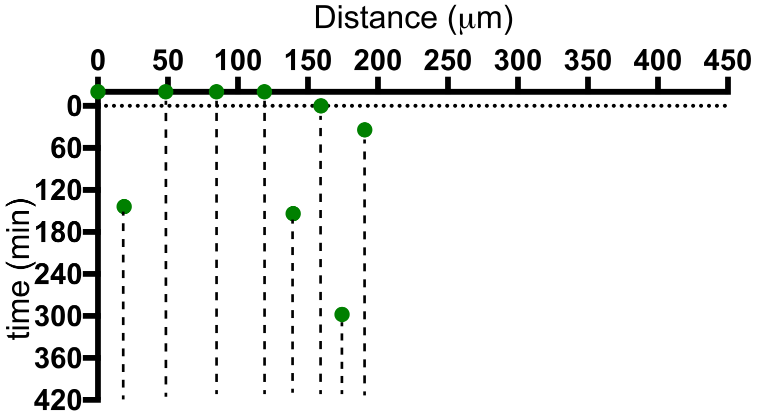

WT emb BML102 b1 b

WT emb1 BML110 a

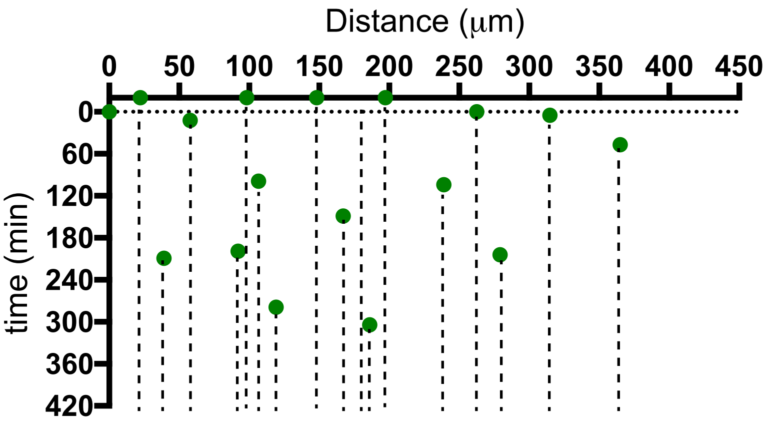

WT emb1 BML110 b

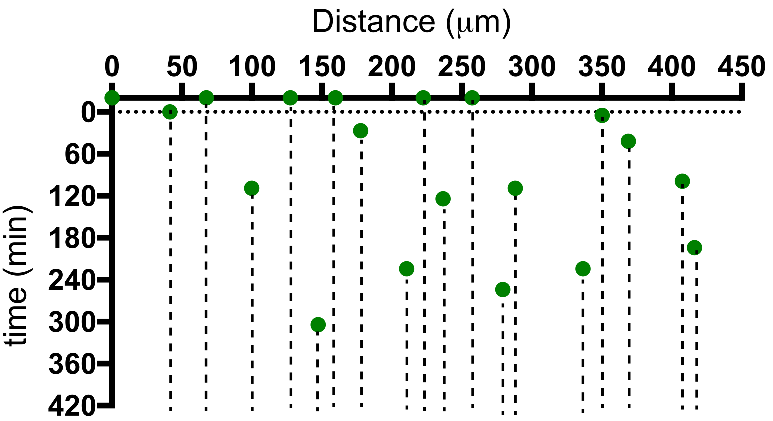

WT emb2 BML110 a

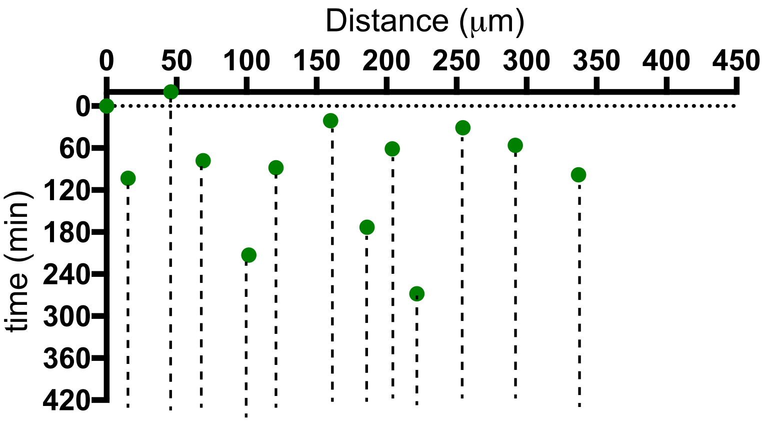

WT emb2 BML110 b

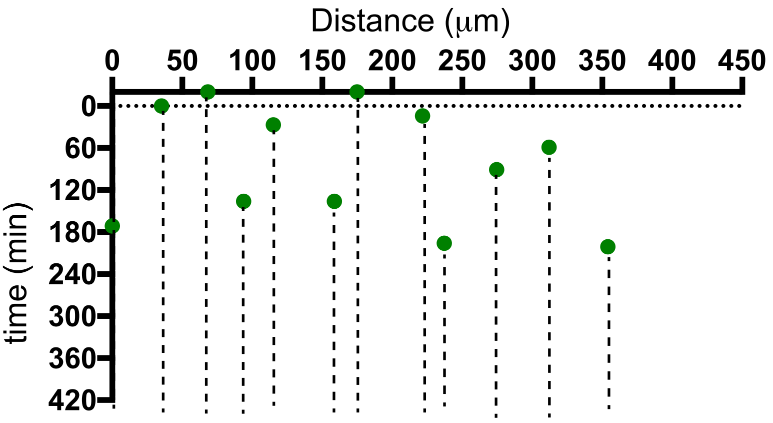

WT emb3 BML110 a

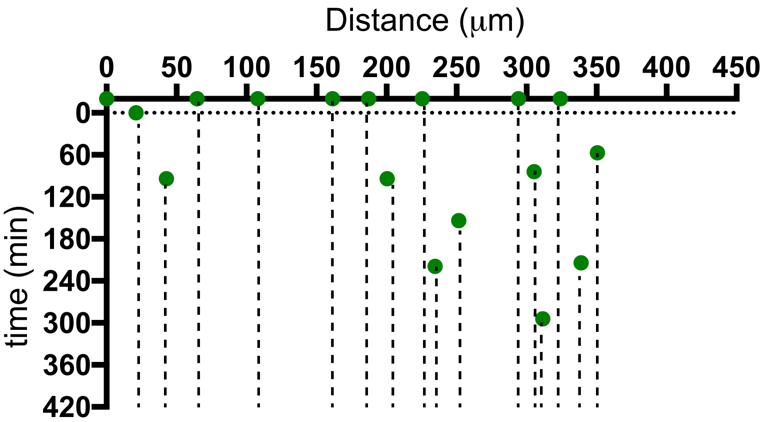

WT emb3 BML110 b

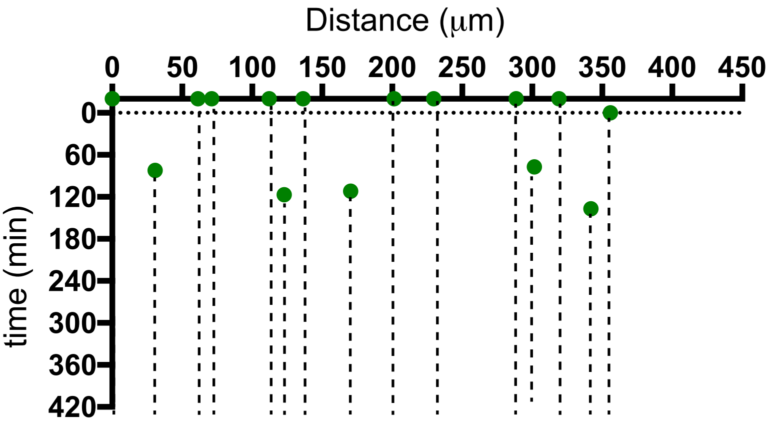

WT emb4 BML110 a

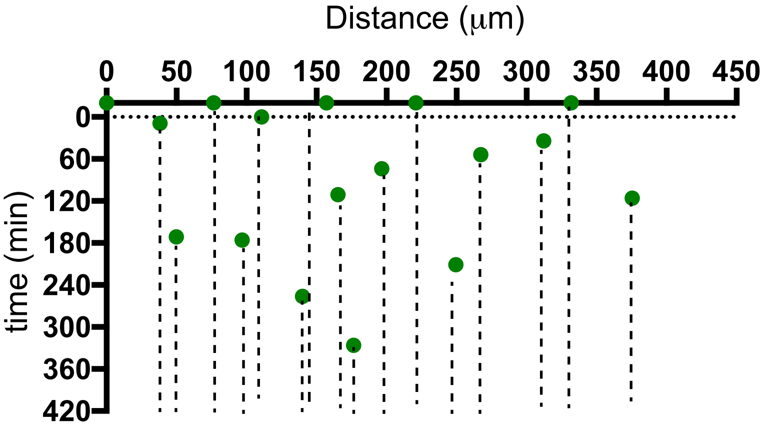

WT emb4 BML110 b

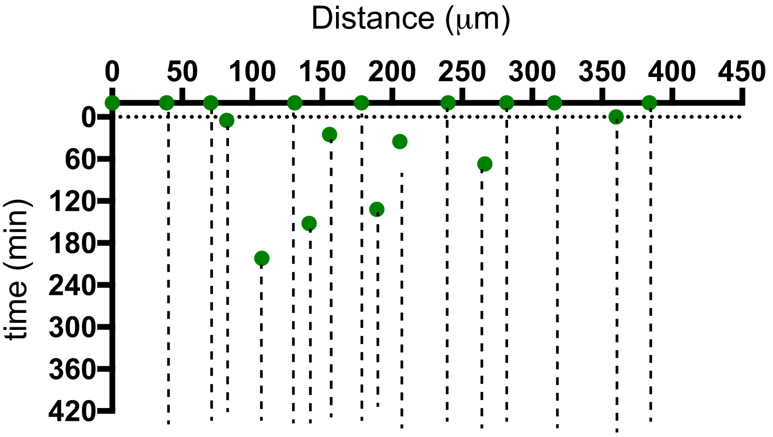

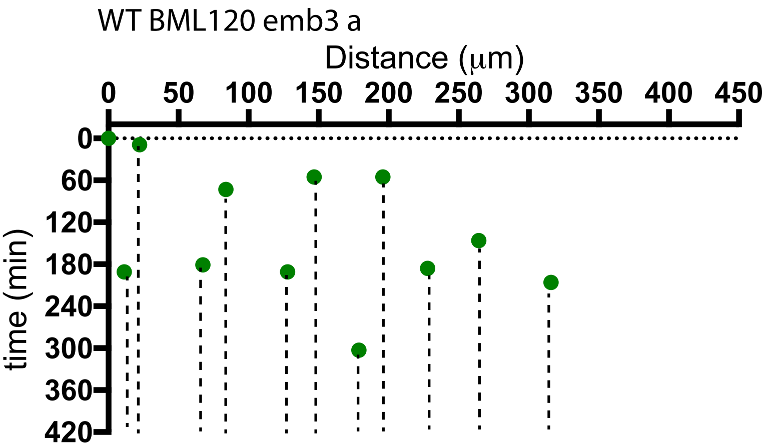
WT BML120 emb3 a
Distance (μm)
0
50
100
150
200
250
300
350
400
450
time (min)
0
60
120
180
240
300
360
420

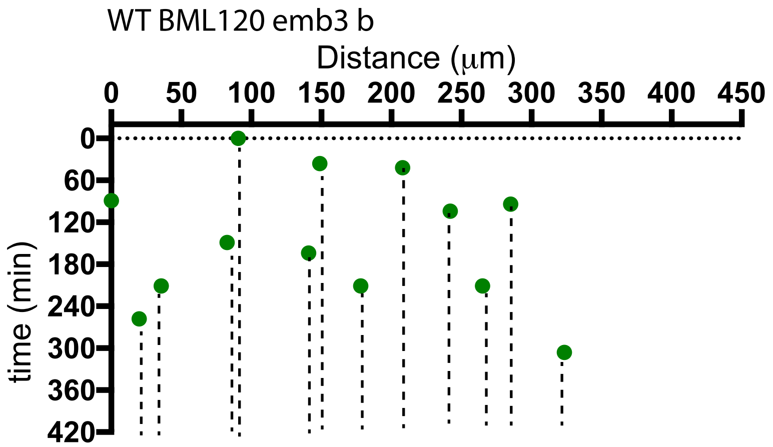
WT BML120 emb3 b
Distance (μm)
0
50
100
150
200
250
300
350
400
450
time (min)
0
60
120
180
240
300
360
420

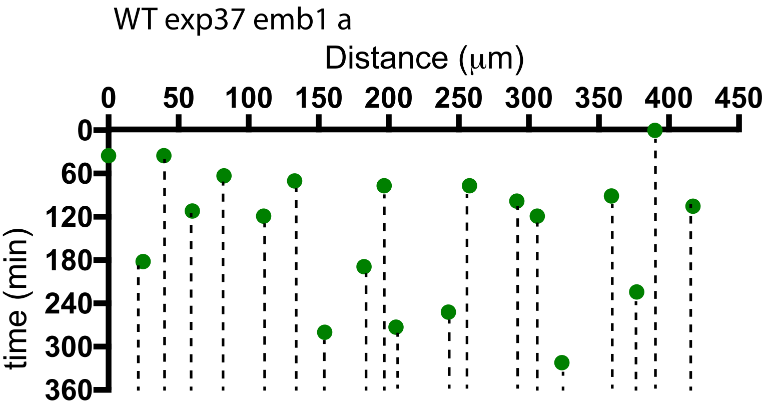
WT exp37 emb1 a
Distance (μm)
0
50
100
150
200
250
300
350
400
450
time (min)
0
60
120
180
240
300
360

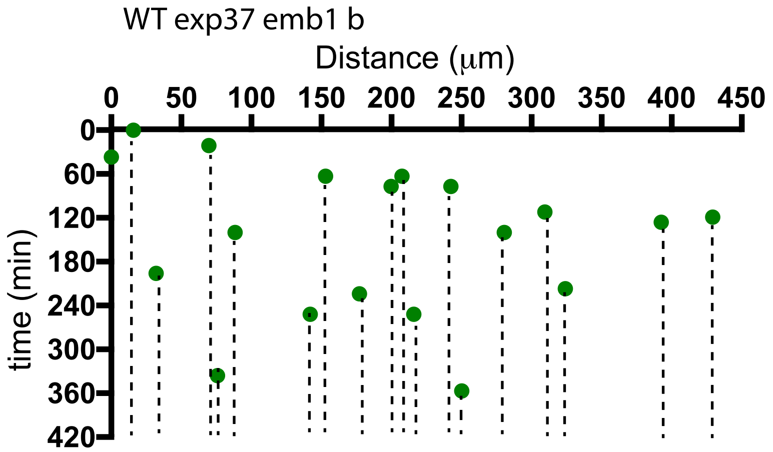
WT exp37 emb1 b
Distance (μm)
0
50
100
150
200
250
300
350
400
450
time (min)
0
60
120
180
240
300
360
420

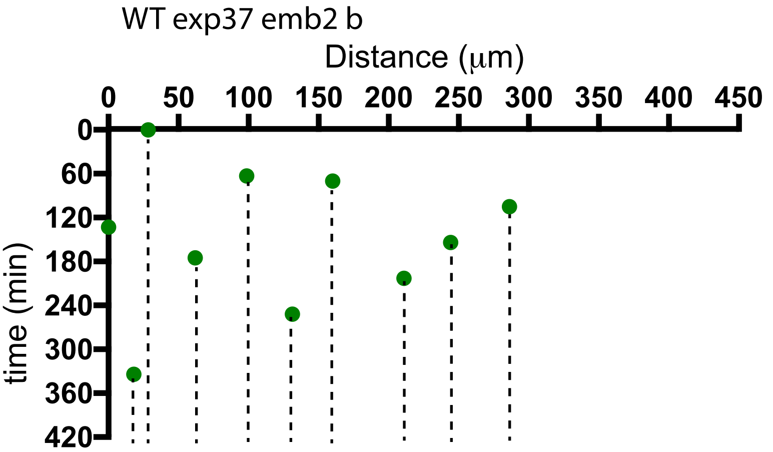
WT exp37 emb2 b
Distance (μm)
0
50
100
150
200
250
300
350
400
450
time (min)
0
60
120
180
240
300
360
420

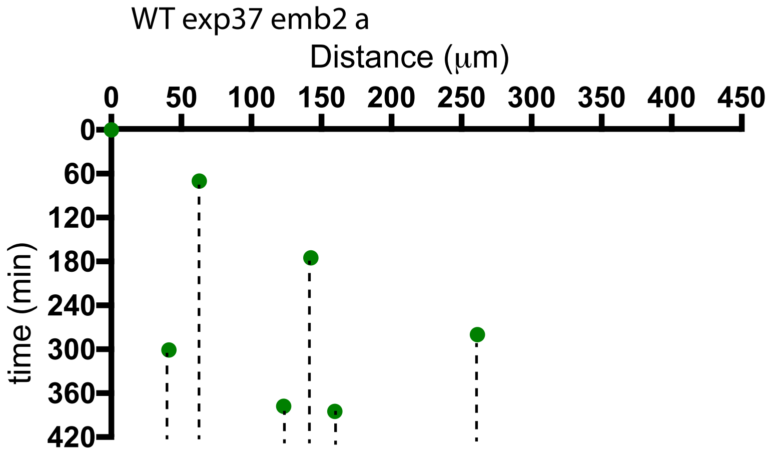
WT exp37 emb2 a
Distance (μm)
0
50
100
150
200
250
300
350
400
450
time (min)
0
60
120
180
240
300
360
420

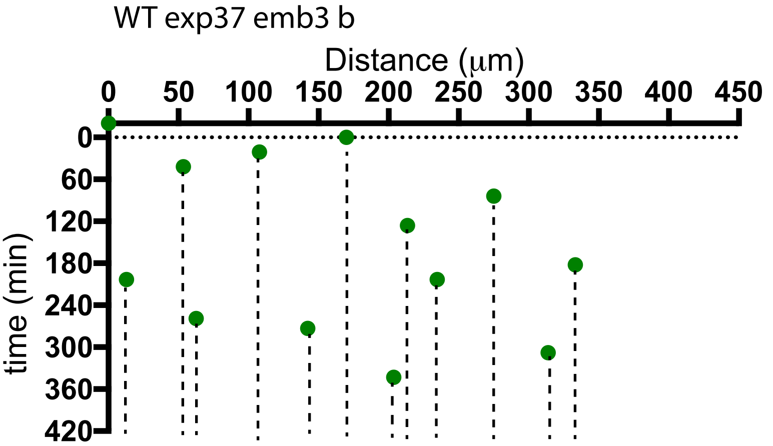
WT exp37 emb3 b
Distance (μm)
0
50
100
150
200
250
300
350
400
450
time (min)
0
60
120
180
240
300
360
420

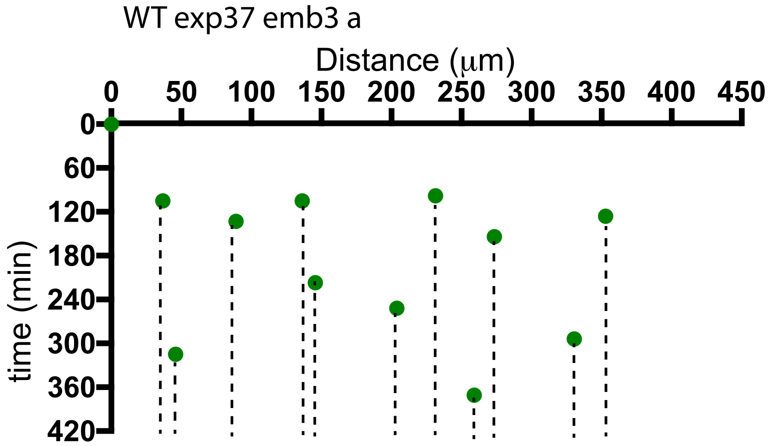
WT exp37 emb3 a
Distance (μm)
0
50
100
150
200
250
300
350
400
450
time (min)
0
60
120
180
240
300
360
420

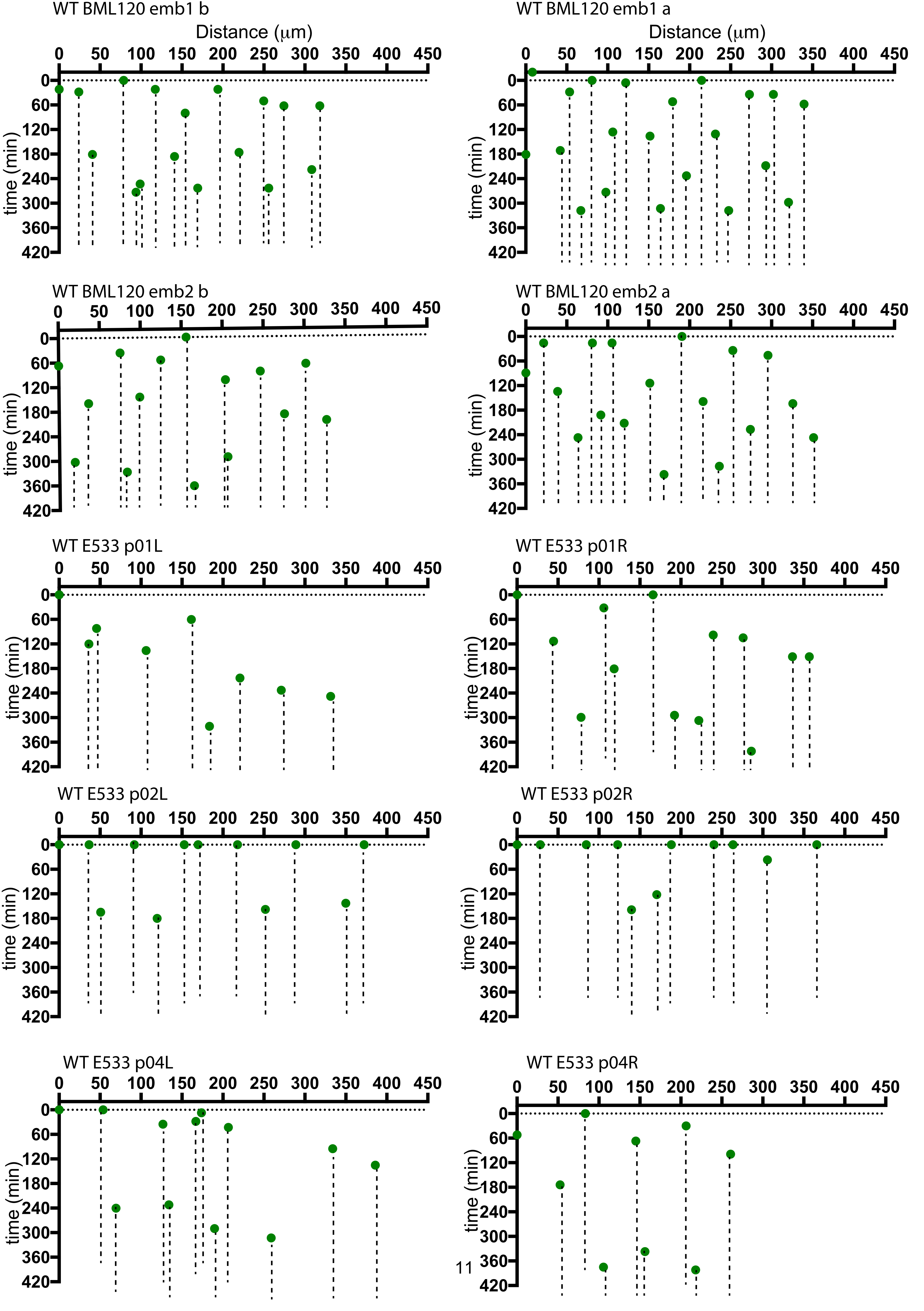
WT BML120 emb1 b
Distance (μm)
time (min)
WT BML120 emb1 a
Distance (μm)
time (min)
WT BML120 emb2 b
time (min)
WT BML120 emb2 a
time (min)
WT E533 p01L
time (min)
WT E533 p01R
time (min)
WT E533 p02L
time (min)
WT E533 p02R
time (min)
WT E533 p04L
time (min)
WT E533 p04R
time (min)

**Figure S5B**

### Spatiotemporal pattern diagrams illustrating Vsx1:GFP neuronal differentiation events in time and space for 50 stretches of developing spinal cord in *lamc1* mutants.

Lamc1 mut 5 R458 b

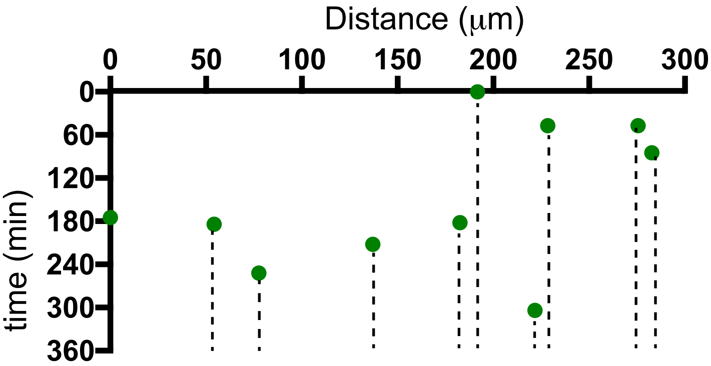

Lamc1 mut 5 R458 a

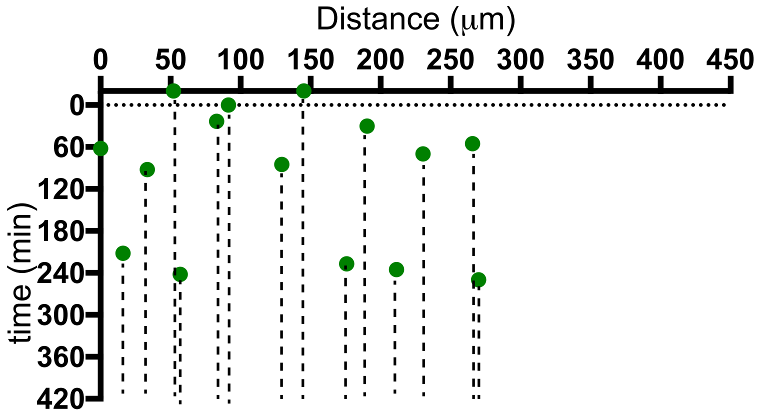

Lamc1 mut emb6 R458 b

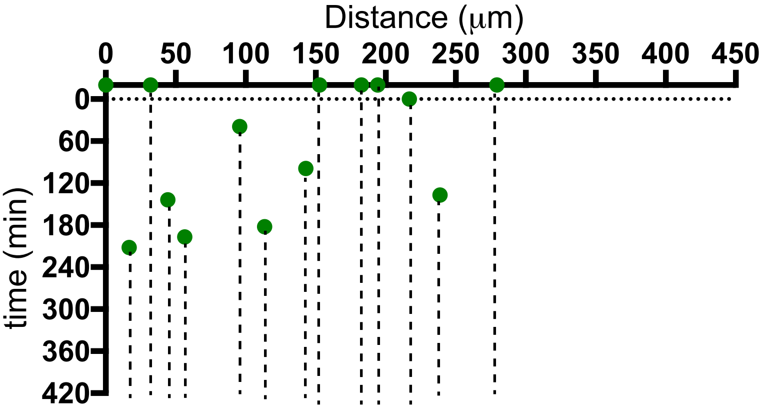

Lamc1 mut emb6 R458 a

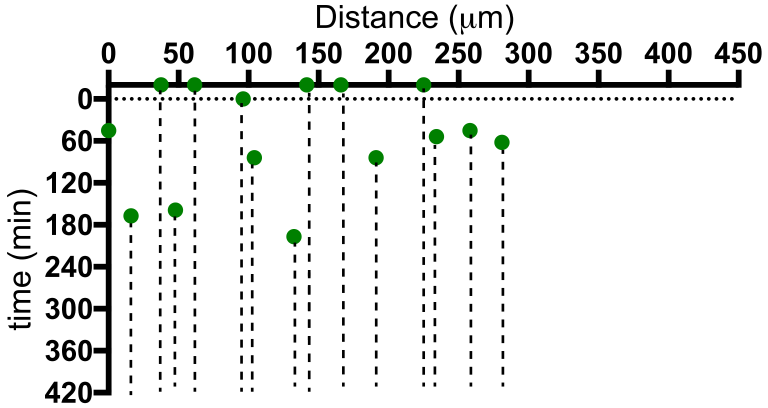

Lamc1 mut emb7 R458 b

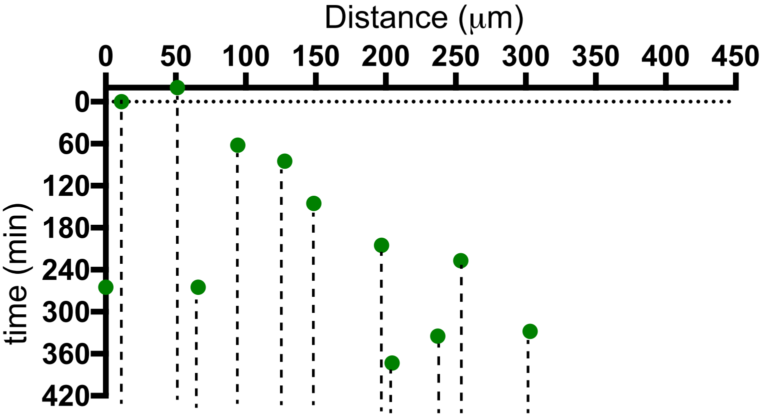

Lamc1 mut emb7 R458 a

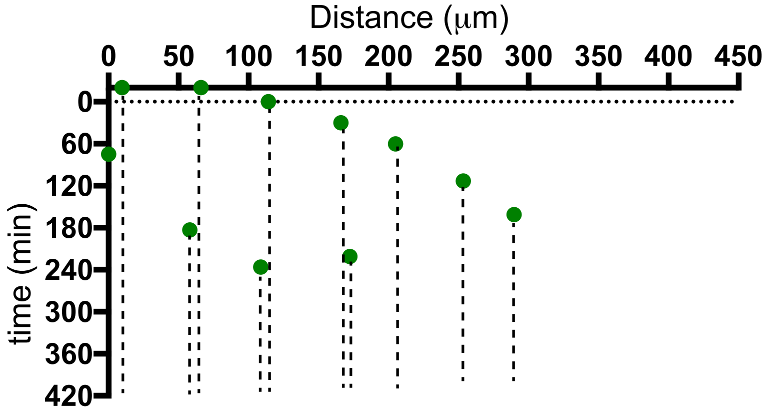

Lamc1 mut emb8 R458 b

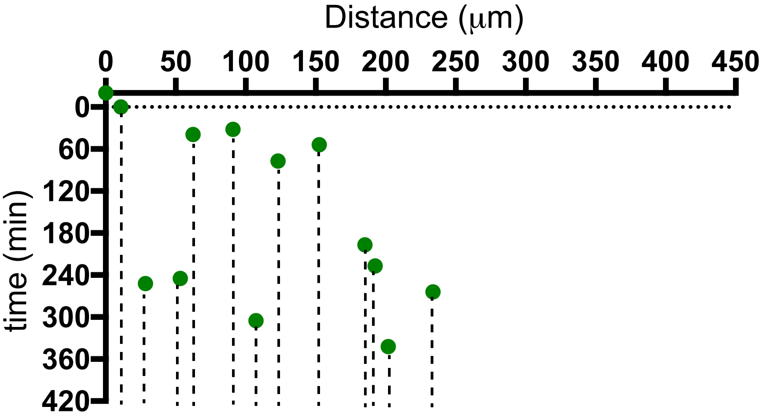

Lamc1 mut emb8 R458 a

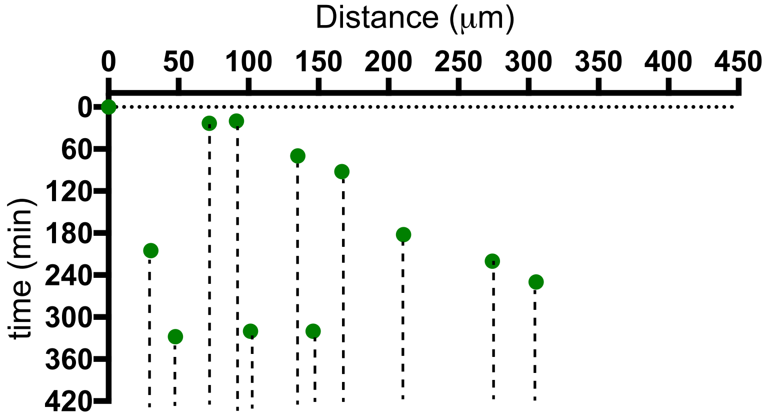

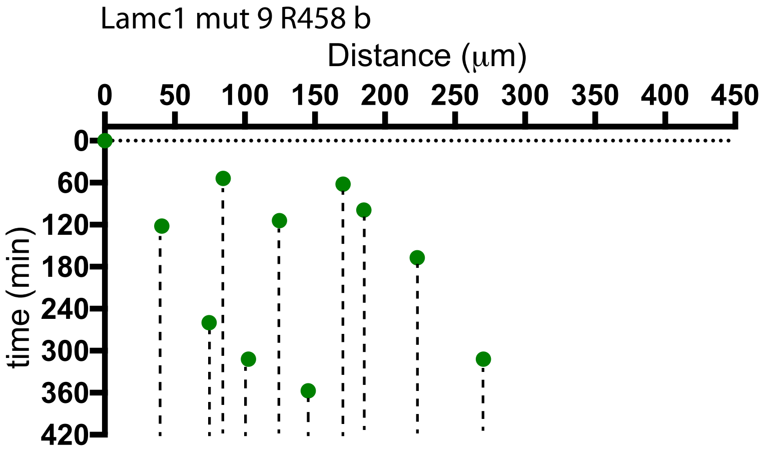
Lamc1 mut 9 R458 b
Distance (µm)
0 50 100 150 200 250 300 350 400 450
time (min)
0 60 120 180 240 300 360 420

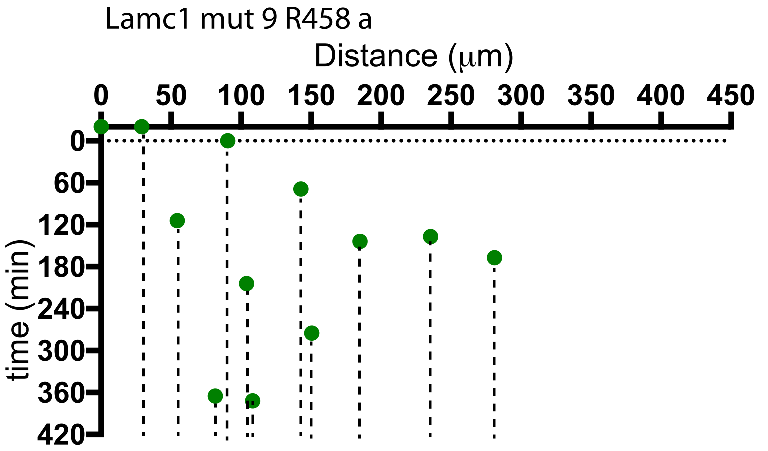
Lamc1 mut 9 R458 a
Distance (µm)
0 50 100 150 200 250 300 350 400 450
time (min)
0 60 120 180 240 300 360 420

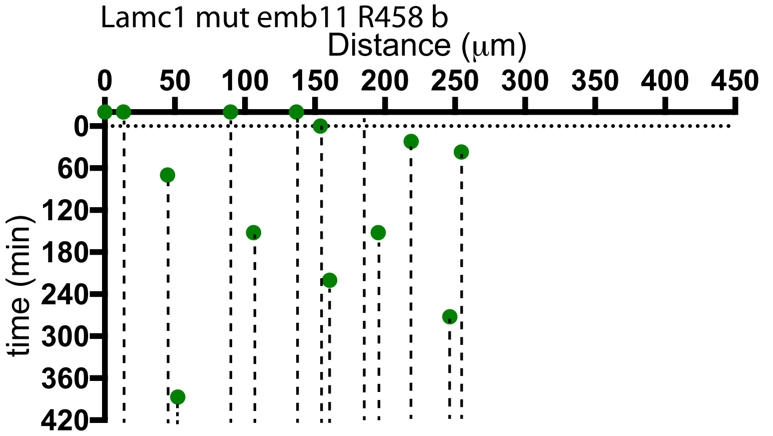
Lamc1 mut emb11 R458 b
Distance (µm)
0 50 100 150 200 250 300 350 400 450
time (min)
0 60 120 180 240 300 360 420

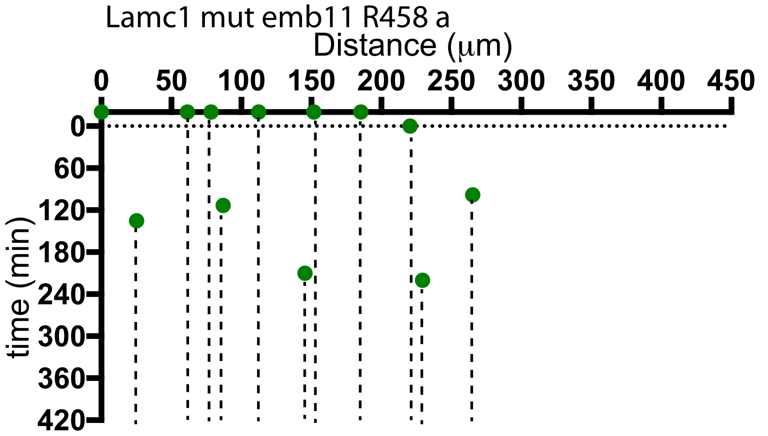
Lamc1 mut emb11 R458 a
Distance (µm)
0 50 100 150 200 250 300 350 400 450
time (min)
0 60 120 180 240 300 360 420

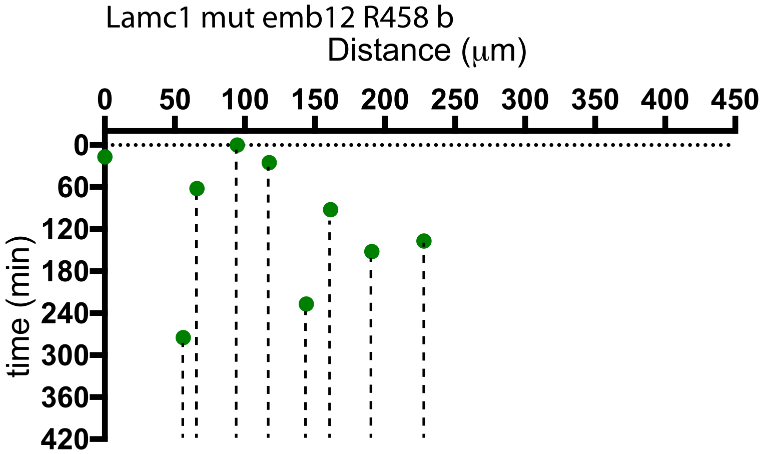
Lamc1 mut emb12 R458 b
Distance (µm)
0 50 100 150 200 250 300 350 400 450
time (min)
0 60 120 180 240 300 360 420

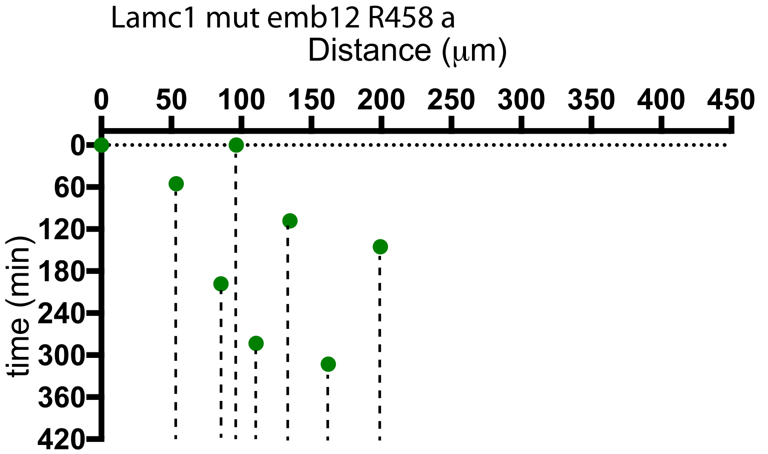
Lamc1 mut emb12 R458 a
Distance (µm)
0 50 100 150 200 250 300 350 400 450
time (min)
0 60 120 180 240 300 360 420

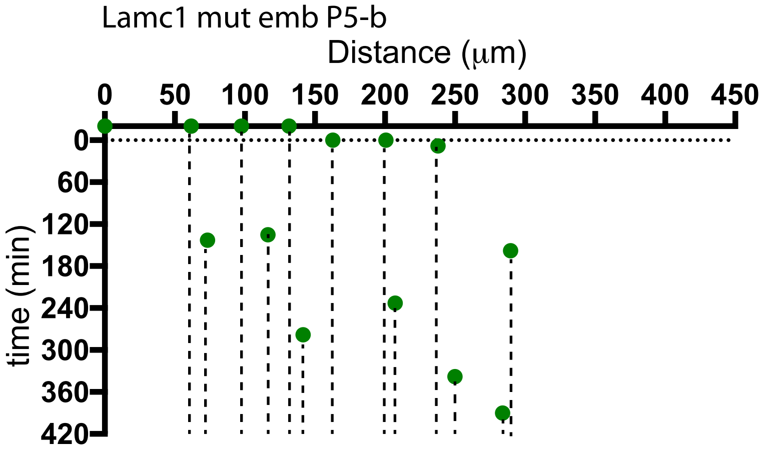
Lamc1 mut emb P5-b
Distance (µm)
0 50 100 150 200 250 300 350 400 450
time (min)
0 60 120 180 240 300 360 420

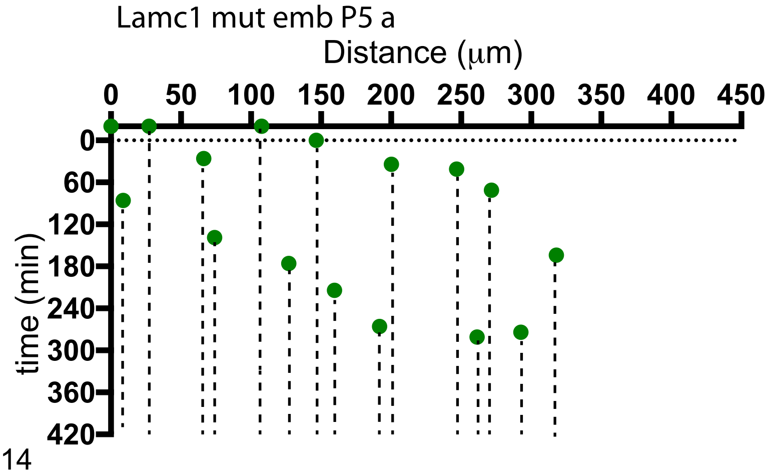
Lamc1 mut emb P5 a
Distance (µm)
0 50 100 150 200 250 300 350 400 450
time (min)
0 60 120 180 240 300 360 420

Lamc1 mut E461 P6 b

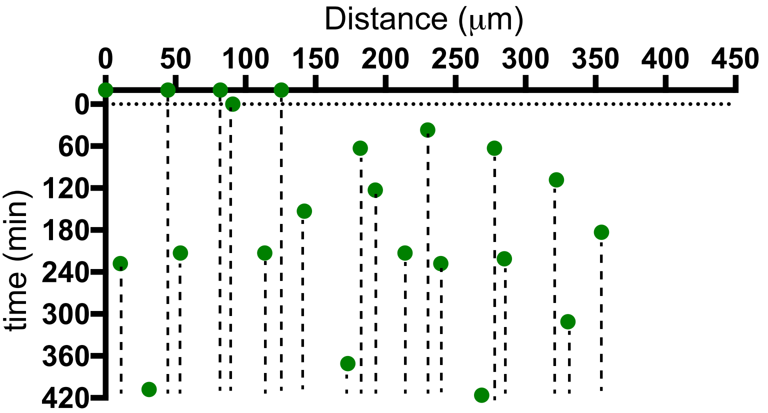

Lamc1 mut E461 P6 a

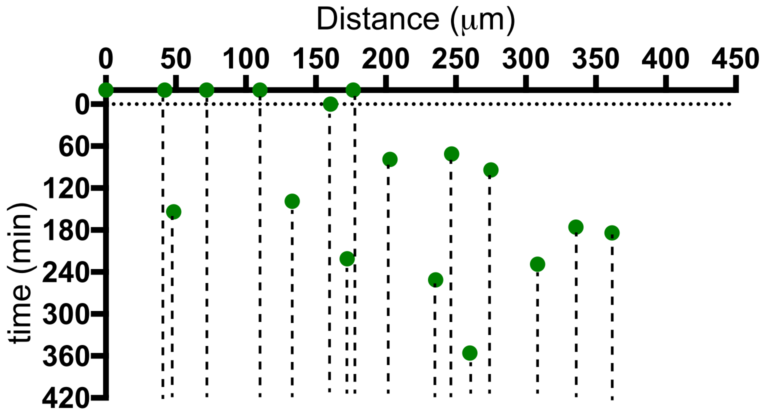

Lamc1 mut E461 P7 b

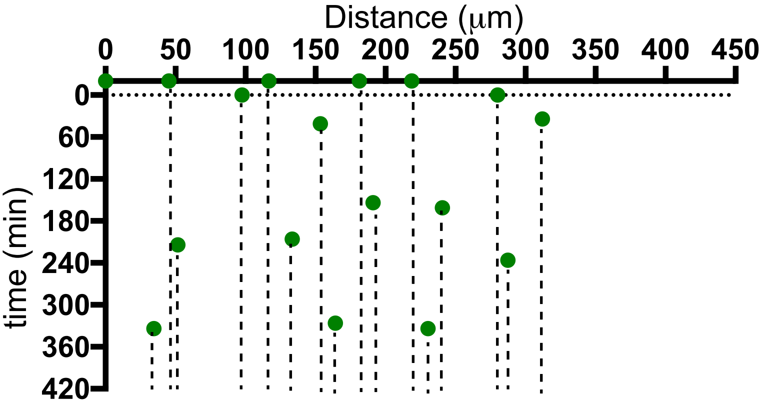

Lamc1 mut E461 P7 a

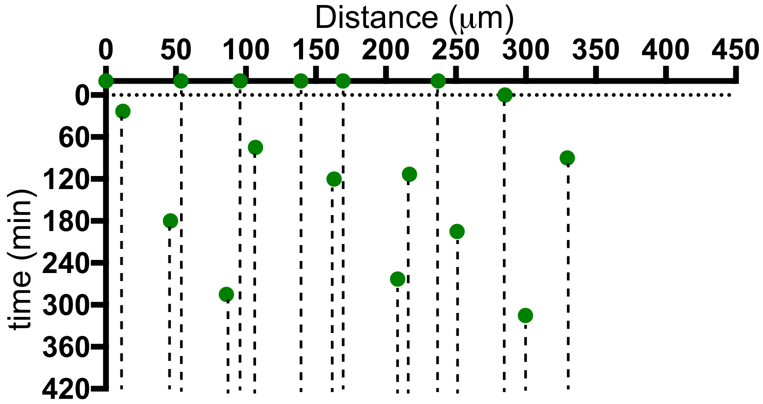

Lamc1 mut E461 P8 b

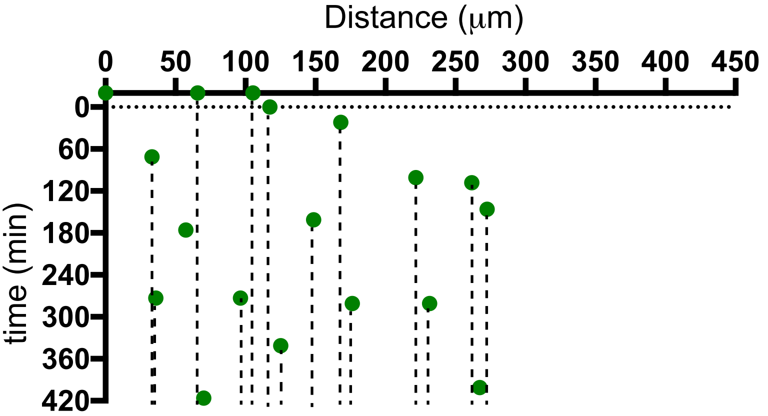

Lamc1 mut E461 P8

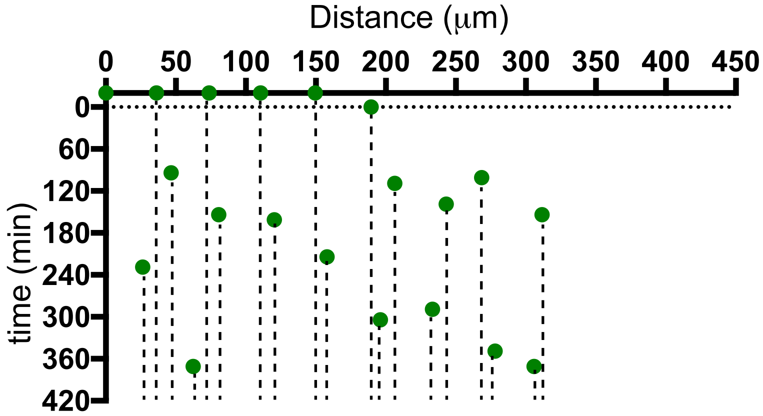

Lamc1 mut E461 P10 b

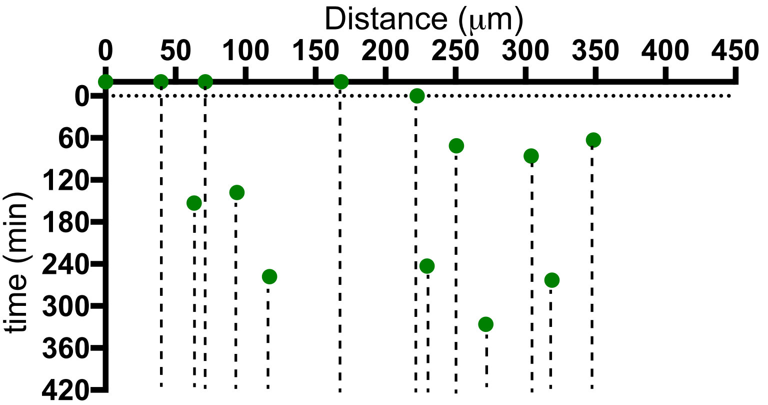

Lamc1 mut E461 P10

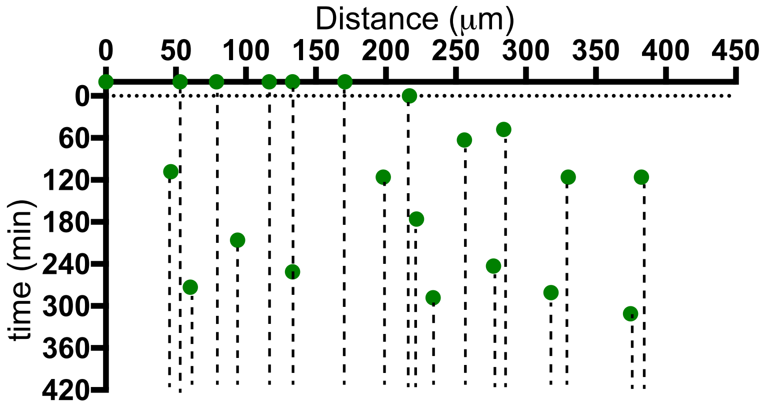

Lamc1 mut E461 P11 b

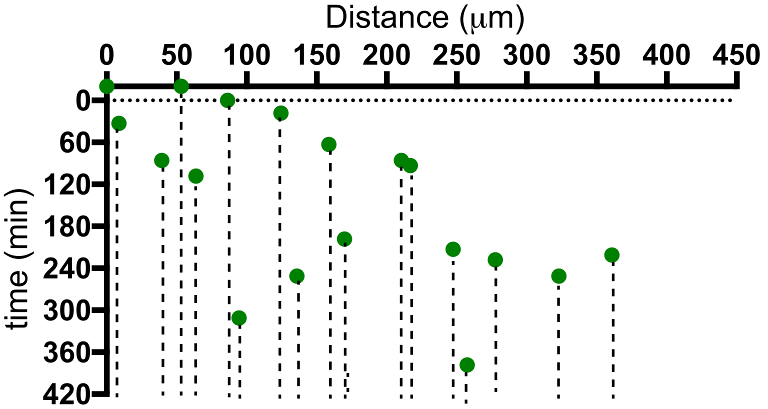

Lamc1 mut E461 P11 a

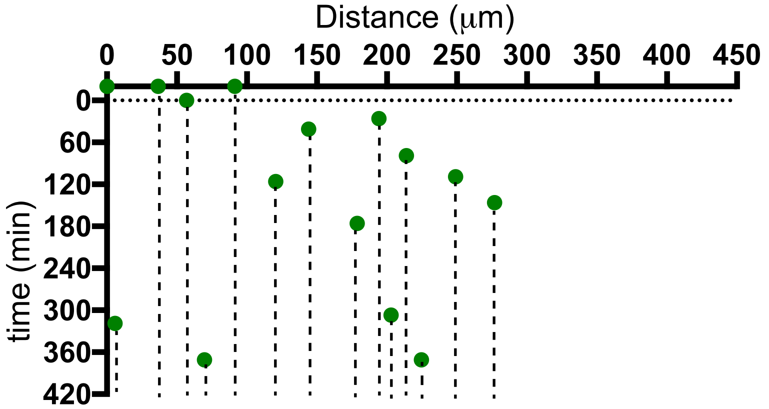

Lamc1 mut E461 P12 b

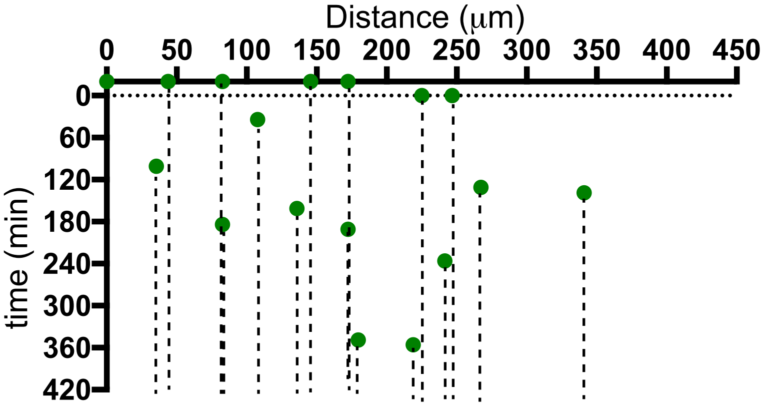

Lamc1 mut E461 P12 a

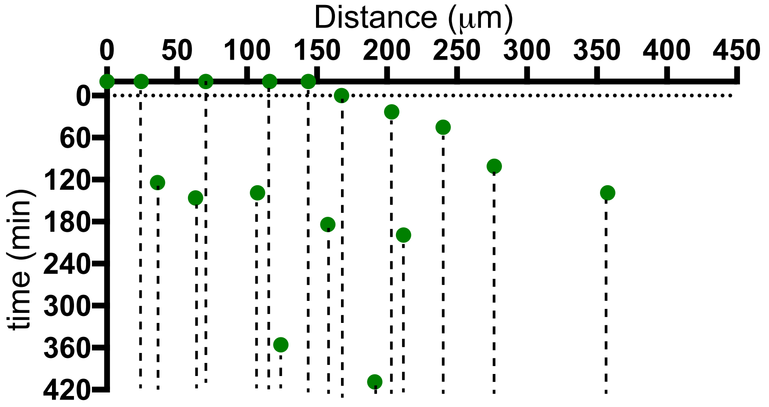

Lamc1 mut E466 emb1 b

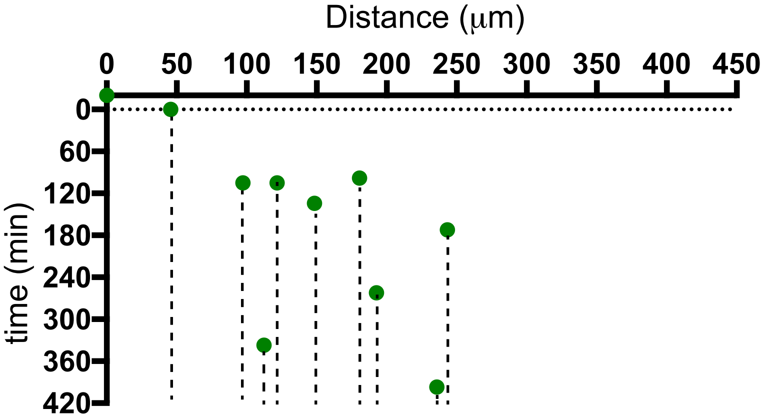

Lamc1 mut E466 emb1 a

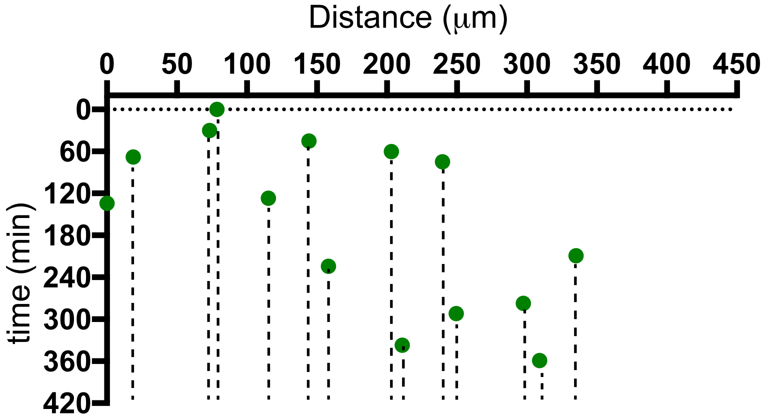

Lamc1 mut E466 emb2 b

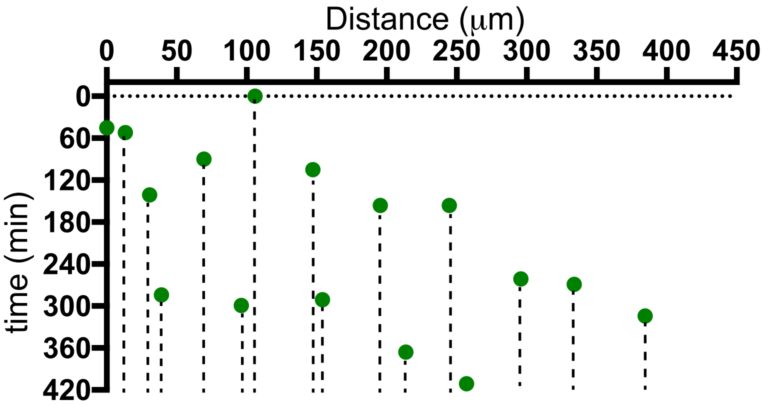

Lamc1 mut E466 emb2 b

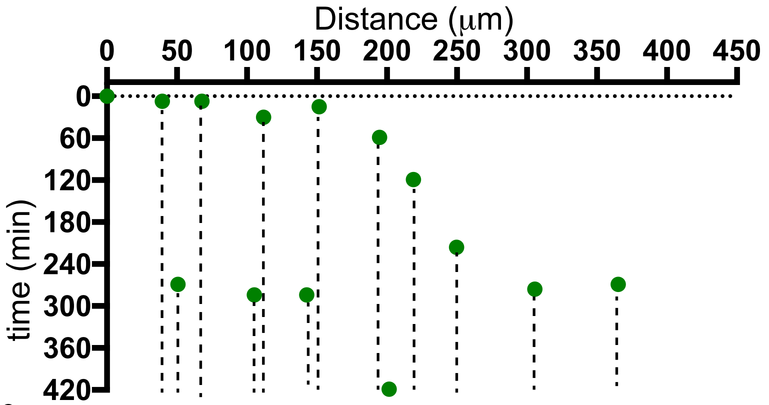

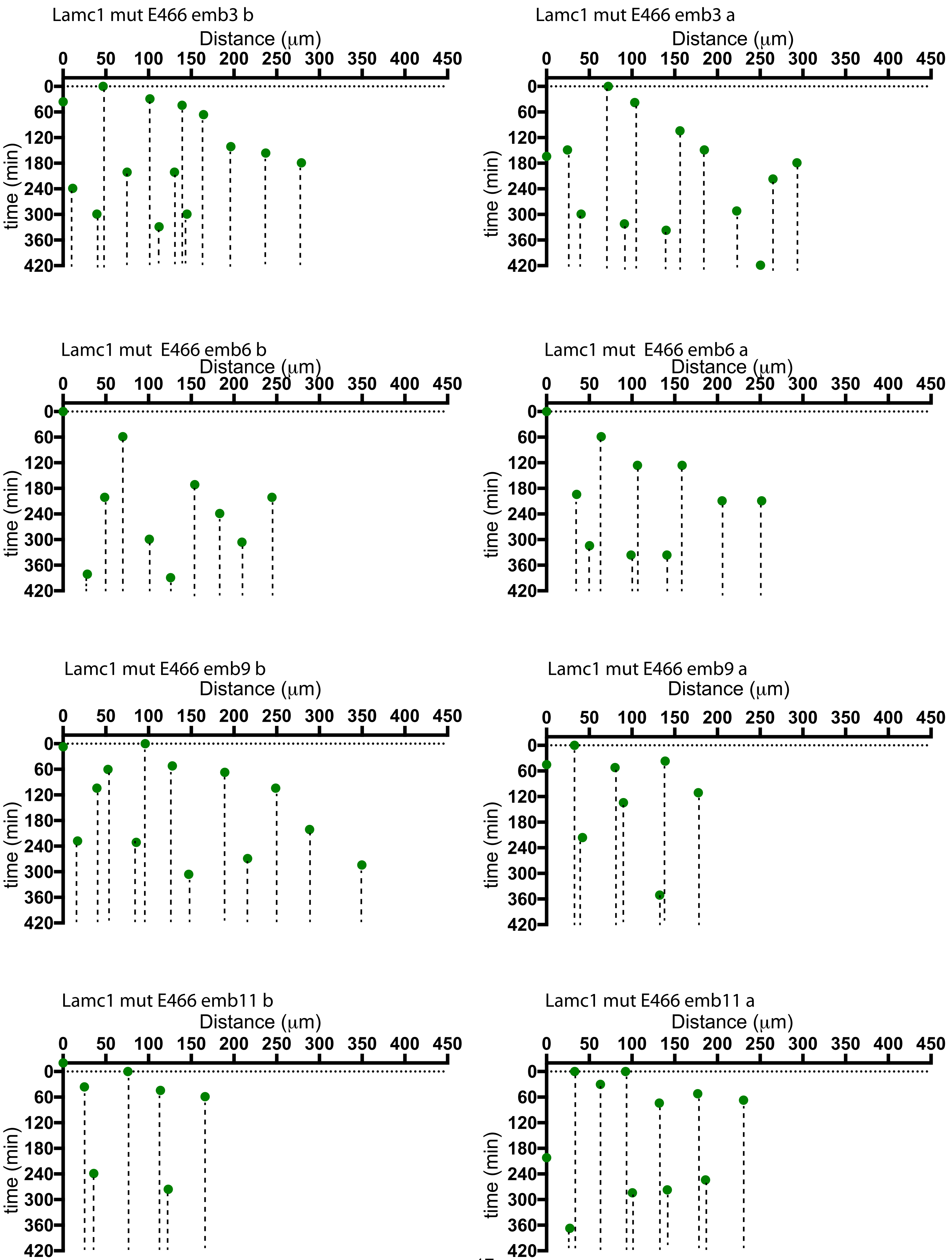
Lamc1 mut E466 emb3 b
Lamc1 mut E466 emb3 a
Lamc1 mut E466 emb6 b
Lamc1 mut E466 emb6 a
Lamc1 mut E466 emb9 b
Lamc1 mut E466 emb9 a
Lamc1 mut E466 emb11 b
Lamc1 mut E466 emb11 a
Distance (μm)
0 50 100 150 200 250 300 350 400 450
time (min)
0 60 120 180 240 300 360 420

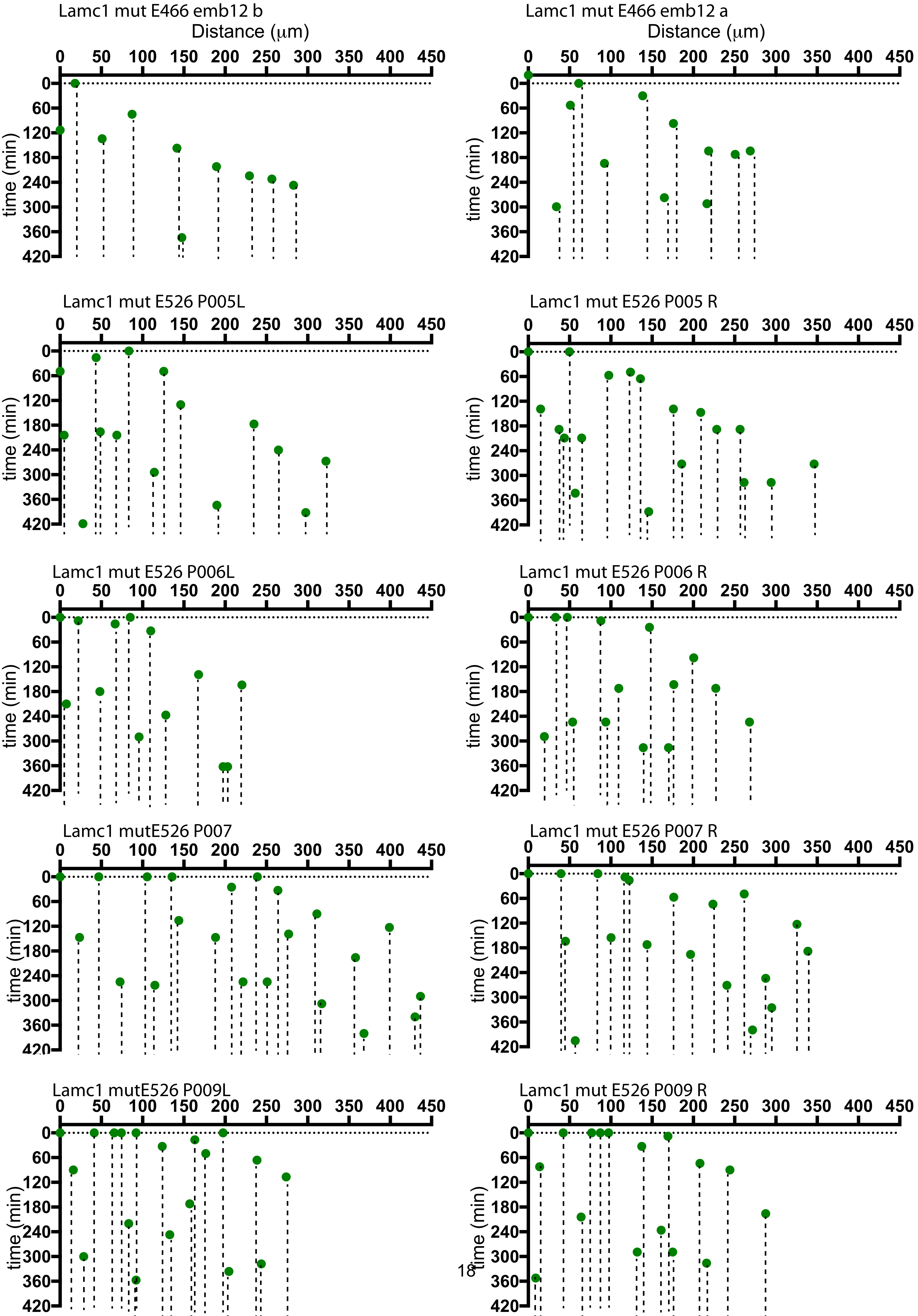
Lamc1 mut E466 emb12 b
Distance (μm)
time (min)
Lamc1 mut E466 emb12 a
Distance (μm)
time (min)
Lamc1 mut E526 P005L
time (min)
Lamc1 mut E526 P005 R
time (min)
Lamc1 mut E526 P006L
time (min)
Lamc1 mut E526 P006 R
time (min)
Lamc1 mutE526 P007
time (min)
Lamc1 mut E526 P007 R
time (min)
Lamc1 mutE526 P009L
time (min)
Lamc1 mut E526 P009 R
time (min)
0 50 100 150 200 250 300 350 400 450
0 60 120 180 240 300 360 420

### Supplementary Information 1

**3 Signaling at soma-to-soma contacts** 28

**4 Predicted changes in *dx* with variations in protrusion length** 29

**5 Sensitivity analysis** 30

**References** 34

### 1 Mathematical and computational details

We developed a theoretical description of lateral inhibition and cell differentiation in a one dimensional tissue (i.e. a row of cells). We construct the row of cells by sampling cell diameters from a normal distribution with mean 11.10$\mu$m and s.d. 4.51$\mu$m, the experimentally measured values in the neuroepithelium. This captures the diversity seen in the cell width of differentiating neurons, dividing cells and neuroepithelial cells. We used our set up to simulate differentiation events in the row of cells under different conditions as described below and in the main text. Here, we provide details of our numerical methods.

#### 1.1 Randomly differentiating tissue

We simulated a randomly differentiating tissue by initiating a row of cells as described above and then allowing cells to differentiate at random. In a row of $n$ cells this corresponds to sampling from $\{1, 2, ..., n\}$ without replacement and assuming that the $i^{th}$ sampled number is equivalent to the $i^{th}$ differentiation event. This allowed us to generate an ordered sequence of differentiation events and then compute the distance between cells (corresponding to the index numbers 1 to $n$) that were sampled successively (Figure S6). In this way we were able to predict the expected distance between successive events in a randomly differentiating tissue (Figure 8A).

#### 1.2 Lateral inhibition driven differentiation

We then wanted to make predictions about the distribution of the distance between successive events if differentiation events, $dx$, were regulated by Notch-Delta mediated lateral inhibition. We describe Notch-Delta signaling following previously published work (Collier et al., 1996; Cohen et al., 2010).

$$\frac{dN}{dt} = R_N \frac{D_{in}^k}{a + D_{in}^k} - \mu N \tag{1}$$

$$\frac{dD}{dt} = R_D \frac{1}{1 + bN^h} - \rho D \tag{2}$$

$$D_{in} = \alpha \sum_{\text{soma-to-soma}} D + \beta \sum_{\text{basal protrusions}} D \tag{3}$$

where $D_{in}$ is the total amount of incoming Delta summed over soma-to-soma and basal protrusion mediated contacts. The parameters $\alpha$ and $\beta$ represent the relative amount of Delta at the soma-to-

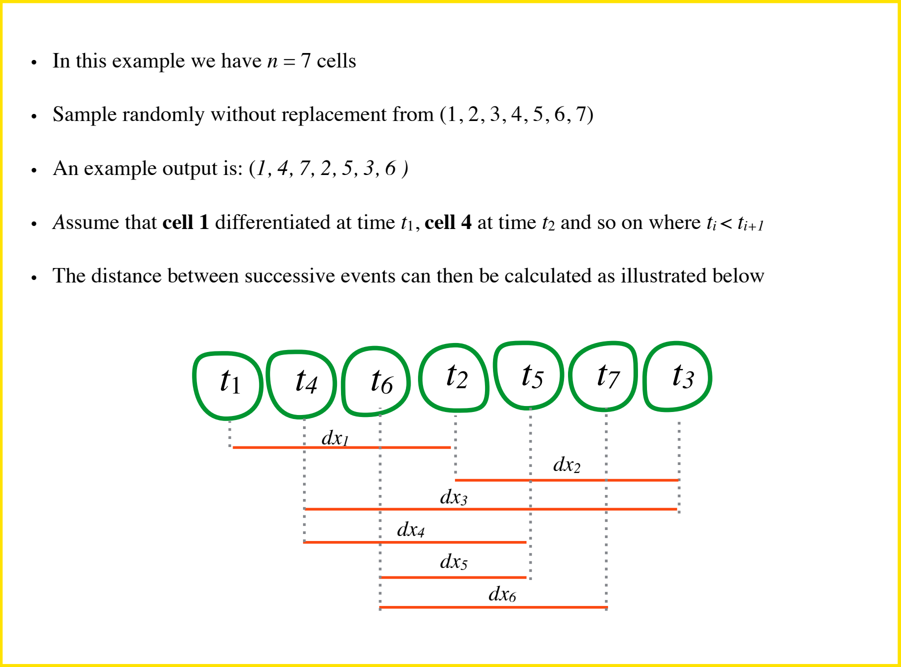

Figure S6: **Algorithm for the generation of a randomly differentiating spinal cord.** The positioning and size of cells were set according to experimental measurements.

soma and in the basal protrusions respectively or the strength of the signal at the two locations. In the analysis presented in the main text we assumed that $\alpha = 0$ so that only basal protrusions mediate Notch signalling. We also relaxed this assumption (see Section 2) to investigate whether Notch signalling at soma-to-soma contacts could also be important. $R_N$ and $R_D$ are the baseline production rates for Notch and Delta molecules, $a$ and $k$ are parameters that determine how strongly incoming Delta induces Notch signaling, whereas $b$ and $h$ determine the strength of inhibition of Delta from Notch levels within the same cell. Finally, $\mu$ and $\rho$ are the degradation rates of Notch and Delta.

We model protrusion dynamics following experimentally measured parameters so that the maximum length reached by basal protrusions follows a normal distribution with mean 42.6$\mu$m and standard error 4.7 $\mu$m. The extension rate is set to be 1.7 times larger than the retraction rate (as indicated by the measured data). We also assumed that mutant protrusion extension is slower than WT reflecting experimental measurements. We assume that cells begin extending their protrusions with a probability that depends on the levels of their Notch expression so that differentiation becomes more likely as Notch levels fall below a threshold. We implement this following previous work (Hunter et al., 2016) by

Table S1: **Parameters and definitions** Definition of parameters in mathematical model. The table also indicated the values used for all figures that use simulated data in the main text and supplementary material.

| **Variable** | **Definition** | |
|---|---|---|
| $N$ | Notch levels | |
| $D$ | Delta levels | |
| $D_{in}$ | Total incoming Delta | |
| $P_d$ | Arm extension and neuronal differentiation probability | |
| **Parameter (fixed)** | **Definition** | **Value** |
| $R_N$ | Baseline Notch production rate | 1.0 |
| $R_D$ | Baseline Delta production rate | 1.0 |
| $a$ | Dissociation constant in Notch production | 0.01 |
| $k$ | Hill coefficient in Notch production | 2 |
| $b$ | Dissociation coefficient in inhibition of Delta by Notch | 100 |
| $h$ | Hill coefficient in inhibition of Delta by Notch | 3.0 |
| $\mu$ | Notch degradation rate | 1.0 |
| $\rho$ | Delta degradation rate | 1.0 |
| $p$ | Maximal differentiation probability per unit time | 0.2 |
| $N_{th}$ | Notch threshold for differentiation | $10^{-4}$ |
| $q$ | Hill coefficient for differentiation probability | 1 |
| **Parameter (varied)** | **Definition** | |
| $\alpha$ | Delta signaling amount/strength at soma-to-soma contacts | 0 (Figure 8 , 0.1 (Figure S12)) |
| $\beta$ | Delta signaling amount/strength at basal protrusions | 0 (Figure 8A), 1 (Figure 8 B-F; Figure S11D-F, Figure S14) |
| $T_{ext}$ | Duration of basal protrusion extension period | 0.05 (Figure 8B; Figure S11D-F; Figure S12A), 0.07 (Figure 8C; Figure S11G- |

computing the probability of entering differentiation using a Hill function,

$$P_{diff} = p\frac{N_{th}^q}{N_{th}^q + N^q} \tag{4}$$

for each cell, where $N$ is the Notch expression of that cell and the parameters $N_{th}$ and $q$ determine a Notch threshold and the window around this threshold that lead to differentiation (Figure S7). The prefactor $p$ is the upper limit of the likelihood of differentiation per time step in the simulation. Differentiated cells no longer participate in lateral inhibition. In addition, protrusions are high in Delta but are assumed to carry a negligible number of free notch receptors (e.g. due to cis-inhibition) and so they only send but do not receive a signal (Sprinzak et al., 2010, 2011).

The values of all model parameters for all figures presented in the main and supplemental text are provided on Table 1.

##### 1.3 Simulation details

We initiate all simulations by randomly assigning each cell Notch and Delta levels sampled from $\mathcal{N}(R_N, 0.01R_N)$ and $\mathcal{N}(R_D, 0.01R_D)$ respectively where $\mathcal{N}(\mu, \sigma)$ denotes the Normal distribution with mean $\mu$ and s.d. $\sigma$ for values of $R_N$ and $R_D$ given on Table 1. Following this the Notch and Delta levels of each cell evolve according to Eq. (1-3) which we solved numerically using the Euler method (Euler step set to 0.01). At each step in the simulation each individual cell has a probability of initiating protrusion

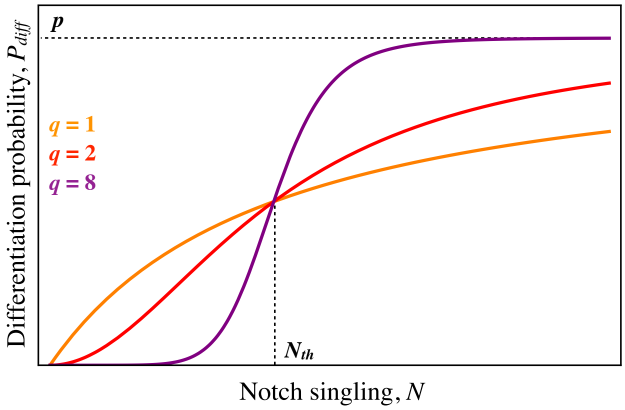

Figure S7: **Hill function determining the probability of cell differentiation as a function of Notch signaling.** The probability of differentiation is a Hill function of a cell's levels of Notch. The parameter $N_{th}$ determines the Notch signaling levels at which half-maximal probability of differentiation is reached. The Hill coefficient $q$ determines the sharpness of the transition from 0 to maximal probability of differentiation.

extension that is computed using Eq. (4). Cells that begin extending protrusions spend $T_{ext}$ a.u. of time extending their protrusions and $T_{ext}/1.7$ a.u. of time retracting their protrusions, reflecting the relative amount of time cells were experimentally observed spending in the protrusion extension and retraction stages respectively. Once full protrusion retraction is achieved a cell is assumed to have differentiated to a neuron and no longer participates in the process of lateral signaling. The simulation parameter $T_{ext}$ was set to 0.05 units of time for all WT simulations and 0.05*(mean length of experimental *lamc*1 mutant /experimental WT basal protrusions)*$dT$ units of time in short protrusion simulations where $dT = 1.4$ reflecting that lamc1 protrusions extended 1.4 times slower than WT basal protrusions. The retraction time in mutant protrusions in the simulations was set to 1.1 times their extension time, again reflecting experimental measurements. We further discuss the role of $T_{ext}$ in Section 5. A detailed outline of the algorithm we used throughout our analysis is shown on Figure S8.

Simulation code was written on c/c++. Simulated and experimental data were analyzed using scripts written on Wolfram Mathematica (Inc.).

##### 1.4 Comparison of simulated data to experiments

We performed simulations on a row of 50 cells (mean cell diameter = 11.10$\mu$m). The simulation was run until all cells differentiated. We repeated simulations for a given set of parameters 100 times to generate data for each experiment. We recorded the time and position of all events, computed the distance between sequential differentiation events and analyzed all sequential events that occurred within 80$\mu$m

```
Establish a line of n cells with apical radial sizes A, based on measured values (mean = 11.10, sd = 4.51)
for cells i=1:n
        A(i) = Random Normal(mean,  s.d.) // diameter of ith cell

pos(1) = 0 // position of first cell
for cells i=2:n
        x(i) = x(i-1) + A(i-1) + A(i)  // all other cells positioned relative to previous cell and according to their size

Assign each cell levels of Notch and Delta
for cells i=1:n
        N(i) = Random Normal(1.0,  0.01)
        D(i) = Random Normal(1.0 ,  0.01)

At each time step update Notch, Delta and arm distribution
For each time step
        For all cells i=1:n
                // Compute incoming delta from cell membrane contacts from immediate neighbours
                D_in_CM (i) = D(i+1) + D(i-1)

                // Compute incoming delta from basal protrusion contacts
                For all other cells j = i:n; j != i
                        If (i>j && DIF(j)==0 && (jth cell's right arm's length) >0 && (jth cell's right arm's length) >=
                        pos(i)-pos(j)) //DIF(j) = 0 if jth cell has not differentiated and 1 otherwise
                        OR
                        If (i<j && DIF(j)==0 && (jth cell's left arm's length) > 0 && (jth cell's left arm's length) >=
                pos(j)-pos(i))
                        Then
                        D_in_BP(i) = D_in_arms(i) + D(j)

                D_in_total(i) = alpha* D_in_CM(i) + beta* D_in_BP(i)

                // Update Notch levels
                Notch(i) = Notch(i) + (– µ.Notch(i) + R_N.D_in(i)k /(a+D_in_total(i)k ))*dt +
                RandomNormal(0,Notch(i)*rand_error) // random_error = 0.01 and dt = 0.05
                // Update Delta levels
                Delta(i) =  Delta(i) + (– ρ.Notch(i) + R_D 1 /(1 + b*Notch(i)h ))*dt
                + RandomNormal(0,Delta(n)*rand_error)

                // Update basal protrusions if time > 1.0 a.u.
                        If cell i has no basal protrusions, initiate extension with probability equal to:
                                value = 1.0 – p*N(i)^q/( N(i)^q + N_th(i)^q)

                        If basal protrusions extending, extent further by eurler_step*T_ext:
                                BP_left_L(i) = BP_left_L(i) + eurler_step*T_ext*Normal(F_mean, F_SE )
                                BP_right_L(i) = BP_right_L(i) + eurler_step*T_ext*Normal(F_mean, F_SE)
                                //F_mean = 42.3, F_SE = 3.09

                        If a basal protrusion has exceeded Normal(F_mean, F_SE ) initiate retraction.

                        If basal protrusion retracting, retract further by eurler_step*T_ext*1.7
                                BP_left_L(i) = BP_left_L(i) - eurler_step*T_ext*1.7* Normal(F_mean, F_mean*0.1 )

                                BP_right_L(i) = BP_right_L(i) - eurler_step*T_ext*1.7* Normal(F_mean,
                                F_mean*0.1 )

                        When retraction complete, cell differentiates and exits signalling

Return differentiation time and position for each cell
```

Figure S8: **Pseudo code.** Outline of the algorithm used to generate simulated data.

of one another. The spatial impact of the protrusions is only present at a length-scale that is relevant to the protrusion length. When our system is viewed at much larger length scales the spatiotemporal patterns we report become irrelevant. We therefore restricted our analysis to cells that differentiated within 80$\mu$m of one another, approximately two times the average protrusion length. The simulated data was then used to obtain the distribution of the distance between successively differentiating cells $dx$. We performed this analysis for randomly differentiating tissues and for tissues where differentiation was influenced by different models of lateral inhibition (basal protrusions only, soma-to-soma only, basal protrusions and soma-to-soma; Figure S9). The simulated data shown in the main text assume only basal protrusions mediate signaling. The role of soma-to-soma signaling is explored in a latter section.

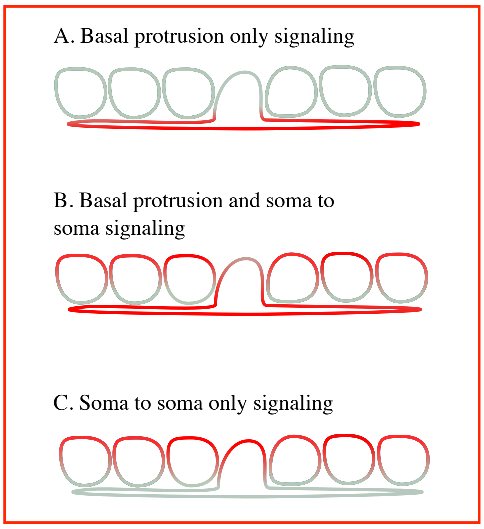

Figure S9: **Signaling models considered in theoretical set-up.** Red indicates the presence and gray the absence of signaling. A. Only basal protrusions can contribute to lateral signaling, B: basal protrusions and soma-to-soma contacts participate in lateral signaling, C: only soma-to-soma contacts contribute to lateral signaling.

#### 2 Pairwise differences in space and time

In order to futher investigate the coupling between the distance between any two differentiated cells and their time of differentiation we computed the distance in space, $\Delta x$ (not to be confused with $dx$ in the main text which is the distance between sequential differentiation events), and differentiation time, $\Delta t$, between all pairs of cells in each experiment and different versions of the theoretical model set-up. We then asked how the distributions of $\Delta x$ and $\Delta t$ depend on one another.

##### 2.1 Experimental data

The distribution of the pairwise position differences $\Delta x$ for all pairs follows an approximately uniform distribution on the measured interval (Figure S10A). When we restrict this distribution to cells that differentiate within one hour of each other, however, we observe a change in the distribution: very few cells differentiate within less than 30$\mu$m of one another and the distribution of $\Delta x$ becomes centered around 60$\mu$m (Figure S10B). Furthermore, $\Delta t$ and $\Delta x$ were negatively correlated (Spearmann's Rho = -0.26; Spearmann's Rank test p-value = 2.2 $10^{-12}$ )[1]. We also plotted the mean $\Delta t$ for cells that differentiated within a specific space interval of one another (Figure S10C). The smaller the distance between two cells the larger the difference in their time of differentiation. This effect appears to be present up until an interval of 50-60$\mu$m consistent with the average length of protrusions in the wild type.

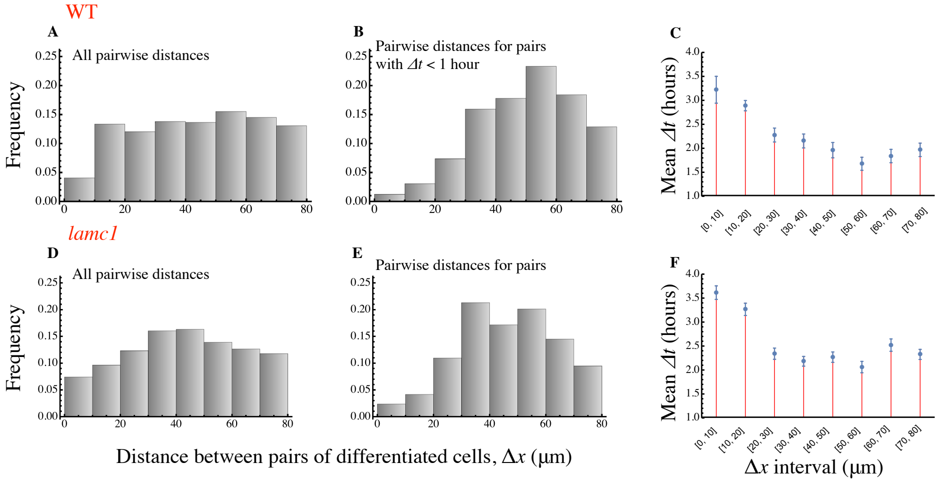

Figure S10: **Neurons that are born closer in time tend to differentiate further apart in space.** Distribution of pairwise distances between all differentiating neurons (A for experimental WT and D for *lamc*1 mutant) and neurons that differentiated less than 1 hour apart from one another (B for experimental WT and E for *lamc*1 mutant). C+F show the mean pairwise time difference in the time of differentiation ($\Delta t$) for varying intervals of pairwise spatial differences $\Delta x$ for the WT (C) and lamc1 mutant (F). All plots were produced using all measured pairs that satisfied $\Delta x < 80\mu$m.

We repeated the same analysis for the *lamc*1 mutant data. We once again found a negative correlation between $\Delta t$ and $\Delta x$ (Spearmann's Rho = -0.16; Spearmann's rank test p-value = 4.6 $10^{-10}$). Furthermore, the distribution of $\Delta x$ shifts with very few cells differentiating right next to each other. However, the distribution of $\Delta x$ conditional on $\Delta t <$ 1hour is shifted to the left in the *lamc*1 data when compared to the WT experimental data (Fig. S10B versus E). This is consistent with shorter basal protrusions governing the spatiotemporal dynamics in the *lamc*1 mutant. We again plotted the mean $\Delta t$ for cells that differentiated within a specific space interval of one another (Figure S10F). The smaller the distance between two cells the larger the difference in their time of differentiation. Unlike the wild type data, this effect is only present up until an interval of 20-30$\mu$m, consistent with a reduced range in

[1]The Spearmann's Rho and test measures a negative relationship between two variables that needs not be linear. Small p-values indicate a significant result, i.e. an interdependence between the tested variables.

lateral inhibition as reflected by the reduction in the protrusion length.

#### 2.2 Theoretical predictions

We then asked whether these observations are consistent with a randomly differentiating tissue or a tissue where basal protrusions mediate lateral inhibition. In a randomly differentiating tissue (where basal protrusions extend but do not signal) we get no correlation between $\Delta t$ and $\Delta x$ (Spearmann's Rho = 0.00185; Spearmann's Rank test p-value = 0.502 ), and conditioning the distribution of $\Delta x$ on $\Delta t$ has no impact (Figure S11A-C).

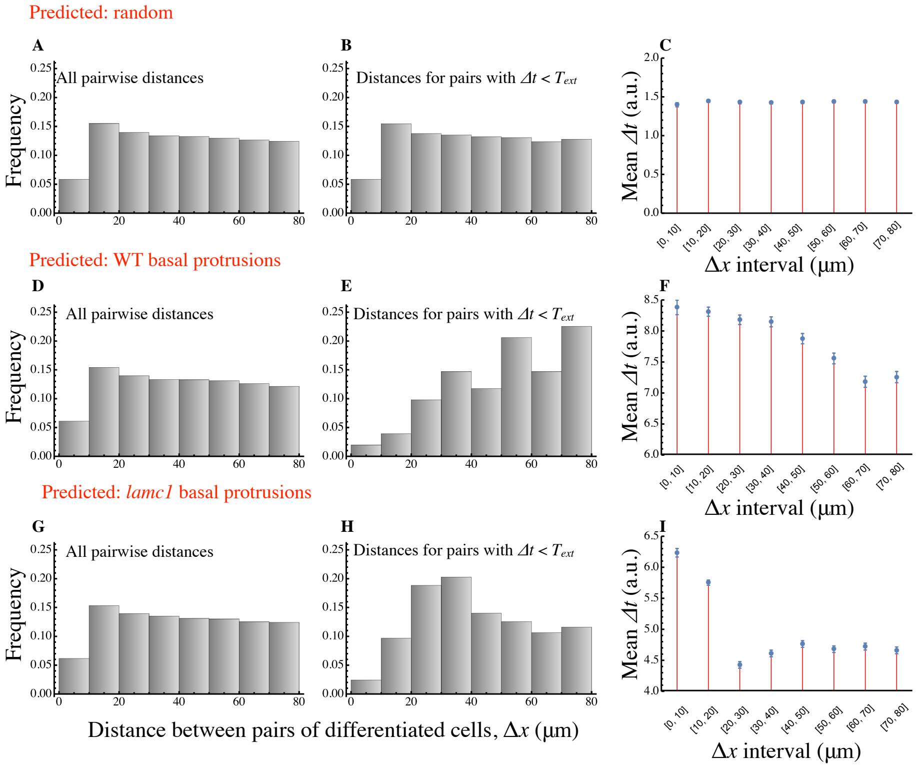

Figure S11: **Theoretical model supports the role of basal protrusions in patterning spatiotemporal dynamics of neuron differentiation.** Distribution of pairwise distances between all differentiating neurons (A. for random differentiation, D. for WT basal protrusions and G. for short basal protrusions) and neurons that differentiated less than $T_{ext}$ time apart from one another in the simulations (B. for random differentiation, E. for WT basal protrusions and H. for short basal protrusions). C, F+I show the mean pairwise time difference in the time of differentiation ($\Delta t$) for varying intervals of pairwise spatial differences $\Delta x$ for a randomly differentiating tissue (C), a tissue where WT basal protrusions mediate lateral inhibition (F) and a tissue where lamc1 like protrusions mediate lateral inhibition (I). Plots were produced using all simulated pairs that satisfied $\Delta x < 80\mu$m. Parameters used for the simulations are outlined in Table 1.

On the other hand, simulations where differentiating cells extend signaling protrusions of WT length lead to negatively correlated $\Delta t$ and $\Delta x$ (Spearmann's Rho = -0.10; Spearmann's Rank test p-value <

$10^{-20}$) and the distribution of $\Delta x$ shifts to the right when we condition on $\Delta t$ much like the experimental data (Figure S10A, B versus Figure S11D, E). Furthermore, when we plotted the mean $\Delta t$ for cells that differentiated within a specific space interval of one another we saw similar trends to those observed experimentally (Fig. S11F versus Figure S10C).

The same simulations but with short signaling basal protrusions also led to negatively correlated $\Delta t$ and $\Delta x$ (Spearmann's Rho = -0.12; Spearmann's Rank test p-value $< 10^{-20}$), but with a weaker distribution shift for $\Delta x$ when conditioning on $\Delta t$ and a reduced range for lateral inhibition as in the experimental data (Figure S10D-F versus Figure S11G-I). Taken together, these results further support our hypothesis that the spatiotemporal dynamics of neuronal differentiation is contingent upon lateral inhibition mediated by the long and transient basal protrusions we see *in vivo*.

### 3 Signaling at soma-to-soma contacts

Notch signaling typically occurs at soma-to-soma contacts between cells that are direct neighbors of one another (Lai, 2004). We therefore asked if soma-to-soma contacts could also play a role in our system. To investigate this we run simulations that incorporate lateral inhibition at some-to-soma contacts. This implies non-zero values for $\alpha$ in Eq. (3).

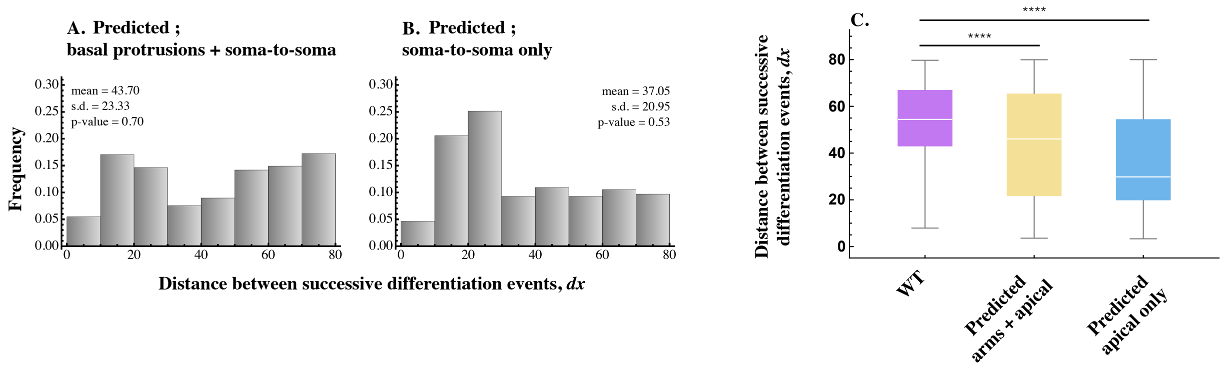

Figure S12: **Soma-to-soma signaling does not explain experimentally observed distributions.** Predicted distributions of the distance between sequential events when WT basal protrusions and soma-to-soma contacts mediate lateral inhibition (A) and when only soma-to-soma contacts mediate lateral inhibition (B). C shows the box plots of the WT data alongside the data shown in A and B. The simulated distributions considering soma-to-soma signaling do not match experimental observations.

When signaling that occurs at all cell contacts is included (i.e. protrusion to soma *and* soma-to-soma) the predicted distribution differs significantly from that observed experimentally (Figure S12A, C; Kolmogorov-Smirnov test, p-value $< 10^{-6}$). The predicted and observed distributions are even more different when we assume that lateral inhibition is only mediated at soma-to-soma membrane (and not basal protrusions) contacts (Figure S12B, C; Kolmogorov Smirnov test, p-value $< 10^{-10}$). In fact, in this latter case the predicted mean value for $dx$ is below that of a randomly differentiating tissue. This is because signaling taking place only at somal membrane contacts leads to differentiation events that occur in a typical checker-like pattern where cells that are one or two cell diameters apart tend to differentiate at a similar time (Collier et al, 1996, Hadjivasiliou et al 2016). This can be seen by the peaks at $dx = 30\ \mu$m in our histograms (Figure S12A, B). The absence of such a peak in our experimental data (Figure 7C in the main text) suggests that soma-to-soma contacts play a minimal if any role in the mechanism that determines the pattern of differentiation between spinal neurons.

### 4 Predicted changes in $dx$ with variations in protrusion length

In the main text we have shown that a specific change in the average maximum length reached by protrusions does not lead to the same change in the average distance between sequential events (Figure 8E). A change $dl$ in the protrusion length is only expected to lead to a change of $0.22dl$ in the average value of $dx$. This can be understood as follows. Consider a single differentiating cell which extends

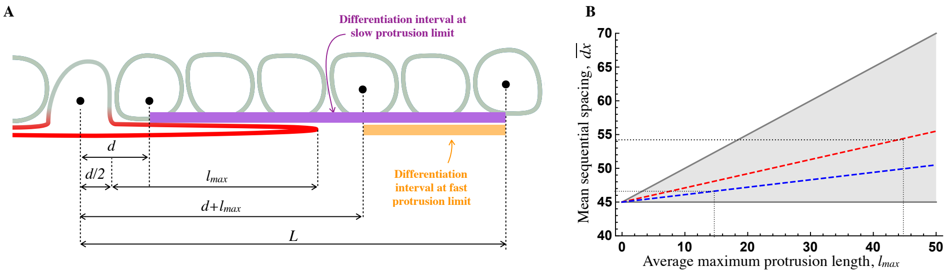

Figure S13: **The average distance between differentiation events depends linearly on the average maximum length reached by protrusions.** A: The region where the next differentiation event is expected to occur following protrusion extension. Two limiting cases are illustrated: one where protrusion extension is slow and inefficient and so the subsequent differentiation event is expected to occur anywhere between distance $d$ and $L$ from our cell of interest. The second case illustrated represents instantaneous protrusion extension and immediate later inhibition. In this case the subsequent differentiation event is expected to occur anywhere between distance $d/2 + l_{max}$ and $L$. B: The expected relationship between the average distance between sequential events $\bar{dx}$ and the average maximum protrusion length $l_{max}$ in the two limiting cases described in A (gray lines) and the expected curves in the WT (red) and lamc1 mutant (blue).

a protrusion of length $l_{max}$ and inhibits any cell within a distance $l_{max}$ from differentiating while the protrusion is present as shown in the diagram in Figure S13A. In the limiting case where the protrusion extends instantaneously the following differentiation event will occur at a distance between $d + l_{max}$

and $L$ from the differentiating cell with equal probability, where $L$ is the maximum distance away from our cell of interest (Figure S13A). It follows that the next differentiation event is expected to occur (on average) at a distance of $\frac{d + L + l_{max}}{2}$ away from our cell of interest (the mean of a Uniform distribution on the interval $(d + l_{max}, L)$). If we substitute $L = 80\mu$m (the maximum $dx$ value in our analysis) and $d = 10\mu$m (the average cell diameter) we obtain, $dx = 45 + 0.5\ l_{max}$. Assuming that the distribution of sequential differentiation events is stationary (i.e. time independent) it follows that,

$$\bar{dx} = 45 + 0.5\ \bar{l}_{max} \tag{5}$$

This means that in the limiting case where protrusions extend extremely fast a change in the average protrusion length equal to $dl$ will lead to a change in $\bar{dx}$ equal to only $0.5dl$.

Now consider a second case where the protrusions extend extremely slowly so they do not effectively inhibit neighboring cells from differentiating. In this case, neighboring cells will differentiate anywhere between $d$ and $L$ away from the differentiating cell and the expected value for $dx$ become independent of the protrusions so that,

$$\bar{dx} = 45 \tag{6}$$

We expect a real tissue to lie in between these two limiting cases so that,

$$\bar{dx} = 45 + \phi\ \bar{l}_{max} \tag{7}$$

where $\phi$ is a constant between 0 and 0.5 and depends on the timescale of protrusion extension and lateral inhibition relative to the timescale of differentiation (shaded region in Figure S13B).

We can compute $\phi$ for our experimental data by substituting $\bar{dx} = 54.0\mu$m and $\bar{l}_{max} = 42.6\mu$m for the WT and $\bar{dx} = 45.3\mu$m and $\bar{l}_{max} = 12.3\mu$m for the $lamc1$ mutant. It follows that $\phi_{WT} = 0.22$ and $\phi_{lamc1} = 0.024$. The decrease in the slope in the mutant is consistent with a reduced speed in protrusion extension, as observed experimentally.

#### 5 Sensitivity analysis

In this section we discuss the sensitivity of our simulated data and conclusions to variations in key parameters. We specifically explore the sensitivity of our conclusions to variations in parameters that determine the coupling between the basal protrusion growth dynamics and lateral inhibition. Parameters that determine feedback between Notch and Delta signaling have been explored in previous studies and we base our analysis the on these published works (Collier et al., 1996; Cohen et al., 2010).

##### 5.1 Random simulations

The simulation of neuronal differentiation in a randomly differentiating tissue is fully independent of any parameters. Our predictions were obtained using a random sampling algorithm on tissues of similar size and structure to the experiments (see Section 1.4). Therefore, our conclusion that the spatiotemporal dynamics observed experimentally are unlikely to come from a randomly differentiating tissue (p-value $< 10^{-10}$ for WT data and $< 10^{-6}$ for *lamc*1 data) is independent of any model parameters.

##### 5.2 Basal protrusion signaling simulations

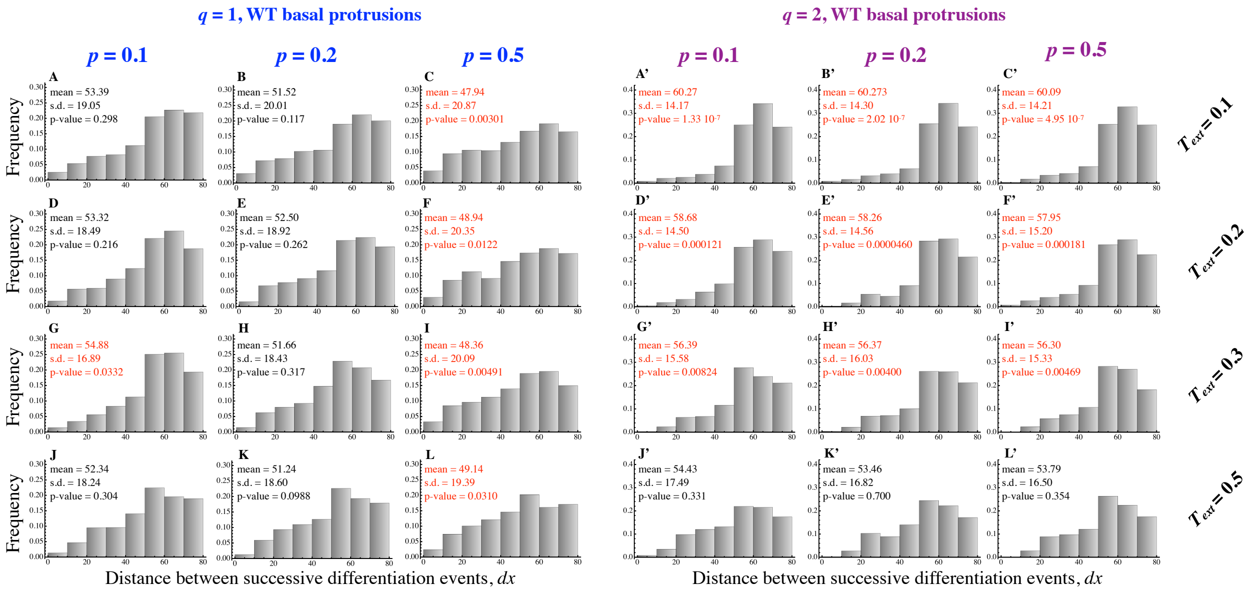

Figure S14: **Signaling simulations for various values of $p$, $q$ and $T_{ext}$ when lateral inhibition is mediated only through basal protrusions.** Histograms of the distances between successive differentiation events ($dx$) in the simulations. The mean and s.d. of $dx$ are shown together with the p-value when the distribution was compared to the WT experimental data. Numbers in red indicate a significant deviation from the WT experiments. Simulations were repeated 100 times and simulation parameters other than the ones varied in this analysis are given in Table 1.

To explore the dependency between protrusion and differentiation dynamics we varied three key parameters: the Hill exponent $q$ (see Fig. S7; Eq. (4)), the speed of the protrusion growth determined by the duration of the protrusions extension period $T_{ext}$, and the upper limit for the probability of differentiation per simulation step, $p$. For each variation we run simulations as described in Section

1.4 and compared the simulated distribution of the distance between sequential differentiation events between the simulations and WT experimental data as in the main text using the Kolmogorov-Smirnov test. Figure S14 shows the distributions for different parameters. We found that the comparison between simulation and experiment remains n.s. so long as protrusion dynamics and the cellular decision to differentiate are tuned together. When the baseline probability of differentiation is very high (Figure S14C, F,I, L) or the extending basal protrusions are too fast (Figure S14 A-C and A'-C') the simulated distribution diverges from the experiment. Very high probability to enter differentiation per time (higher $p$) leads to a reduction in the average $dx$. On the other hand, fast basal protrusions together with a high Hill coefficient lead to more narrow $dx$ distributions with larger average $dx$. Furthermore, higher values for $q$ also lead to more narrow $dx$ distributions for the same values of $p$ and $T_{ext}$ (Fig. S14A'-L'). However, key features of the simulated distribution remain robust to these variations. In particular, the peak near $dx = 60\mu m$ and the skewed distribution away from small values of $dx$ are seen in all our simulations. Hence, this analysis suggests that the exact behavior of the spatiotemporal dynamics depends on the coupling between the basal protrusion dynamics (actual speed of extension and retraction) and the initiation of cell differentiation as a response to levels of Notch signaling. The same should hold true in a real tissue: a weak dependency of differentiation on the basal protrusion dynamics and lateral inhibition would lead to weaker correlations in spatiotemporal dynamics of neuron differentiation.

##### 5.3 Basal protrusion and soma-to-soma signaling simulations

We repeated the analysis now allowing lateral inhibition to take place both through basal protrusions and at membrane-membrane contacts. With this combination of signalling, nearly all parameter combinations we tested give $dx$ distributions that deviate from the WT data (Figure S15). When we allowed very weak signaling at soma-to-soma contacts ($\alpha = 0.01$) some of our simulations were not significantly different from WT simulations. Such small values of $\alpha$ lead to membrane-to-membrane signaling is so weak it has a very minor impact on dynamics. These results suggest that soma-to-soma contacts may only contribute very weakly to lateral inhibition prior to protrusion extension.

##### 5.4 Soma-to-soma signaling simulations

When cells only signal at their soma contacts the basal protrusion dynamics do not matter. To explore whether soma-to-soma only signaling could lead to WT-like distributions we run simulations varying $p$ and $\alpha$. None of the simulated distributions were close to resembling the WT experimental data (Figure

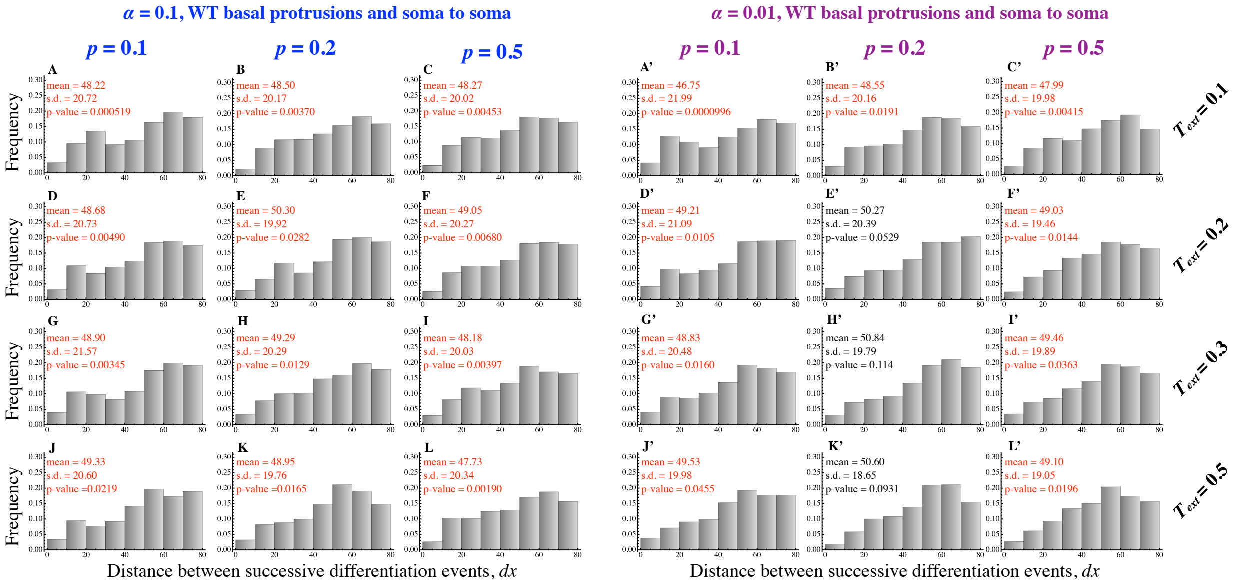

Figure S15: **Signaling simulations for various values of $p$, $\alpha$ and $T_{ext}$ when lateral inhibition is mediated only through basal protrusions and soma-to-soma contacts.** Histograms of the distances between successive differentiation events $(dx)$ in the simulations. The mean and s.d. of $dx$ are shown together with the p-value when the distribution was compared to the WT data. Numbers in red indicate a significant deviation from the WT experiments. Simulations were repeated 100 times and simulation parameters other than the ones varied in this analysis are given in Table 1.

S16). As in the main text, we find a bias for $dx$ between $20\mu$m to $30\mu$m, the typical distance between cells that are two to three membranes apart. Weaker soma signaling (reducing $\alpha$ to 0.01) led to distributions more similar to those of a randomly differentiating tissue. Therefore, our analysis suggests that the observed dynamics are unlikely to be due to lateral inhibition mediated at soma-to-soma contacts alone.

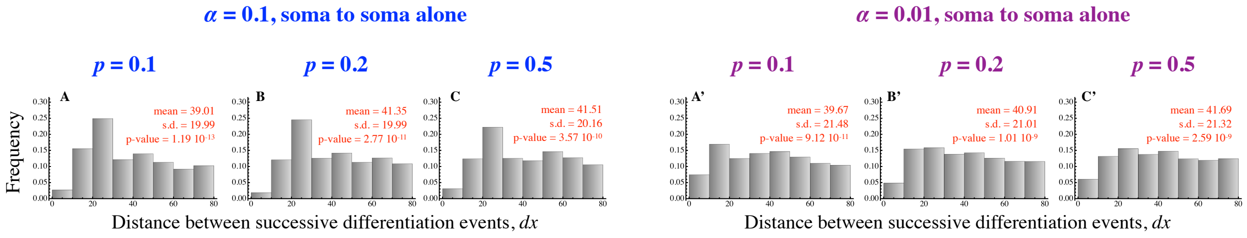

Figure S16: **Soma-to-soma only signaling simulations for varying $p$ and $\alpha$ when lateral inhibition takes place only at soma-to-soma contacts.** Histograms of the distances between successive differentiation events $(dx)$ in the simulations. The mean and s.d. of $dx$ are shown together with the p-value when the distribution was compared to the WT experimental data. Numbers in red indicate a significant deviation from the WT experiments. Simulations were repeated 100 times and simulation parameters other than the ones varied in this analysis are given in Table 1.
